# A kinetic operational model of agonism incorporating receptor desensitization for G-protein-coupled receptors

**DOI:** 10.1101/761726

**Authors:** Sam R.J. Hoare, David A. Hall, Lloyd J. Bridge

## Abstract

Pharmacological responses are modulated over time by regulation of signaling mechanisms. The canonical short-term regulation mechanisms are receptor desensitization and degradation of the response. Here for the first time a pharmacological model for measuring drug parameters is developed that incorporates short-term mechanisms of regulation of signaling. The model is formulated in a manner that enables measurement of drug parameters using familiar curve fitting methods. The efficacy parameter is *k_τ_*, which is simply the initial rate of signaling before it becomes limited by regulation mechanisms. The regulation parameters are rate constants, *k_DES_* for receptor desensitization and *k_D_* for response degradation. Efficacy and regulation are separate parameters, meaning these properties can be optimized independently of one another in drug discovery. The parameters can be applied to translate in vitro findings to in vivo efficacy in terms of the magnitude and duration of drug effect. When the time course data conform to certain shapes, for example the association exponential curve, a mechanism-agnostic approach can be applied to estimate agonist efficacy, without the need to know the underlying regulatory mechanisms. The model was verified by comparison with historical data and by fitting these data to estimate the model parameters. This new model for quantifying drug activity can be broadly applied to the short-term cell signaling assays used routinely in drug discovery and to aid their translation to in vivo efficacy, facilitating the development of new therapeutics.

**Highlights:**

- Regulation of signaling impacts measurement of drug effect
- Receptor desensitization is incorporated here into a kinetic model of signaling
- Drug effect and signaling regulation can now be measured independently
- The analysis framework is designed for signaling assays used in drug discovery
- These new analysis capabilities will aid development of new therapeutics

## 1. Introduction

Pharmacological theory has provided the concepts and analytical framework now used routinely in modern drug discovery and development (Rang, 2006). In lead optimization, potency (EC_50_) and maximal effect (E_max_) are used to establish structure-activity relationships and predict the in vivo effectiveness of new molecules. In pharmacological models, these empirical measures are converted into chemical drug parameters such as affinity and efficacy (Kenakin, 2009b). Most pharmacological theory and much drug development are based upon G-protein-coupled receptors (GPCRs). From bacteria to higher organisms these receptors have evolved to respond to a remarkable diversity of signals, including photons, ions, neuromodulators, hormones and enzymes (Fredriksson et al., 2003). They have proved to be highly tractable targets for small molecule therapeutics (Sriram and Insel, 2018).

GPCR signaling is regulated in order to prevent over-stimulation of cellular responses (Hausdorff et al., 1990; Krupnick and Benovic, 1998). These mechanisms act to attenuate signaling on persistent or repeated exposure of the receptor to the agonist. Regulation occurs at two levels in the signal transduction process, at the level of signal generation by the receptor (receptor desensitization), and at the level of signal degradation downstream of the receptor. In the receptor desensitization mechanism, agonist-bound receptor is phosphorylated on intracellular residues by kinase enzymes (Inglese et al., 1993; Stadel et al., 1983). The phosphorylated receptor then interacts with arrestin (Krupnick and Benovic, 1998; Lohse et al., 1990). Arrestin binding prevents access of G-protein to the receptor, blocking G-protein signaling. The signal, once generated, can be degraded by regulatory processes in the signaling pathway. These include metabolic processes that reduce the level of second messenger molecules such as cAMP and inositol phosphates (Berridge, 1993; Chang, 1968; Naccarato et al., 1974), and export mechanisms such as the efflux of calcium ions (Carafoli, 1991).

Regulation of signaling is not included in the pharmacological models used routinely to measure agonist activity in drug development. [Numerous models have been developed to simulate system behavior, revealing the manifestation of mechanisms in signaling data (Leff, 1986; Riccobene et al., 1999; Violin et al., 2008; Xin et al., 2008). These models are not routinely used to measure ligand activity.] Notably, there is no pharmacological parameter for regulation of signaling in common use. Incorporating regulation of signaling would be useful for a number of reasons. It could aid prediction of in vivo activity; agonists that differ in their regulation of signaling would be anticipated to differ in their in vivo activity, especially for prolonged responses (Hothersall et al., 2016) or on repeated dosing (Tay et al., 2018). In analyzing SAR, it could tease apart effects of chemical substitutions on signaling efficacy versus receptor desensitization. Currently, these processes are effectively combined in the response measurement and this can result in time-dependence of drug parameter estimates (Leff, 1986; Navratilova et al., 2007; Riccobene et al., 1999). This effect complicates the comparison between compounds and between signaling assays, especially in biased agonism assessment (Klein Herenbrink et al., 2016; Lane et al., 2017); in principle, differences at early time points can be largely eliminated at later time points.

A pharmacological model that incorporates regulation of signaling must incorporate time because regulation of signaling is dynamic. Regulation controls how the response evolves over time, for example fade (the response declines following a peak) and leveling off (the response reaches a plateau). Recently a dynamic pharmacological model for G-protein-coupled receptor signaling was presented for quantifying agonist efficacy in routine drug discovery pharmacology (Hoare et al., 2018). This model included one of the two regulation of signaling axes, degradation of the response. In the current study, the model is extended to incorporate receptor desensitization. This provides a model that can be broadly applied to the short-term cell signaling assays typically used in drug discovery. The aims off this study were:

1. To formulate the model
2. Determine its manifestation in the shape of time course data
3. Determine the drug parameters that can be measured by curve fitting
4. Develop a readily-adoptable data analysis framework using generic, familiar equations found in commercial curve-fitting software (e.g. GraphPad Prism, Sigmaplot, XLfit).
5. Measure drug parameters by applying the model to experimental data.

## 2. The model

### 2.1. Framework

Recently a model was developed to quantify the kinetics/dynamics of agonist action on G-protein-coupled receptor signaling pathways (Hoare et al., 2018). The model is based on the primary characteristic of GPCR signaling – intermolecular interaction between components in the signal transduction cascade (Gilman, 1987). For example, receptor interacts with G-protein, which then interacts with adenylyl cyclase, which catalyzes the formation of cAMP, which interacts with protein-kinase A, which, through a series of further intermolecular interactions, results in gene expression. This series of interactions is modeled by reducing it to a single global interaction between the receptor and the signaling system [the approach used in the operational model of agonism (Hoare et al., 2018)]. Specifically, the precursor of the response (*E_P_*) is transformed to the response (*E*) by the action of the receptor (*RA*) on the signal transduction system. This is represented in Fig. 1, in the green-shaded region. The precursor is termed here “Response precursor.” (In the original model this was termed “Transduction potential.” (Hoare et al., 2018)] While formulated on explicit mechanistic principles, the model can also be derived as an adaptation of general, less-mechanistic classical pharmacological models. Specifically, the model is an adaptation of the operational model of agonism (Black and Leff, 1983), in which the kinetic constant *k_E_* replaces the equilibrium constant *K_E_* (Hoare et al., 2018). Consequently, the kinetic model can be viewed as an adaptation of established pharmacological theory. The model is also analogous to classical enzyme kinetics, where *RA* is the enzyme, *E_P_* the substrate, and *E* the product. The difference between the model and enzyme kinetics is that in the former *RA* coupling to *E_P_* is sufficiently transient that the *RA-E_P_* complex does not deplete *RA* or *E_P_*, whereas in the latter the enzyme-substrate complex contributes appreciably to the total concentration of enzyme.

**Fig. 1.**
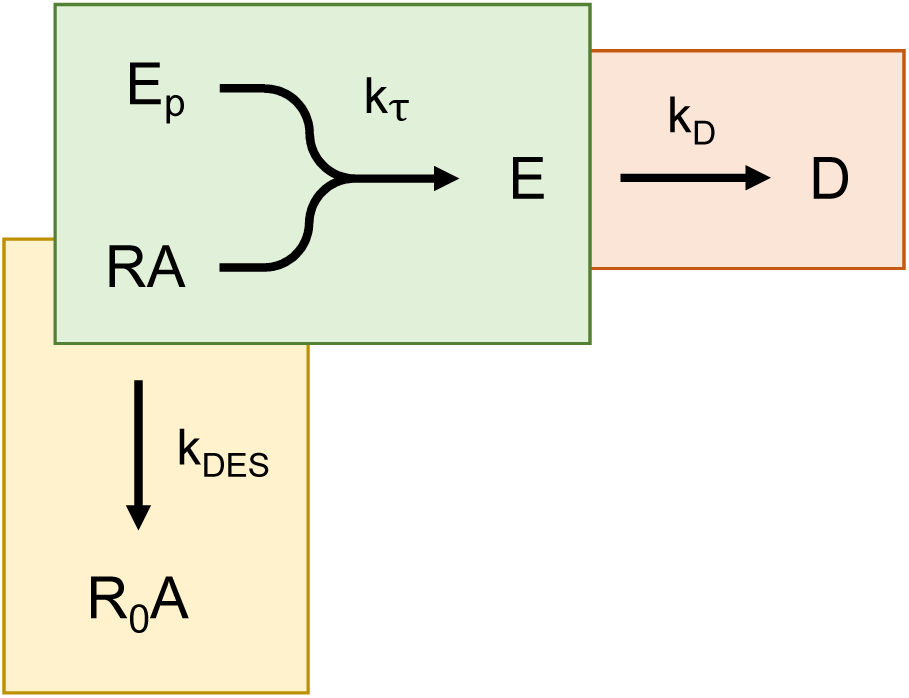
Kinetic operational model of agonism incorporating regulation of signaling. Both mechanisms of short-term signaling regulation are included – receptor desensitization (yellow) and response degradation (pink). The response generation process is in green. In this process, the response precursor (*E_P_*) is converted to the response (*E*) by the agonist-occupied receptor (*RA*). This proceeds at a rate defined by *k_τ_*, the transduction rate constant, which is the initial rate of response generation by the agonist-occupied receptor. Receptor desensitization is represented by transformation of *RA* into the inactive receptor *R*_0_*A* that does not generate a response (because it can’t couple to *E_P_*). The rate constant for desensitization is *k_DES_*. Degradation of response is represented by decay of *E* to *D*, governed by the degradation rate constant *k_D_*. This model is an extension the original kinetic operational model (Hoare et al., 2018), extended to incorporate receptor desensitization.

Regulation of signaling can be incorporated into this model. The regulation mechanisms are receptor desensitization (Fig. 1, yellow region), response degradation (Fig. 1, pink region), and depletion of response precursor. Receptor desensitization results in a decrease in the number of receptors that can couple to the signal transduction pathway to generate the response (the number of active receptors). For G-protein-coupled receptors, the number of active receptors is reduced typically by receptor phosphorylation and subsequent arrestin binding (Krupnick and Benovic, 1998). This is easily accommodated within the framework of the model because the receptor concentration is a variable within it. Receptor desensitization can be represented by a decrease of the agonist-occupied receptor concentration over time (Fig. 1, yellow region). Response degradation is the process by which the signal, once generated, is cleared over time. Examples of response degradation processes include breakdown of second messenger molecules, such as cAMP by phosphodiesterases (Chang, 1968), and de-activation of G-protein by hydrolysis of bound GTP to GDP by the intrinsic GTPase activity of the G-protein (Gilman, 1987). Some signals are decreased by clearance of the signaling species from the relevant compartment, for example efflux of cytosolic Ca^2+^ ions (Carafoli, 1991). Response degradation is incorporated into the model as an exponential decay of the response (Fig. 1, pink region), as described previously (Hoare et al., 2018). Signal transduction can also be dampened over time by a third mechanism, depletion of the response precursor. The most familiar example is in calcium signaling, in which depletion of Ca^2+^ from intracellular stores limits the amount of Ca^2+^ that can be released into the cytosol (Yu and Hinkle, 2000). This property can be incorporated into the model as a decrease of response precursor over time, as described previously (Hoare et al., 2018).

It is well known that additional processes effect GPCR responses, especially over longer timeframes. The receptor can resensitize and so re-participate in signaling (Ferguson, 2001). Recently, it has emerged that regulatory processes can result in prolonged signaling. For example, receptor internalization can result in a long-lived signaling complex in intracellular vesicles (Ferrandon et al., 2009; Hothersall et al., 2016). These mechanisms can be incorporated by extending the model and such models are currently being evaluated.

### 2.2. Formulating the model

The model was formulated in a manner that yields measurable parameters of ligand efficacy and response regulation that can be applied to drug discovery, for example to establish structure-activity relationships in lead optimization (Kenakin, 2009a).

#### 2.2.1. Efficacy – response generation

In the response generation step, agonist-bound receptor (*RA*) interacts with a precursor of the response (*E_P_*) and converts it to the response (*E*). The rate of response generation is defined by mass action, i.e. is a function of the concentration of the interacting species ([*RA*] and *E_P_*) and a rate constant (*k_E_*). The rate is termed the transduction rate constant (*k_τ_*). It is defined as:

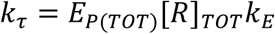

where *E_P_*_(*TOT*)_ is the total response precursor available in the system and [*R*]*_TOT_* the total concentration of receptors. *k_τ_* can be described as the initial rate of response generation. It is the rate of response generation before the response is appreciably affected by regulation processes. It is analogous to the initial rate of enzyme activity, [*S*]*_TOT_*[*E*]*_TOT_k_CAT_*, where *S* is substrate, *E* enzyme and *k_CAT_* the catalytic rate constant.

*k_τ_* is the efficacy of the agonist. It is the macroscopic efficacy of the agonist in the system being measured and as such can be viewed as a kinetic analogue of traditional efficacy parameters, such as *τ* of the operational model (Black and Leff, 1983). *k_τ_* is determined by the agonist because it is dependent on *k_E_*, the microscopic efficiency of the agonist for generating the response. In other words, agonists with different efficacies manifest different values of *k_τ_* for the response being measured because they differ in the value of *k_E_*. *k_τ_* is also defined by system parameters that are independent of the agonist, specifically [*R*]*_TOT_* and *E_P_*_(*TOT*)_. As a result, the value of *k_τ_* for a single agonist can differ between response systems. This can occur for three reasons: differing receptor concentration (different [*R*]*_TOT_*); differing levels of response precursor (different *E_P_*_(*TOT*)_; and differing types of response precursor that the receptor is coupling to (different *k_E_*).

Experimentally, *k_τ_* can be measured in routine pharmacological assays using familiar curve fitting methods and equations, as described in Section 3. A maximally-stimulating concentration of agonist can be used to measure *k_τ_*. This property enables measurement of agonist efficacy regardless of the kinetics of agonist binding to the receptor, as described below (Section 2.2.3). Importantly, *k_τ_* can be measured independently of the rate(s) of response regulation, as described in Section 2.2.2.

#### 2.2.2. Response regulation

The canonical signaling regulation mechanisms in the model are receptor desensitization and response degradation (Fig. 1). In the desensitization mechanism, the active receptor *RA* is desensitized, forming the inactive receptor, *R*_0_*A*, which cannot couple to the response precursor (Fig. 1, yellow region). Receptor desensitization is modeled as an exponential decay of the active receptor concentration. It is assumed to be an exponential decay process because arrestin binding to the receptor has been shown to proceed according to an exponential process (for example, see (Berglund et al., 2003)). The rate is defined by the receptor desensitization rate constant *k_DES_*. The half-time for receptor desensitization is 0.693 / *k_DES_*. *k_DES_* is an agonist-dependent parameter because it is dependent on the nature of the agonist bound to the receptor. Regarding response degradation, the mechanism is assumed to be an exponential decay process, as described previously (Hoare et al., 2018) (Fig. 1, pink region). The rate is defined by *k_D_*, the response degradation rate constant, with the half-time for degradation being 0.693 / *k_D_*. This rate is independent of the agonist because response degradation occurs downstream of the receptor.

It is critical to note that the agonist efficacy parameter *k_τ_* is independent of the system regulation parameters (*k_DES_* and *k_D_*) in this model. This means that agonist efficacy can be measured in a system in which the response is regulated, as shown in Section 3. This emerges from the microscopic features of the model. At the level of a single receptor, the process of response generation is independent from the processes of receptor desensitization and response degradation. Mathematically, the capacity of a single agonist-bound receptor to generate the response is governed by *k_E_* (see green panel in Fig. 1), not by *k_DES_* or *k_D_*. This is because mechanistically: 1) receptor desensitization only affects the number of active receptors, not the capacity of a single active receptor to generate the response. 2) Response degradation (*k_D_*) does not affect response generation because it is downstream of the receptor. From a macroscopic perspective, the efficacy parameter measured in the experiment, *k_τ_*, is independent of the regulation parameters for two reasons: 1) It is the initial rate of signal transduction, i.e. the rate before the response is affected by the regulation processes. 2) The regulation parameters *k_DES_* and *k_D_* are absent in the expression defining *k_τ_* (*E_P_*_(*TOT*)_[*R*]*_TOT_k_E_*).

#### 2.2.3. Agonist binding

In classical pharmacological models, it is implicitly assumed that agonist-receptor binding is at equilibrium at the time point at which the response is measured (Bdioui et al., 2018; Black and Leff, 1983; Colquhoun, 1998; Rang, 2006; Slack and Hall, 2012). In a kinetic model, it cannot be generally assumed that agonist is at equilibrium with the receptor at all time points of response measurement (unless a maximally-effective concentration of agonist is employed – see below). The equilibrium assumption is probably reasonable for low potency ligands when the response progresses over timescales of at least several minutes, for example second messenger generation and gene expression (Bdioui et al., 2018). Such ligands (EC_50_ in the micromolar range) equilibrate rapidly even at the low concentrations required to span the concentration-response curve (equilibration time < 1 min) (Hoare et al., 2018). However, high potency agonists (EC_50_ in the low nM range) equilibrate more slowly, especially at the low concentrations required to span the concentration-response curve (at least several minutes). In addition, lack of equilibration is likely to be an issue for responses that are extremely rapid (Bdioui et al., 2018; Charlton and Vauquelin, 2010). For example, the rise of intracellular Ca^2+^ occurs within a few seconds of agonist application. This problem can be solved by incorporating the kinetics of agonist binding into the model. This approach is applied for the new models in this study (see Appendix A, Supplementary Fig. S2, Supplemental information). This approach was applied to the original kinetic operational model (Hoare et al., 2018) and a variant of the operational model that assumes agonist binding is rate-limiting (Slack and Hall, 2012).

Fortunately, agonist efficacy can be determined in a way that minimizes the equilibration problem. This is by virtue of the fact that a saturating concentration of agonist can be used to measure agonist efficacy (see Section 3). The higher the concentration of ligand, the more rapidly it associates with the receptor, due to mass action. For example, let us consider a slowly-equilibrating agonist with an equilibrium dissociation constant of 10 nM and a *t*_1/2_ for association, at 10 nM, of 10 min. If this agonist is applied at the maximal concentration typically employed in in vitro signaling assays (10 μM), association with the receptor is greatly accelerated (association *t*_1/2_ of 1 second). [These values are calculated from the equation for the observed association rate of receptor-ligand association, *k_obs_* = [*A*]*k*_1_ + *k*_−1_, with a *k*_1_ (association rate constant) value of 3.5 × 10^6^ M^−1^min^−1^ and *k*_–1_ (dissociation rate constant) value of 0.035 min^−1^].

### 2.4. Equation format

In a kinetic functional assay, the response to agonist is measured over multiple time points and the response value is plotted against time. Equations were derived for analyzing response vs time data to obtain estimates of the model parameters (Appendix A). The equations are of a form that can be used by curve fitting software commonly-employed in drug discovery and receptor research, for example Prism (GraphPad Software, Inc.) XFfit (ID Business Solutions Ltd.) and SigmaPlot (Systat Software, Inc.). These equations take the analytic form, *y* = *f*(*t*), where *y* is response and *f*(*t*) is a function of time and pharmacological parameters. This is referred to as an *E vs t* equation in this study. [Note that certain programs allow fitting of data to differential equations rather than their analytic solutions, such as Dynafit from BioKin Ltd, (Kuzmic, 2009).]

Some of the models resulted in systems of nonlinear differential equations, which are often difficult to solve analytically. Surprisingly, analytical solutions were obtained for most of these systems. Importantly, this allowed extension of the model to incorporate agonist binding kinetics (for example, Appendix A.5) and to accommodate simultaneous receptor desensitization and precursor depletion (Appendix A.9).

## 3. Results

The manifestation of the model in response time course data is first evaluated using simulated data generated using the model equations. The models are then verified by comparison with historical experimental data. In these historical studies, regulation mechanisms were manipulated (effectively deleted), enabling a piecewise verification of the model. The model is then applied to the experimental data to obtain estimates of the model parameters, using standard curve-fitting procedures. In the course of this investigation, a pattern emerged in which the shape of the time course was determined by the number of regulation processes. No regulation processes results in a straight line (Section 3.1). A single regulation process results in an “association” exponential curve, here termed a horizontal exponential curve (Section 3.2). Two processes (an input process and an output process) results in a rise-and-fall exponential curve (Section 3.3). This property enabled a generalized approach to be used to analyze these data (Sections 3.2.4 and 3.3.4). In this approach, agonist efficacy (*k_τ_*) could be measured without knowledge of the underlying regulation mechanism. More complex curves resulted from multiple mechanisms acting in concert.

The models were considered in order of increasing complexity:

1. An unregulated response, giving a straight line E vs t profile (Section 3.1).
2. A single regulation mechanism, giving a horizontal exponential curve (Section 3.2).
3. Receptor desensitization or precursor depletion (input process), and response degradation (output process), resulting in a rise-and-fall curve (Section 3.3).
4. More complex mechanisms resulting in more complex curve shapes (Section 3.4).

Some of the models were described in the original study – Models 1,3,4 and 8 (Hoare et al., 2018). Here they are considered again in the context of regulation of signaling and in order to enable comparison with the new models.

In this section the models are considered for a maximally-stimulating concentration of agonist, which enables measurement of efficacy (*k_τ_*) and the regulation parameters. Data for multiple concentrations of agonist are given in Supplementary information.

### 3.1. Unregulated response – straight line profile (Model 1)

In the absence of any regulation mechanism, the model predicts a linear increase of the response over time. In other words the *E vs t* profile is a straight line. This makes sense because there is nothing to stop the signal being generated, and there is no degradation of the signal, so the response accumulates continuously. This mechanism is shown schematically in Fig. 1 (green-shaded region) and in Scheme 1 below:

**Scheme 1.**
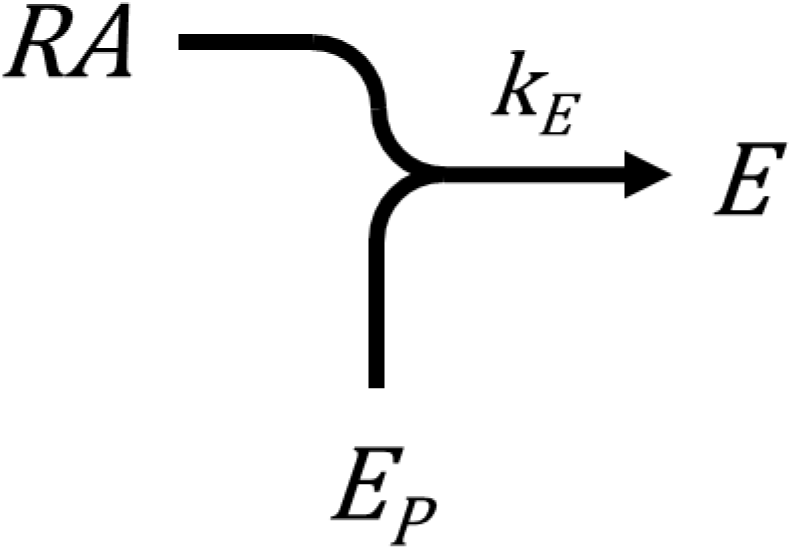

This model has been described previously, being the minimal mechanism of the kinetic operational model (Hoare et al., 2018). Response is generated from response precursor, at a rate governed by *k_τ_*. The linear time course profile predicted by this model is shown in Fig 2A. The linear profile is also evident in the *E vs t* equation for the model, derived in Appendix A.1 (Eq. 1):

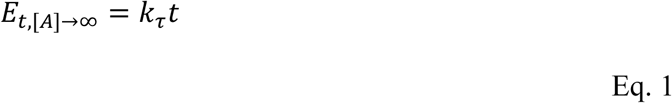

**Fig. 2.**
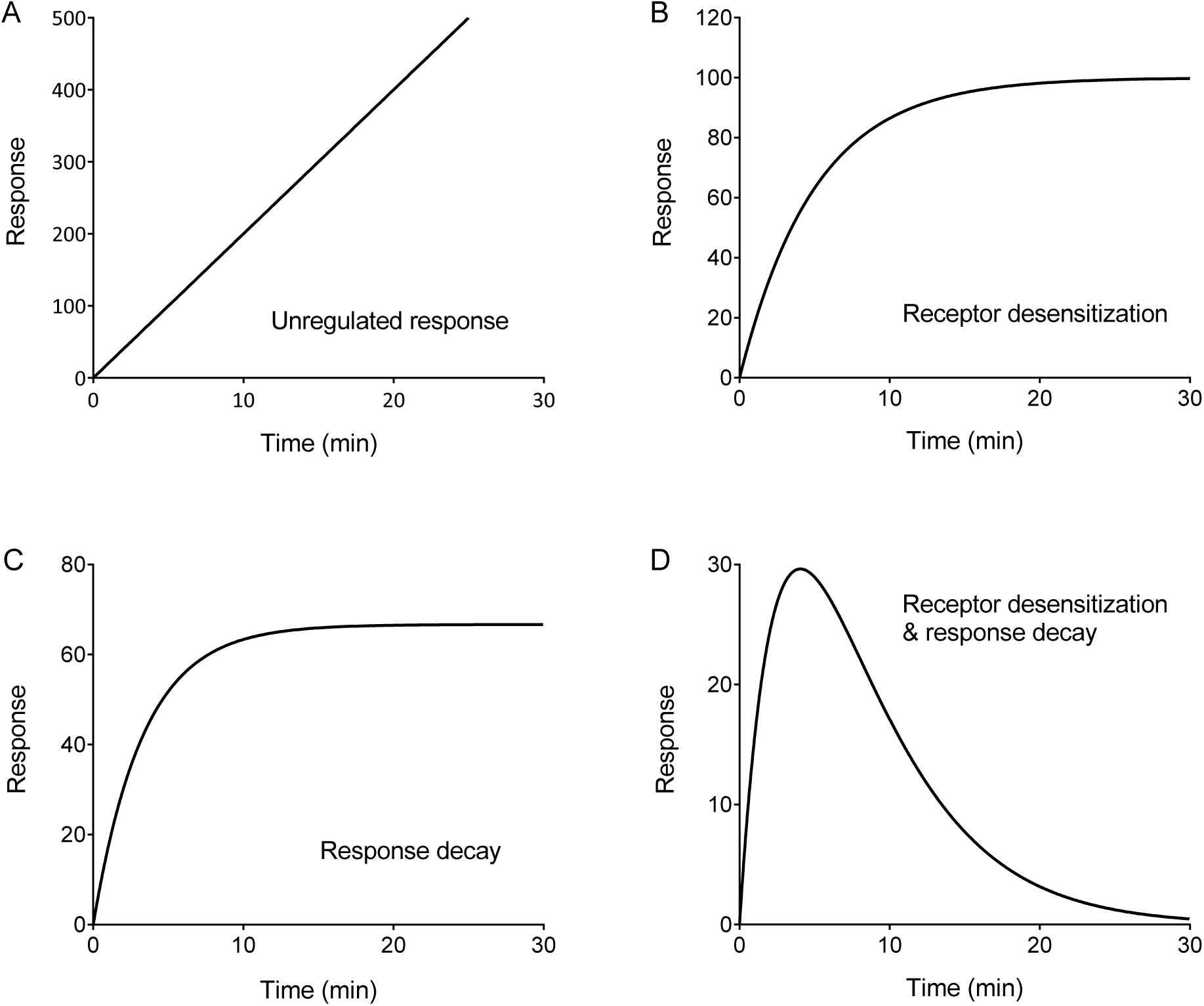
Examples of curve shapes resulting from kinetic response models. (a) A straight line profile results when there is no response regulation (Model 1, Section 3.1). (b) and (c) A horizontal exponential curve results from a single regulation process [receptor desensitization (b) or response degradation (c), Models 2 and 3, Sections 3.2.1. and 3.2.2.]. (d) A rise-and-fall exponential profile results when two regulation processes are in operation, receptor desensitization and response degradation (Model 6, Section 3.3.1.). Data were simulated with Eq. 1 (a), Eq. 2 (b), Eq. 3 (c) and Eq. 6 (d) with the following parameter values: *k_τ_*, 20 response units.min^−1^; *k_DES_*, 0.20 min^−1^; *k_D_*, 0.3 min^−1^.

This is the equation for a straight line, where the gradient is *k_τ_* and the intercept is zero. (Note this is the equation for a maximally-stimulating concentration of agonist.)

In order to validate the model, the literature was searched for GPCR responses with minimal regulation, i.e. with minimal receptor desensitization, response degradation and precursor depletion. The inositol phosphates (IP) signaling pathway is useful for this evaluation because response degradation can be completely blocked by Li^+^, which inhibits the terminal step of inositol phosphate catabolism (dephosphorylation of inositol monophosphate by inositol monophosphatase) (Naccarato et al., 1974). In this response there is minimal precursor depletion because the precursor, phosphatidylinositol 4,5-bisphosphate, is continuously generated (Xu et al., 2003). Consequently, an inositol phosphates assay run in the presence of Li^+^ can be reasonably assumed to be minimally impacted by two of the regulation mechanisms, response degradation and precursor depletion. This leaves us with the remaining mechanism, receptor desensitization.

Blocking receptor desensitization in systems with minimal degradation and depletion results in a linear time course of the response, consistent with the model. This is shown in the following examples. Receptor desensitization in the inositol phosphates pathway has been blocked by genetic ablation of β-arrestin. Embryonic fibroblasts were isolated from arrestin knock-out mice and GPCR signaling dynamics evaluated in these cells (Kohout et al., 2001). In this system, the IP time course was a straight line. This was shown for angiotensin II via the AT_1_ angiotensin receptor (Kohout et al., 2001), and for thrombin via the proteinase-activated PAR1 receptor (Paing et al., 2002). The AT_1_ receptor data are reproduced in Fig. 3. A second example of diminished receptor desensitization is provided by the mammalian gonadotropin releasing hormone GnRH_1_ receptor. This receptor lacks the C-terminal tail (Eidne et al., 1992). This region of the receptor is a critical determinant of canonical receptor desensitization because it contains the sites of receptor phosphorylation and is a determinant of arrestin binding (Krupnick and Benovic, 1998). A linear time course of inositol phosphates generation has been observed for the mammalian GnRH_1_ receptor expressed endogenously and heterologously in numerous cell lines (Davidson et al., 1994; Davis et al., 1986; Heding et al., 1998). (Data from two of these studies are reproduced in Figs. 4 and 5.) [Longer durations of GnRH exposure (typically > 20 min) results in attenuation of the response, owing to arrestin-independent receptor internalization (Heding et al., 2000).]

**Fig. 3.**
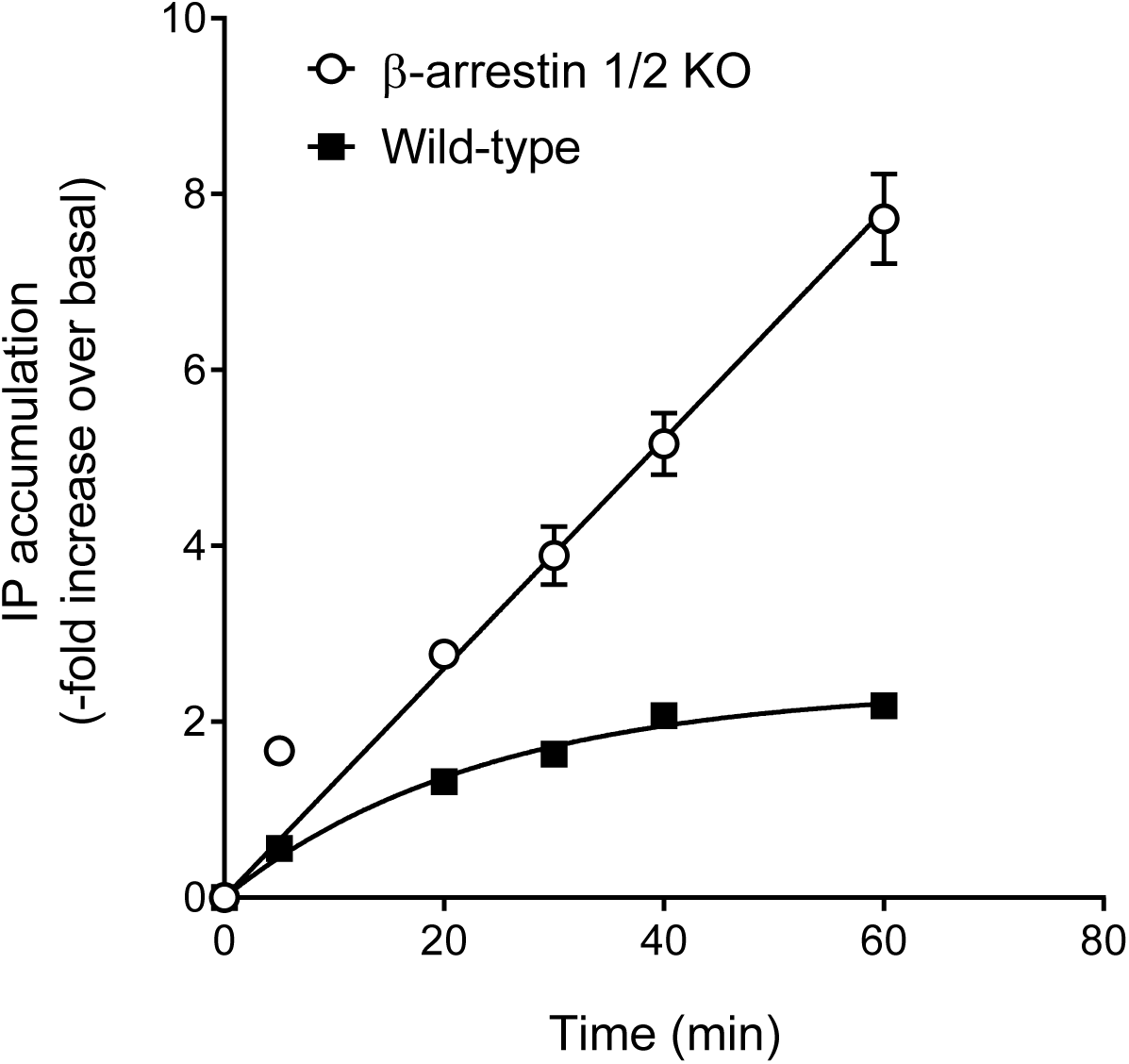
Receptor desensitization: Angiotensin-stimulated inositol phosphates accumulation in mouse embryonic fibroblasts via the AT_1_ angiotensin receptor (Kohout et al., 2001). Mouse fibroblasts were isolated from arrestin double knock out mice or wild type mice. In the absence of arrestin, the response is unregulated (straight line). In the presence of arrestin, the receptor becomes desensitized on application of the agonist, resulting in the horizontal exponential curve, consistent with the kinetic desensitization model (Section 3.2.1.). Data for wild-type cells were fit to the receptor desensitization model (horizontal exponential equation, Section 3.2.1), giving a *k_τ_* value of 0.10-fold.min^−1^, and a *k_DES_* value of 0.042 min^−1^ (translating to a half-time for receptor desensitization of 17 min). Data for the arrestin knock out cells were fit to the no-regulation model (straight-line equation, Section 3.1), giving a *k_τ_* value of 0.13-fold over basal.min^−1^. A maximally-stimulating concentration of angiotensin II was applied (100 nM). Data are from Fig. 3C of (Kohout et al., 2001), extracted using a plot digitizer [WebPlotDigitizer (Rohatgi, 2018)].

**Fig. 4.**
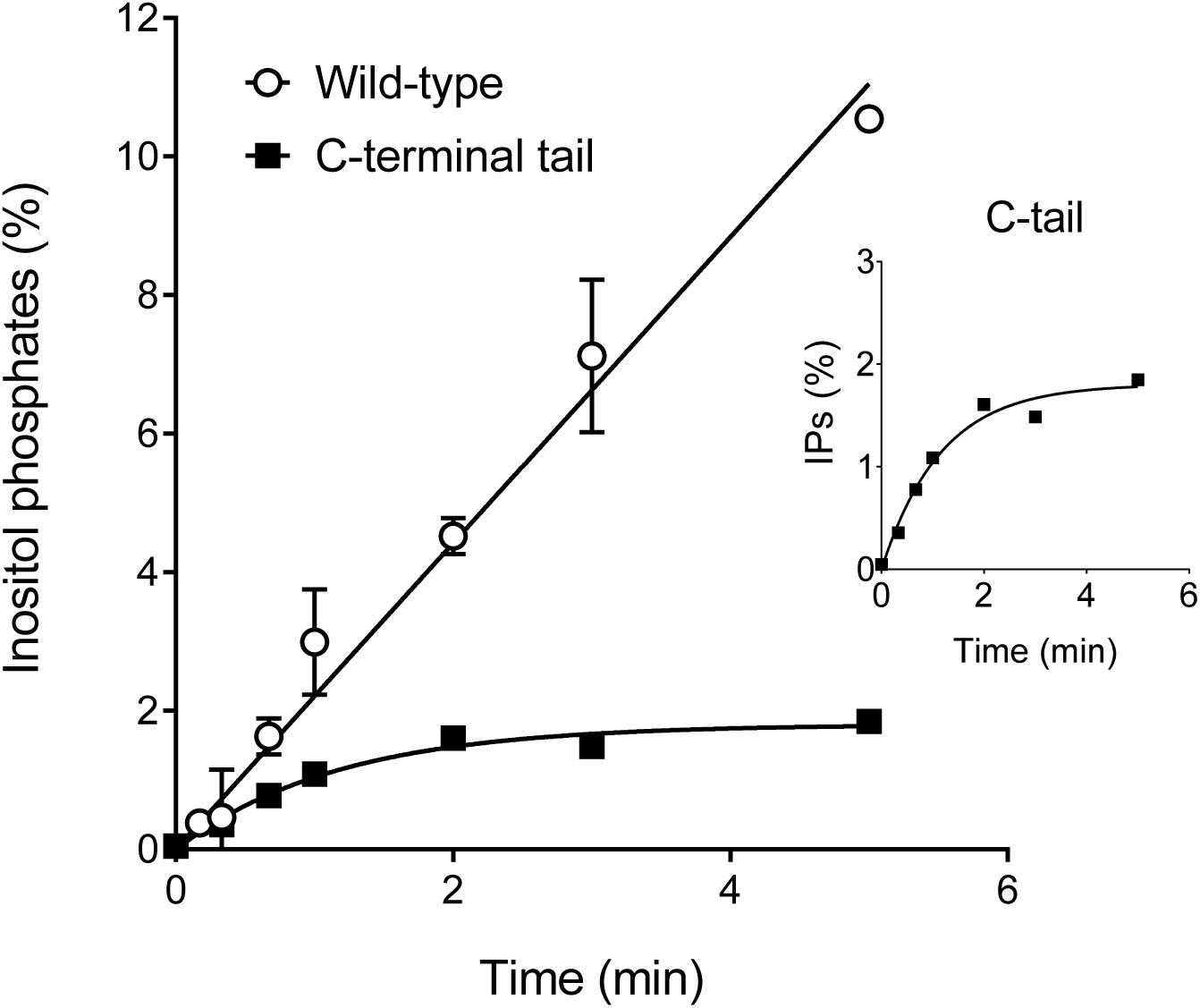
Receptor desensitization: GnRH-stimulated inositol phosphates accumulation via the rat GnRH_1_ receptor with a C-terminal extension in HEK293 cells (Heding et al., 1998). The mammalian GnRH_1_ receptor lacks a C-terminal tail and so does not undergo arrestin-mediated receptor desensitization. Incorporating a C-terminal tail from a receptor that does (TRH receptor) results in arrestin-mediated receptor desensitization (Heding et al., 2000; Heding et al., 1998). Data for the C-terminally-extended receptor were fit to the receptor desensitization model (horizontal exponential equation, Section 3.2.1), giving a *k_τ_* value of 1.5%.min^−1^, and a *k_DES_* value of 0.86 min^−1^ (translating to a half-time for receptor desensitization of 0.81 min). Data for the wild-type receptor were fit to the no-regulation model (straight-line equation, Section 3.1), giving a *k_τ_* value of 2.2%.min^−1^. A maximally-stimulating concentration of GnRH was applied (1 μM). Data are from Fig 4 of (Heding et al., 1998), with basal response subtracted, extracted using a plot digitizer [WebPlotDigitizer (Rohatgi, 2018)].

**Fig. 5.**
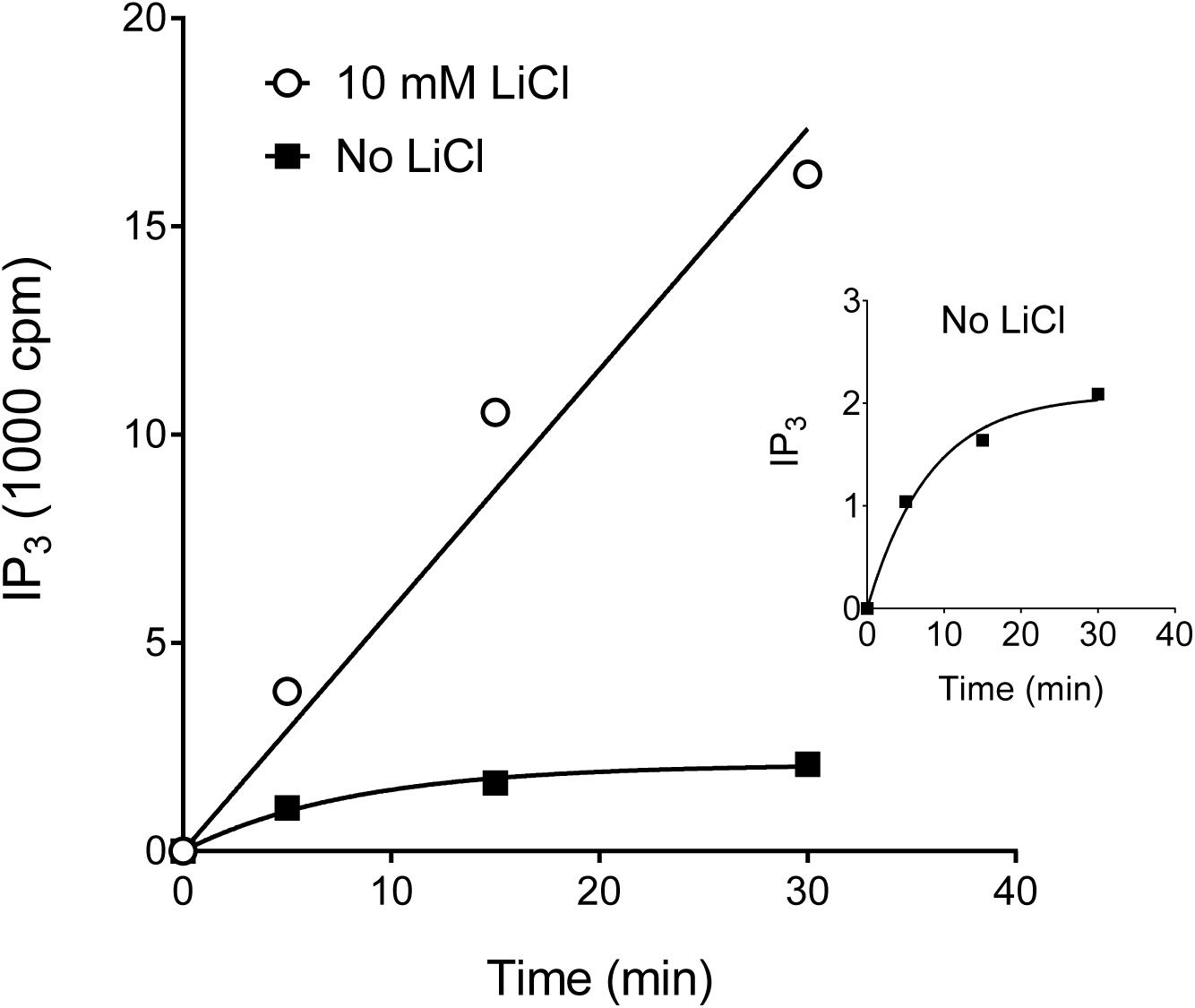
Response degradation: GnRH-stimulated inositol phosphates accumulation in rat granulosa cells (Davis et al., 1986). In the absence of Li^+^, inositol phosphates are degraded, resulting in the horizontal exponential curve, consistent with the kinetic response degradation model (Section 3.2.2.). Data for the absence of Li^+^ were fit to the response degradation model (horizontal exponential equation, Section 3.2.2), giving a *k_τ_* value of 250 cpm IP_3_.min^−1^. Data for the presence of Li^+^ were fit to the no-regulation model (straight-line equation, Section 3.1), giving a *k_τ_* value of 580 cpm IP_3_.min^−1^. A maximally-stimulating concentration of GnRH was applied (85 nM). Data are from Fig. 3 of (Davis et al., 1986), extracted using a plot digitizer [WebPlotDigitizer (Rohatgi, 2018)].

The *k_τ_* value can be estimated by fitting experimental data to Eq. 1. This can be done by fitting the *E vs t* data for maximally-stimulating concentration of agonist to a generic straight line equation:

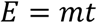

where *m* is the gradient of the line. *k_τ_* is equal to *m* (evident from Eq. 1). This analysis is applied here to the arrestin knock out and GnRH_1_ receptor data (Figs. 3–5). For angiotensin II-stimulated inositol phosphates accumulation via the AT_1_ receptor in arrestin knockout mouse embryonic fibroblasts (Kohout et al., 2001), the *k_τ_* value is 0.13-fold over basal.min^−1^ (Fig 3). For GnRH-stimulated inositol phosphates accumulation via the GnRH_1_ receptor in HEK293 cells (Heding et al., 1998), the *k_τ_* value is 2.2% of total added radioactivity.min^−1^ (Fig. 4). For GnRH-stimulated IP_3_ accumulation via the GnRH_1_ receptor endogenously-expressed in rat granulosa cells (Davis et al., 1986), the *k_τ_* value is 580 cpm IP_3_.min^−1^ (Fig. 5).

### 3.2. Single regulation mechanism – horizontal exponential profile

In this section models are considered in which a single regulation mechanism is in operation – receptor desensitization, response degradation or depletion of response precursor. In all cases the *E vs t* profile is a horizontal exponential curve. (The horizontal exponential curve is defined by the equation *y* = Plateau × (1 –*e^k.t^*). A familiar example is the ligand binding association curve.) This commonality enables a mechanism-agnostic approach to be applied to analyze horizontal exponential time course data (Section 3.2.4.).

#### 3.2.1 Receptor desensitization (Model 2)

Receptor desensitization is represented in the model as a reduction of the active receptor concentration over time (“active” meaning the capacity to generate a response). The basic model of receptor desensitization is shown schematically in Fig. 1 (green and yellow shaded regions) and in Scheme 2 below:

**Scheme 2.**
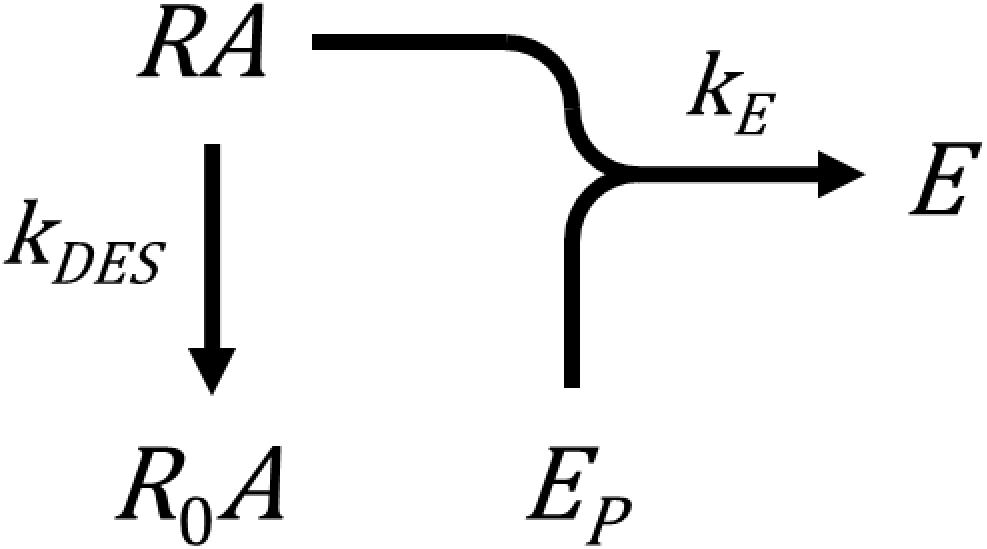

The time course profile predicted by this model is a horizontal exponential curve (Fig 2B). This makes sense intuitively. At early times response is generated rapidly. The response then slows, because response generation is attenuated by the loss of active receptor. Ultimately the response approaches a limit (the horizontal plateau). At this limit the response level does not change because no new response is being generated and the existing response is not degraded. The equation defining response over time for the desensitization model, for a maximally-stimulating concentration of agonist, is Eq. 2, derived in Appendix A.2:

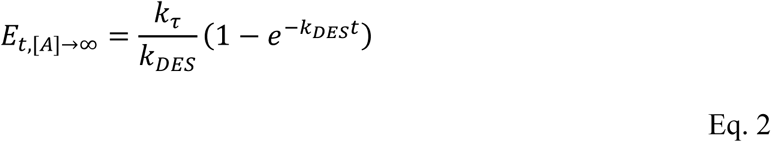

The effect of receptor desensitization on the shape of the time course can be tested by introducing receptor desensitization into a system that is unregulated. This can be done using the systems described in Section 3.1 (Figs. 3 and 4). In the arrestin knock out cell system the effect of desensitization can be tested by comparing wild-type cells that express β-arrestin with the knockout cells that do not. For the AT_1_ inositol phosphate response (Kohout et al., 2001), the shape of the *E vs t* profile is a horizontal exponential curve when arrestin is expressed (Fig. 3), implying receptor desensitization results in a horizontal exponential curve. The same finding has been reported for the PAR1 receptor (Paing et al., 2002). For the GnRH_1_ receptor system, receptor desensitization has been introduced by incorporating a C-terminal tail into the receptor, from a receptor that is desensitized (the thyrotropin-releasing hormone TRH1 receptor) (Heding et al., 1998). This results in a horizontal exponential curve (Fig. 4). The horizontal exponential time course in these receptor desensitization systems provides experimental verification for the desensitization model.

The model parameters can be estimated by fitting experimental time course data to Eq. 2. A simple way to do this is to analyze the data using a generic horizontal exponential equation. First, the *E vs t* data for a maximally-stimulating concentration of agonist is fitted to the following generic horizontal exponential equation:

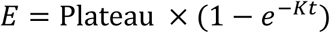

“Plateau” is the response at the horizontal asymptote (i.e. *E* as *t* → ∞) and K is the observed rate constant. This can be done in Prism using the built-in “One phase association” equation (Motulsky, 2019a). Fitting the *E vs t* data to this equation gives an estimate of Plateau and of K. From these values it is possible to calculate *k_τ_* and *k_DES_*, providing a maximally-stimulating concentration of agonist is employed. Under this condition, *k_τ_* is simply the Plateau multiplied by the rate constant:

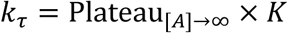

*k_DES_* is equal to the rate constant K:

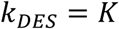

This analysis is now applied to the data in Figs. 3 and 4. For the AT_1_ receptor response in arrestin-expressing cells, the Plateau value was 2.4-fold, and the K value was 0.042 min^−1^ (Fig. 3). Multiplying Plateau by K gives the *k_τ_* value 0.10-fold.min^−1^. If this analysis is valid, the *k_τ_* value should be equal to the value in the non-desensitized system (arrestin knock-out cells), assuming the only difference between the systems is receptor desensitization. The data are in reasonable agreement with this assumption; the *k_τ_* value in arrestin knockout cells was 0.13-fold.min^−1^ (Fig. 3). The *k_DES_* value is equal to K. The *k_DES_* value was 0.042 min^−1^. This translates to a half-time of desensitization of 17 min for the angiotensin II-occupied AT_1_ receptor in this system. For the GnRH1 receptor engineered with a C-terminal tail, the Plateau value was 1.8 % and the K value was 0.86 min^−1^ (Fig 4). The calculated *k_τ_* value is 1.5 %.min^−1^, close to the value for the non-desensitized receptor of (2.2 %.min^−1^). The *k_DES_* value, equal to K, was 0.86 min^−1^. This translates to a half-time of desensitization of 0.81 min for the C-terminally-extended GnRH_1_ receptor in this system.

#### 3.2.2. Response degradation (Models 3 and 4)

Response degradation was incorporated into the model as an exponential decay of the response, as described previously (Hoare et al., 2018). The model, Model 3, is shown schematically in Fig. 1 (green and pink shaded regions) and in Scheme 3 below:

**Scheme 3.**
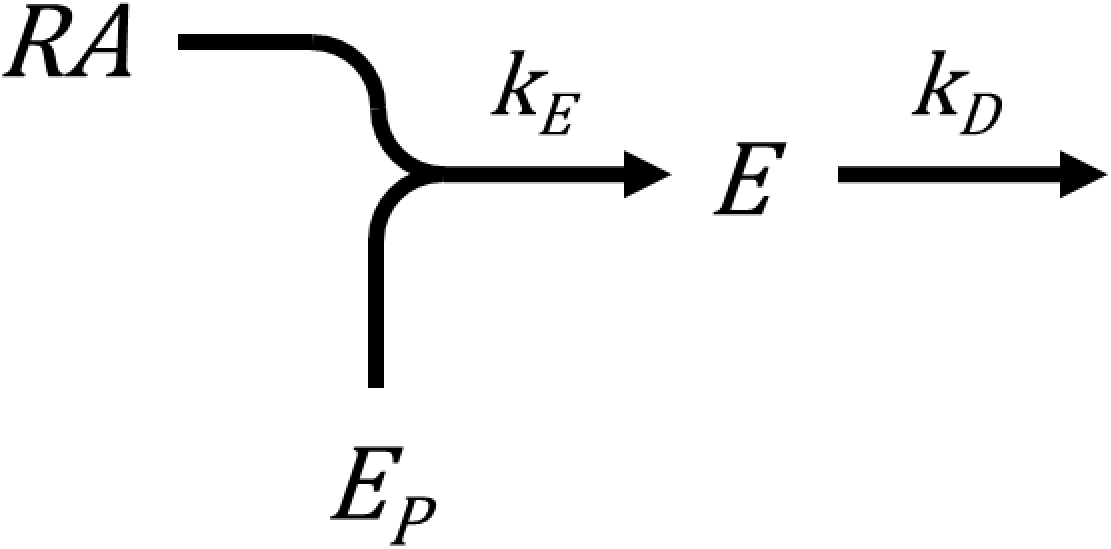

The *E vs t* profile predicted by this model is a horizontal exponential curve (Fig. 2C). This can be rationalized as follows. At first, the response level increases rapidly because the response generation process predominates. As time progresses, the rate of increase slows, as response degradation increases. Finally, a plateau of the response approached, at which there is a steady-state where the rate of response generation equals the rate of response degradation. The equation for this model, at a maximally-stimulating concentration of agonist, is Eq 8, derived in Appendix A.3:

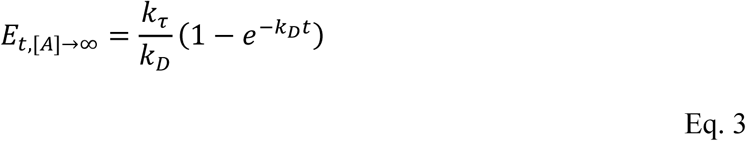

A variant of this model incorporates recycling of the response back to the response precursor (Hoare et al., 2018). It is shown in Scheme 4 below:

**Scheme 4.**
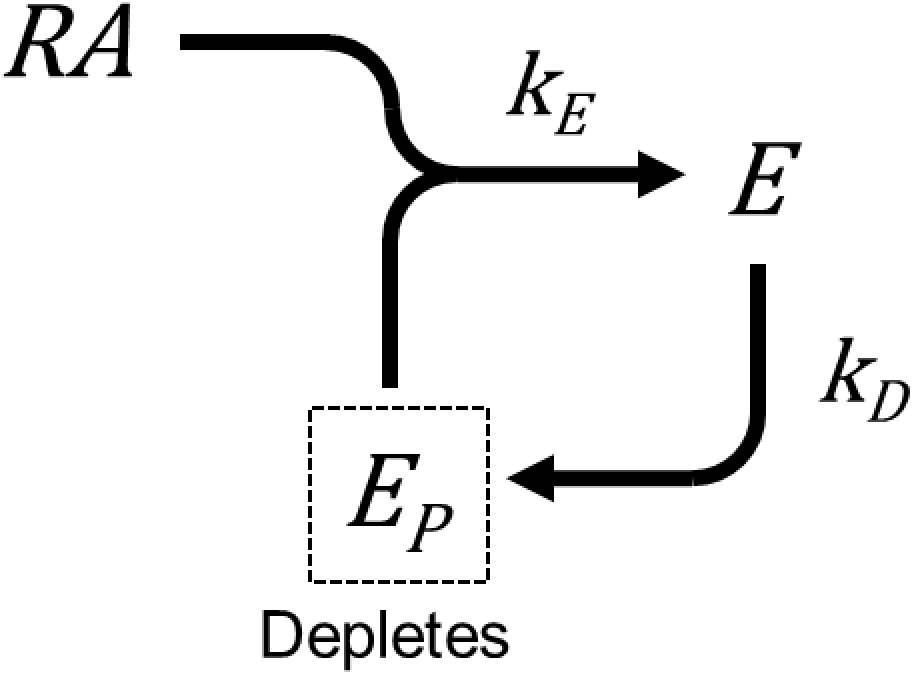

The equation defining this model (Model 4) is a horizontal exponential equation, derived in Appendix A.4 (Eq. 4),

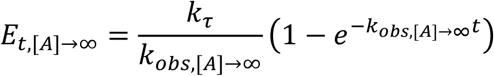

where,

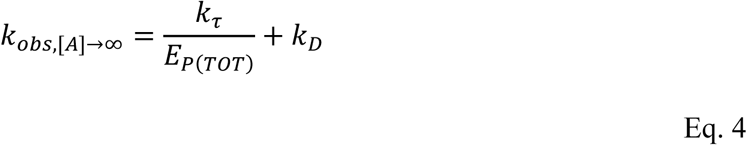

This variant of the degradation model is similar to the one without recycling. The only difference in the equation is that *k*_*obs*,[*A*]→∞_ replaces *k_D_*.

The effect of response degradation on the shape of the time course can be investigated by leaving out the response degradation inhibitor. For the inositol phosphates response this can be done by omitting Li^+^, the inhibitor of the terminal step of inositol dephosphorylation (Naccarato et al., 1974). Using the GnRH_1_ receptor enables degradation to be examined in isolation from receptor desensitization because this receptor does not desensitize acutely (Section 3.1). Data from this system are reproduced in Fig. 5 (Davis et al., 1986). Omitting Li^+^ results in a horizontal exponential curve (Fig. 5), implying response degradation results in a horizontal exponential curve. This finding is consistent with the model.

A limitation of the degradation inhibitor approach is noted here. The most familiar example of response degradation is breakdown of cAMP by phosphodiesterase (PDE) enzymes. This process is inhibited, and cAMP elevated, by PDE inhibitors. This treatment is used routinely in measuring cAMP signaling mediated by GPCRs. However, standard methods do not completely inhibit PDE activity, such that some degradation of cAMP remains. This is because of solubility limits of the most-commonly used PDE inhibitor, isobutylmethylxanthine. A concentration commonly applied is 0.5 mM, which inhibits degradation by only 75% (Schulz and Mailman, 1984). Subtype-selectivity is an issue with another commonly used inhibitor, rolipram. This compound is selective for PDE4 (Houslay et al., 2005) but cell lines used routinely for the study of GPCRs express multiple PDE subtypes and appreciable cAMP levels can be detected in the presence of subtype-selective PDE inhibitors (Motte et al., 2017). Incomplete inhibition of cAMP breakdown can complicate the interpretation of the effect of response degradation on cAMP signaling (Gimenez et al., 2015; Xin et al., 2008). In the context of the kinetic model in this section, application of a response degradation inhibitor at a partially effective concentration does not linearize the *E vs t* profile but instead results in a decrease of the rate constant and an increase of the plateau (data not shown).

The model parameters can be estimated by fitting experimental time course data for a maximally-stimulating agonist concentration to the generic horizontal exponential equation:

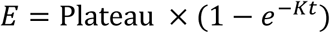

*k_τ_* can be determined by multiplying the Plateau by K, as is evident from Eqs. 8 and 11. This method can be applied whether or not the response is recycled, evident from inspecting the equations. This analysis was applied to the GnRH signaling data in Fig. 5. The Plateau value was 2,100 cpm IP_3_ and the K value 0.12 min^−1^. The resulting value of *k_τ_* was 250 cpm IP_3_.min^−1^. This value was in the same range as that for the unregulated response (580 cpm IP_3_.min^−1^, for the response in the presence of Li^+^, Section 3.1).

It is not possible to determine *k_D_* using this approach unless it is known whether or not response is recycled, i.e. whether Scheme 4 or Scheme 5 applies. This is because the definition of K is different in these two scenarios. If response is not recycled, K is equal to *k_D_*. If response is recycled, K is equal to *k_τ_* / *E_P_*_(*TOT*)_ + *k_D_*.

#### 3.2.3. Depletion of response precursor (Model 5)

In this mechanism, the only process of response limitation is depletion of response precursor. The precursor is depleted because it is converted to the response by the response-generation process. This mechanism results in a horizontal exponential profile. This model, Model 5, is rarely encountered and is presented here for the purpose of completeness.

This model is represented by Scheme 5:

**Scheme 5.**
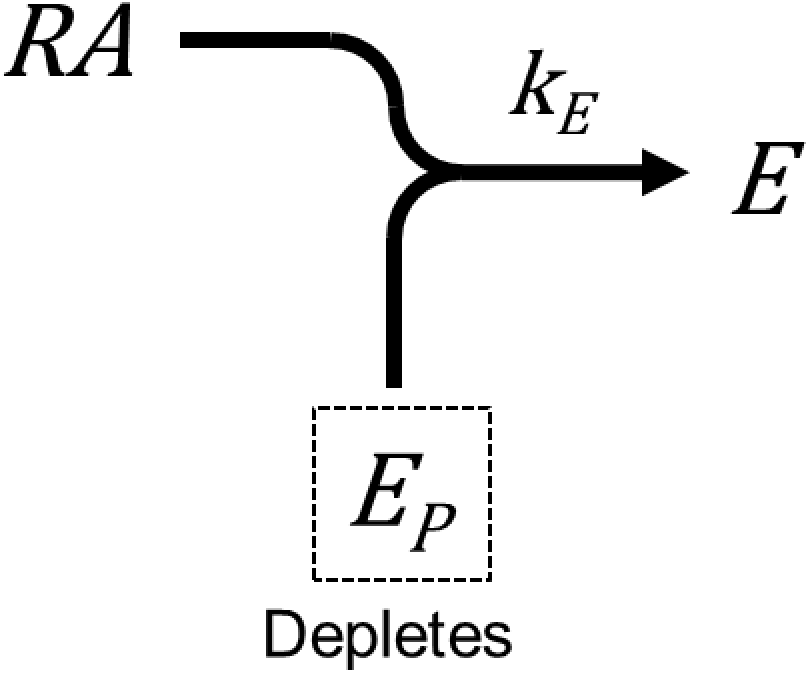

The equation defining this mechanism, for a maximally-stimulating concentration of agonist is Eq. 5, derived in Appendix A.5:

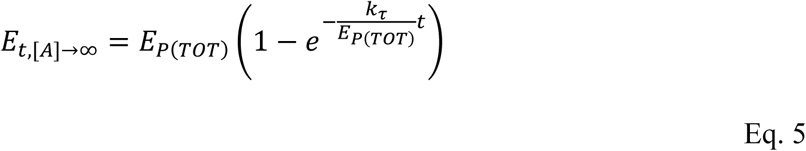

This is a horizontal exponential equation. Analysis using the generic horizontal exponential equation (Section 3.2.1) gives a Plateau value corresponding to *E_P_*_(*TOT*)_ (the total amount of response precursor) and a K value corresponding to *k_τ_* / *E_P_*_(*TOT*)_. *k_τ_* can be determined by multiplying the Plateau value by the K value, as is done for the other horizontal exponential models. An example from the literature can be found in early studies of glucagon-stimulated cAMP accumulation in hepatic membranes (Pohl et al., 1971). This system is minimally regulated, lacking receptor desensitization machinery owing to the use of washed membranes, and lacking phosphodiesterase activity (Pohl et al., 1969). Depletion of response precursor results from the absence of an ATP-regenerating system, ATP being the precursor of cAMP. The resulting *E vs t* profile is a horizontal exponential curve (see Fig. 4B of (Pohl et al., 1971)), consistent with the kinetic model.

#### 3.2.4. Mechanism-agnostic analysis of horizontal exponential time course data for estimating agonist efficacy and affinity

Time course data from all of the horizontal exponential models considered here can be analyzed the same way to determine agonist efficacy for generating the response, i.e. *k_τ_*. This is done by multiplying the Plateau by the rate constant of the horizontal exponential fit for a maximally-stimulating concentration of agonist. The theoretical basis of this approach is evident from the general form of all the model equations (Eqs. 5, 8, 11 and 14):

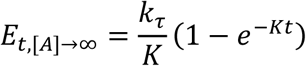

where K is the observed rate constant. From this equation, it is evident that *k_τ_* can be determined by multiplying the Plateau by the rate constant:

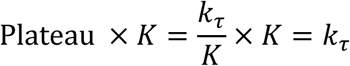

This means that agonist efficacy (*k_τ_*) can be determined without knowing the regulatory mechanism underlying the horizontal exponential profile, assuming the profile results from one of the models described in this study. This analysis method can be described as agnostic regarding the regulatory mechanism. The observed rate constant K can be described as the response regulation rate constant, here termed *k_REG_*. This rate constant governs the time it takes for the response to level off.

This agnostic approach is used here to re-analyze historical data on agonist activity at the μ-opioid receptor, from the study of (Traynor et al., 2002). In this study, the time course data for the response, [^35^S]-GTPγS binding, conform to a horizontal exponential curve and the data were analyzed in the original study using the generic equation to determine the plateau and the rate constant (termed *B*_max_ and *k*, respectively, in the original article) (Traynor et al., 2002). The parameter values from the study are shown in Table 1. From these values, *k_τ_* can be calculated, by multiplying *B*_max_ by *k*. The *k_τ_* values indicate a broad range of agonist activity, spanning 20-fold, from 0.0024 fraction.min^−1^ for nalbuphine to 0.048 fraction.min^−1^ for DAMGO. The results are consistent with the literature, since: 1) Butorphanol and nalbuphine, partial agonists in terms of the *k_τ_* value, are partial agonists in other assays of μ opioid receptor activity (Hoskin and Hanks, 1991). 2) Morphine has a slightly lower *k_τ_* value than DAMGO (Table 1), and has slightly lower E_max_ than DAMGO in [^35^S]-GTPγS binding assays (Traynor and Nahorski, 1995).

**Table 1.** Mechanism-agnostic analysis of μ opioid receptor-stimulated [^35^S]-GTP S binding

| Ligand | $k$<br>(min<sup>-1</sup>) | $B_{\max}$<br>(fraction DAMGO) | $k_{\tau}$<br>(fraction.min<sup>-1</sup>) | $k_{\tau}$<br>(% DAMGO) |
| --- | --- | --- | --- | --- |
| DAMGO | 0.048 | 1.00 | 0.048 | 100 |
| Morphine | 0.039 | 0.84 | 0.033 | 68 |
| Meperidine | 0.036 | 0.80 | 0.029 | 60 |
| Butorphanol | 0.023 | 0.22 | 0.0051 | 11 |
| Nalbuphine | 0.010 | 0.24 | 0.0024 | 4.9 |

Agonist affinity can also be determined using the agnostic approach. This analysis assumes agonist rapidly equilibrates with the receptor. [If it does not, the *E vs t* profile deviates progressively from a horizontal exponential curve as the agonist concentration is decreased (Supplemental Fig. S2).] This analysis is based on the common general form of all the horizontal exponential model equations for non-saturating concentrations of agonist (Eqs. 4, 7, 10 and 13):

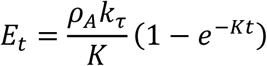

Multiplying the Plateau by the rate constant gives *ρ*_A_*k_τ_*:

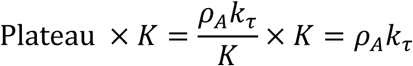

This equation can be expanded to include the agonist concentration as the dependent variable:

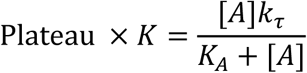

A familiar hyperbola results from plotting Plateau × *K* versus the agonist concentration. The midpoint of the curve, i.e. [*A*] at half-maximal Plateau × *K*, is the agonist affinity, *K_A_*. This type of analysis is typically done using the logarithm of [*A*], employing a four-parameter logistic equation:

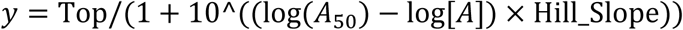

This can be done in Prism using the “log(agonist) vs. response -- Variable slope (four parameters)” equation, with the parameter “Bottom” set to zero (Motulsky, 2019b). The *A_50_* value is *K_A_* and the *y* value at maximally-effective agonist concentrations (“Top” in the equation above) is *k_τ_*. The Hill slope parameter accommodates effects that deviate the concentration range exposed to the receptor from the concentration range added to the assay, for example precision of serial dilution or biological processes that affect local concentration in the vicinity of the receptor. The Hill slope can also allow cooperativity of agonist binding to be incorporated.

### 3.3. Two regulation mechanisms: Rise-and-fall exponential profiles

In unmodified response systems, GPCR signaling is typically regulated at both levels of the regulation system, the response generation process (receptor desensitization or, in the case of calcium signaling, precursor depletion) and the response degradation process. This scenario results in a rise-and-fall time course profile, commonly observed for unmodified GPCR responses, as shown below. The *k_τ_* value can be determined by fitting time course data to a generic equation and this can be done without knowing the mechanism that regulates the response generation process (Section 3.3.3). The mechanisms that result in this profile are presented below. They incorporate response degradation with a second regulation process operating on the response input, either receptor desensitization or precursor depletion.

#### 3.3.1. Receptor desensitization and response degradation (Model 6)

First, the model with receptor desensitization and response degradation is considered. The model combines the yellow, green and pink regions of Fig. 1 and is shown schematically in Scheme 6 below:

**Scheme 6.**
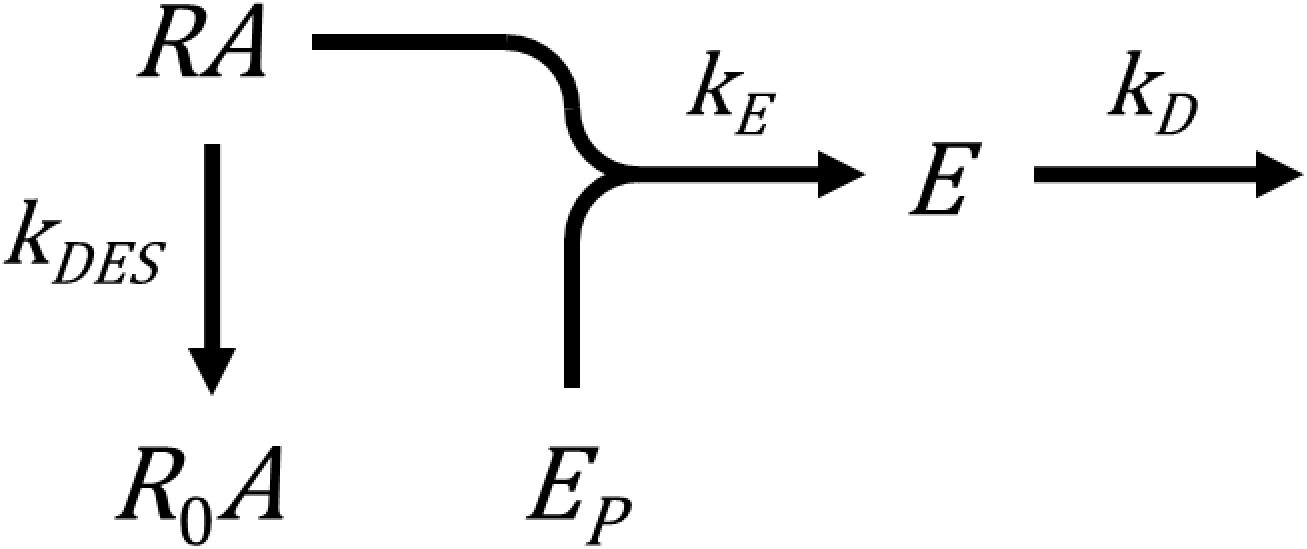

This model predicts a rise-and-fall to zero curve (Fig. 2D). This makes sense intuitively. The response rises rapidly then slows as receptor desensitization starts to slow the rate of response generation. The response becomes further limited owing to response degradation. The response reaches an upper limit (the peak) when the rate of response generation equals the rate of degradation. After this, response degradation predominates over response generation. Less and less new response is generated because the active receptor concentration is declining, and ultimately no new response will be generated because the active receptor concentration will decline to zero. The response level declines because it is being degraded and ultimately the response level falls to zero once all existing response has been degraded.

The equation defining the response over time is, for a maximally-stimulating concentration of agonist, Eq. 6 (derived in Appendix A.6):

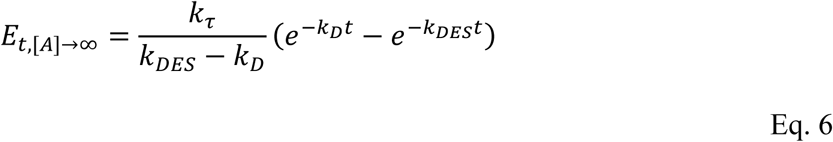

This equation is of the same form as a generic rise-and-fall to zero exponential equation, familiar in pharmacokinetics as the equation defining drug concentration over time after oral dosing:

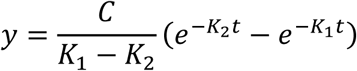

The model was validated by comparison with experimental data. Fig. 6 shows data for the IP_3_ response to cholecystokinin (CCK) via the CCK_1_ receptor in pancreatic acinar cells (Streb et al., 1985). The mechanism of this response conforms to Scheme 6: It is regulated by response degradation [shown by sensitivity to Li^+^, Fig. 6, (Streb et al., 1985)] and by receptor desensitization (Klueppelberg et al., 1991). The shape of the time course is a rise-and-fall to zero exponential curve (solid squares, Fig. 6), consistent with the kinetic model. A second example is diacylglycerol signaling by the AT_1_ angiotensin receptor (Violin et al., 2006) (Fig. 7). This response is modulated by receptor desensitization; it is sensitive to removal of phosphorylation sites from the receptor (Violin et al., 2006) (Fig. 7). It is presumably also modulated by response degradation because diacylglycerol is cleared by diacylglycerol kinases (Merida et al., 2008). The shape of the time course is a rise-and-fall to zero exponential curve (Fig. 7, sold squares), consistent with the kinetic model.

**Fig. 6.**
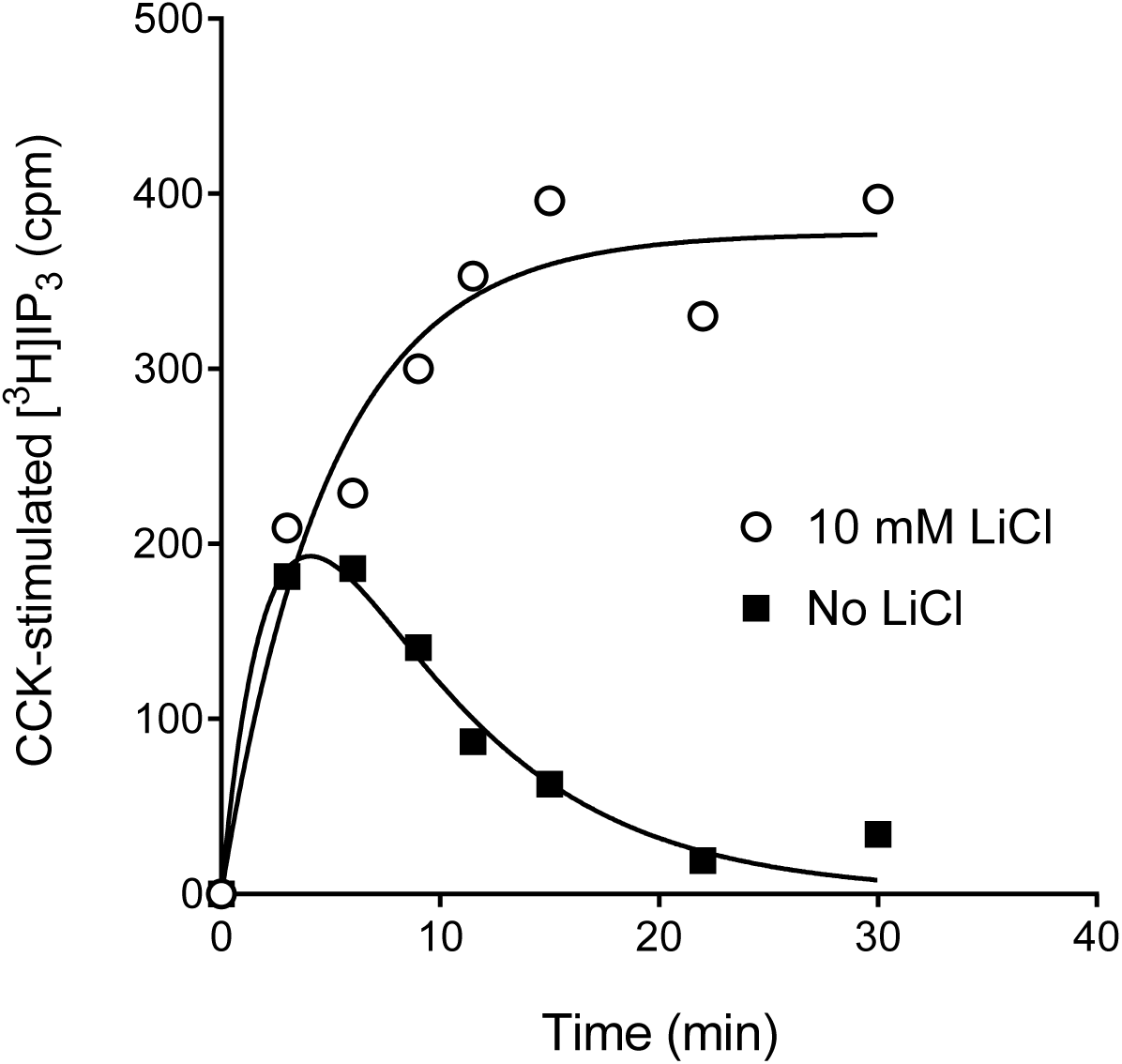
Receptor desensitization with response degradation: CCK-stimulated IP_3_ accumulation in pancreatic acinar cells (Streb et al., 1985). In these cells, the CCK_1_ receptor becomes desensitized on application of agonist (Klueppelberg et al., 1991) and IP_3_ is degraded. This results in a rise-and-fall exponential curve (Section 3.3.1). Data were fit to Model 6, the receptor desensitization and response degradation model (see Section 3.3.1 for details), giving the following fitted values: *k_τ_*, 130 cpm [^3^H]IP_3_.min^−1^; *k_DES_*, 0.14 min^−1^; *k_D_*, 0.39 min^−1^. In the presence of Li^+^, the *E vs t* profile becomes a horizontal exponential curve, consistent with suppressed response degradation and with the remaining regulation process being receptor desensitization (Section 3.2.1). Data for the presence of Li^+^ (i.e. desensitization alone) were fit to Model 2 (Section 3.2.1), giving fitted values of: *k_τ_*, 75 cpm [^3^H]IP_3_.min^−1^; *k_DES_*, 0.20 min^−1^. Data are from Fig. 10 of (Streb et al., 1985), extracted using a plot digitizer [WebPlotDigitizer (Rohatgi, 2018)].

**Fig 7.**
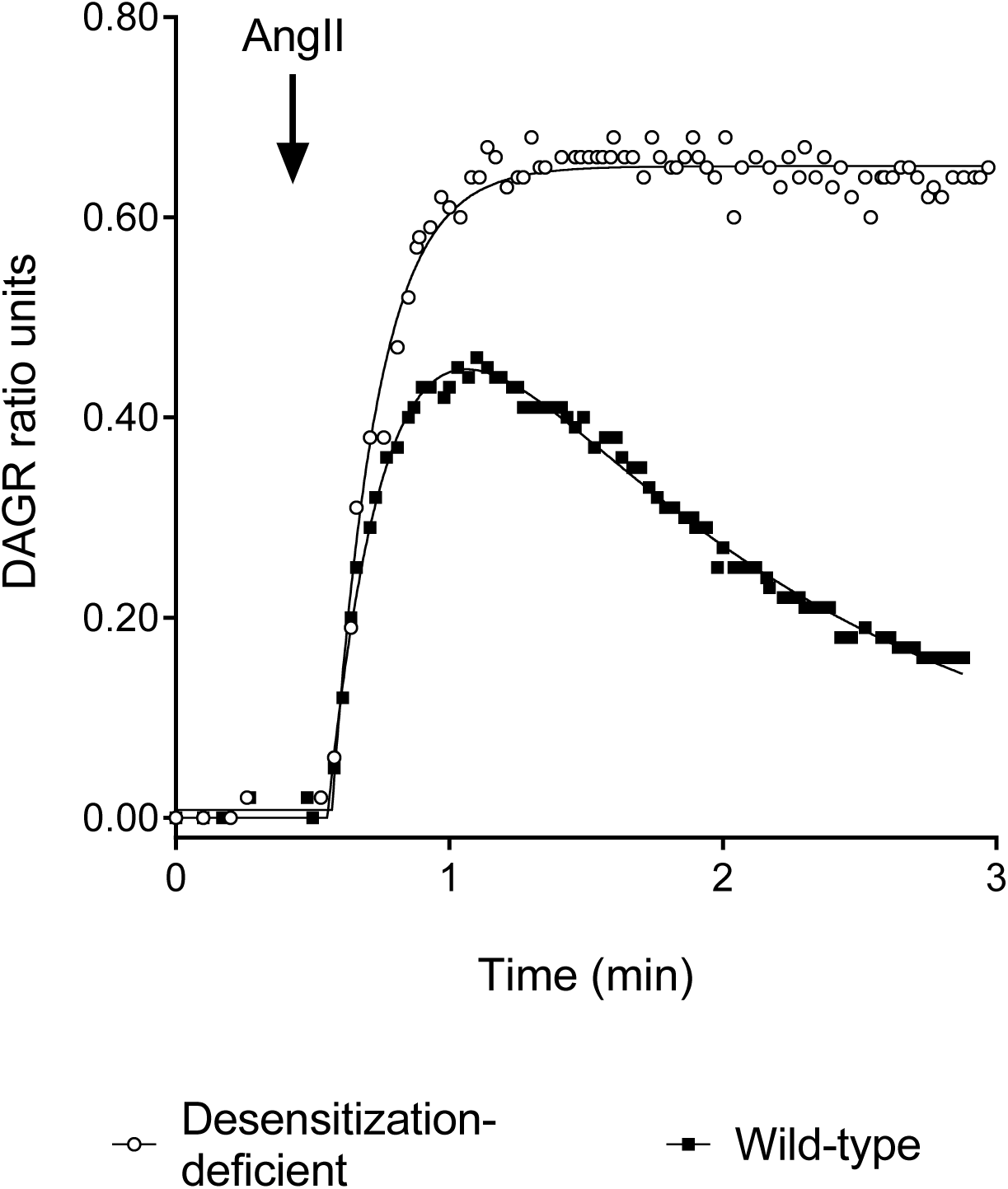
Receptor desensitization with response degradation: Angiotensin II-stimulated diacylglycerol generation via the AT_1_ angiotensin receptor in HEK-293 cells (Violin et al., 2006). Diacylglycerol was quantified using a live-cell biosensor. Diacylglycerol is degraded by diacylglycerol kinases and the AT_1_ receptor is susceptible to desensitization. This results in a rise-and-fall exponential curve (Section 3.3.1.). Data for the wild-type receptor were fit to Model 6, the receptor desensitization and response degradation model (see Section 3.3.1 for details), giving the following fitted values: *k_τ_*, 2.7 DAGR ratio units.min^−1^; *k_DES_*, 0.74 min^−1^; *k_D_*, 4.1 min^−1^. Receptor desensitization can be suppressed using a modified AT_1_ receptor which lacks phosphorylation sites (“Desensitization deficient,” open circles). The *E vs t* profile becomes a horizontal exponential curve, consistent with suppressed receptor desensitization and with the remaining regulation process being response degradation (Section 3.2.2). Data for the desensitization-deficient receptor (i.e. degradation alone) were fit to Model 3 (Section 3.2.2), giving fitted values of: *k_τ_*, 4.0 DAGR ratio units.min^−1^; *k_DES_*, 6.1 min^−1^. Data are from Fig. 2A of (Violin et al., 2006), extracted using a plot digitizer [WebPlotDigitizer (Rohatgi, 2018)].

The data in Fig. 6 and 7 can be analyzed using the model to determine agonist efficacy for generating the response, in other words the *k_τ_* value. A simple way to do this is to fit the data to the generic rise-and-fall to zero equation:

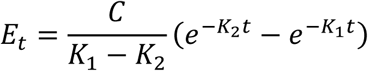

By comparison with Eq. 6, it is evident that *C* is equal to *k_τ_*. This method is applied to the experimental data in Figs. 6 and 7. (Note the analysis assumes a maximally-stimulating agonist concentration, employed in these experiments.). A Prism template is provided in the Supplementary material in which this equation, termed “[Pharmechanics] Rise-and-fall to zero time course” is pre-loaded. In this equation, *K*_1_ is constrained to be greater than *K*_2_.

The fitted *k_τ_* value for CCK-stimulated IP_3_ accumulation was 130 cpm [^3^H]IP_3_.min^−1^ and the *k_τ_* value for angiotensin-stimulated diacylglycerol production was 2.7 DAGR ratio units.min^−1^. The fit also provides estimates of the rate constants *K*_1_ and *K*_2_. For the CCK_1_ response these values were 0.39 and 0.14 min^−1^, respectively, and for the AT_1_ response were 4.1 and 0.74 min^−1^, respectively. The question arises as to which generic rate constants (*K*_1_ or *K*_2_) correspond to which model rate constants (*k_DES_* or *k_D_*). Unfortunately, it is not possible to assign the model rate constants using this equation. This is because *K*_1_ and *K*_2_ are interchangeable, shown by the following relationship:

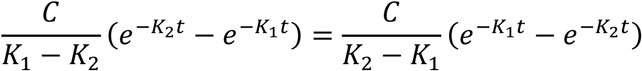

One way to assign the rate (*K*_1_ or *K*_2_) to the process (receptor desensitization or response degradation) is to manipulate the process. In the CCK_1_ response, response degradation was manipulated by blocking it with Li^+^. The resulting response is a horizontal exponential curve, expected for a response regulated solely by receptor desensitization (Section 3.2.1). *k_DES_* can be estimated by fitting these data to the horizontal exponential equation (Section 3.2.1). The resulting *k_DES_* value was 0.20 min^−1^. This was close to the *K*_2_ value for the response not treated with Li^+^ (0.14 min^−1^). Therefore, in this case, *K*_2_ was ascribed to *k_DES_*. This means, by a process of elimination, *K*_1_ corresponds to *k_D_*. Consequently, *k_D_* was estimated to be 0.39 min^−1^. In simple terms, this means the half-life for CCK_1_-receptor desensitization was 5.0 min and the half-life for IP_3_ degradation was 1.8 min.

The same approach was applied to the AT_1_ diacylglycerol response. In this case, receptor desensitization was blocked using a phosphorylation-deficient receptor (Fig. 7). This results in a horizontal exponential curve (Fig. 7), as anticipated for a response regulated solely by response degradation (Section 3.2.2.). Analysing these data as described in Section 3.2.2. gave a *k_D_* value of 6.1 min^−1^. This value was close to that of *K*_1_ of the wild-type response (4.1 min^−1^) so *K*_1_ was ascribed to *k_D_*. As a result, by a process of elimination *K*_2_ was assigned to *k_DES_*, giving a *k_DES_* value of 0.74 min^−1^. From this analysis it is concluded the half-life for AT_1_ receptor desensitization in this system was 0.94 min and the half-life of diacylglycerol clearance was 0.17 min.

#### 3.3.2. Precursor depletion and response degradation – calcium signaling (Model 8)

In this model, the response precursor concentration is depleted by being converted to the response, and the response, once generated, is completely cleared. The model has been presented previously (Hoare et al., 2018). It is represented by Scheme 8 below:

**Scheme 8.**
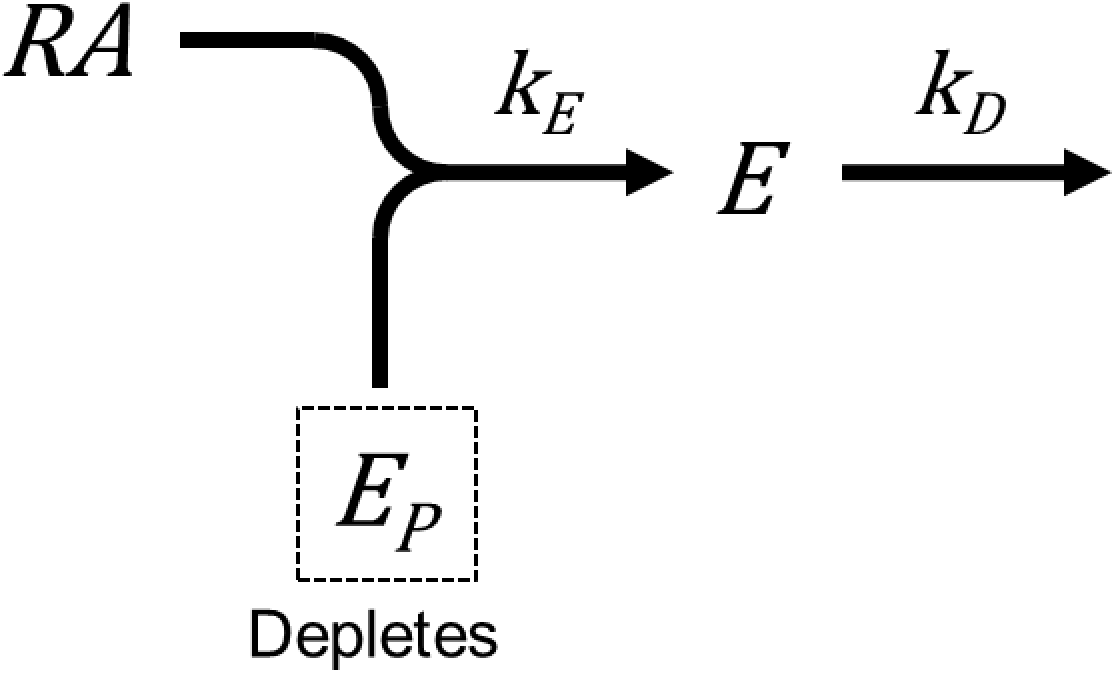

This model predicts a rise-and-fall to zero curve (Fig 8). This makes sense intuitively. The response rises rapidly at first as response generation predominates. Then response generation becomes limited because the precursor becomes depleted, and response is further limited because it degrades. A peak is reached at which the rate of response generation equals the rate of response degradation. Ultimately all the accessible precursor becomes depleted so no new response is generated. The response ultimately declines to zero because the existing response is completely degraded and no new response is generated because the precursor is depleted.

**Fig. 8.**
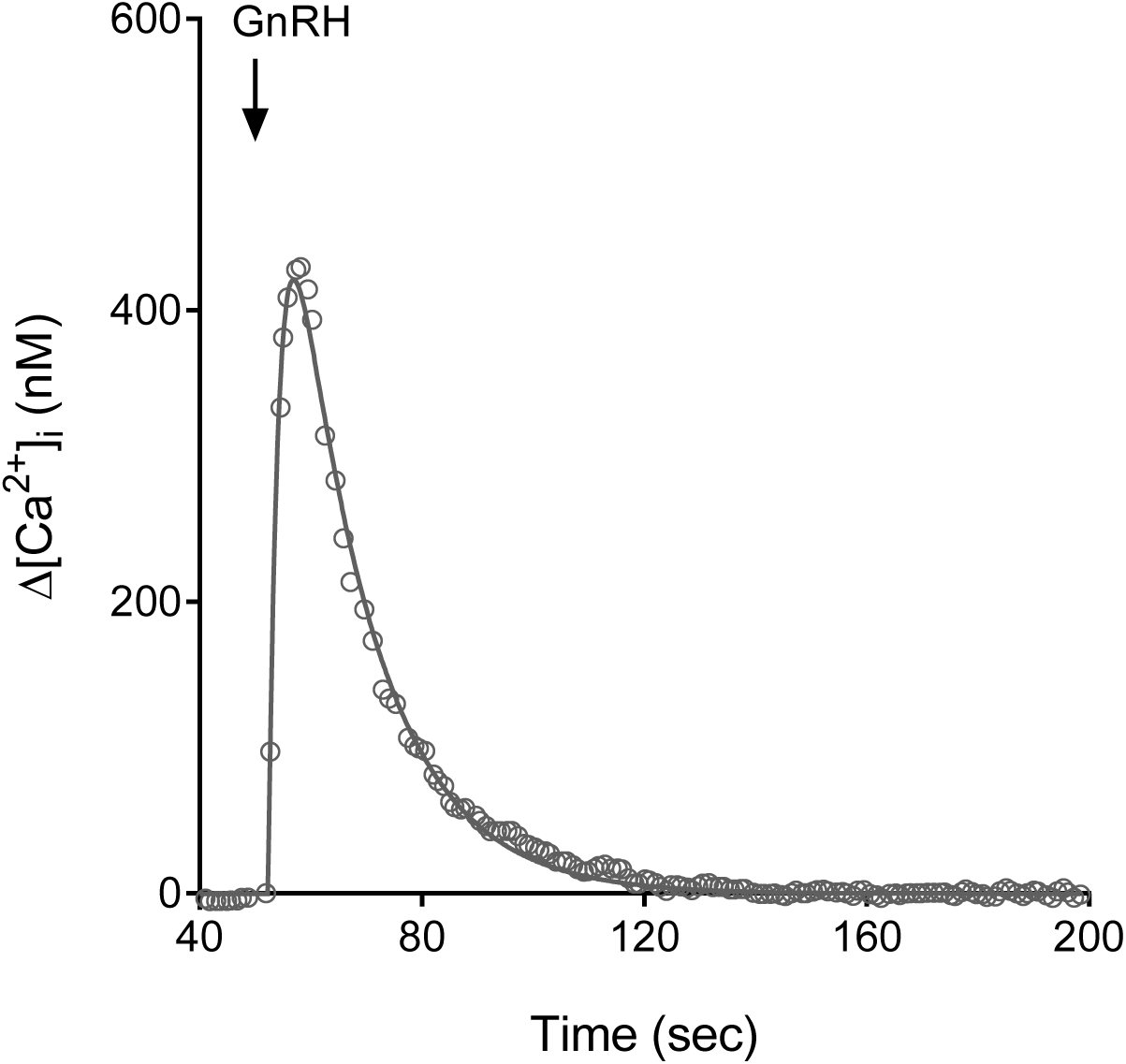
Precursor depletion and response degradation: GnRH-stimulated Ca^2+^ mobilization in pituitary gonadotrophs. Response precursor is Ca^2+^ in the ER and the response is Ca^2+^ released into the cytoplasm on stimulation by GnRH. Precursor is depleted because the calcium store becomes emptied. Response is degraded because cytoplasmic Ca^2+^ is exported out of the cell. (See section 3.3.2). Data were fit to the precursor depletion and response degradation model (section 3.3.2), giving fitted values of: *k_τ_*, 290 nM.sec-1; *k_DEP_*, 0.49 sec-1; *k_D_*, 0.071 sec-1. Data are from Fig. 1A (absence of extracellular Ca^2+^) of (Merelli et al., 1992), extracted using a plot digitizer (Graph Grabber 2.0, Quintessa Ltd., Henley-on-Thames, United Kingdom).

This model can describe calcium signaling via GPCRs in certain circumstances, specifically Ca^2+^ entry into the cytoplasm from the endoplasmic reticulum (ER). The response precursor is Ca^2+^ stored in the ER. Response generation is the increase of cytoplasmic Ca^2+^ concentration resulting from opening of the IP_3_-gated Ca^2+^ channel in the ER membrane (Berridge, 1993; Berridge et al., 2003). Precursor depletion is the decrease of Ca^2+^ in the ER resulting from Ca^2+^ release into the cytoplasm (Hofer et al., 1998; Miyawaki et al., 1997; Yu and Hinkle, 1997; Yu and Hinkle, 2000). Response degradation is the clearance of cytoplasmic Ca^2+^ into the extracellular space by the plasma membrane Ca^2+^ pump (Carafoli, 1991; Hofer et al., 1998). Receptor desensitization likely does not greatly impact the temporal profile of a single Ca^2+^ response for rapid Ca^2+^ responses; receptor desensitization proceeds over minutes, whereas a rapid Ca^2+^ response peaks in a few seconds and is largely cleared within a minute (exemplified in Fig. 8). (Receptor desensitization likely operates to limit the Ca^2+^ response on a second application of the agonist, or to limit sustained Ca^2+^ responses.) The model does not accommodate the refilling of the ER with Ca^2^ (which would represent response recycling). ER refilling can be blocked by removing extracellular calcium, since the ER is refilled primarily by Ca^2+^ that enters into the cell (Hofer et al., 1998; Tsien and Tsien, 1990; Yu and Hinkle, 2000).

The equation defining the response over time is, for a maximally-stimulating concentration of agonist, Eq. 8, derived in Appendix A.8:

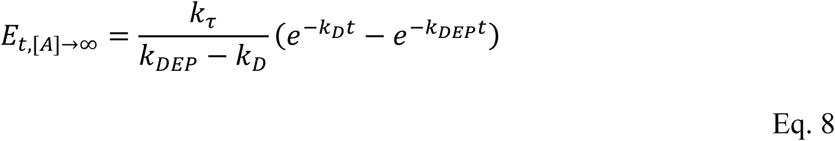

This equation introduces a parameter that represents the rate of precursor depletion. This parameter is the depletion rate constant, termed *k_DEP_*. It is defined as the product of the receptor concentration and the response generation rate constant:

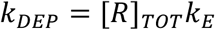

The half-life of response precursor is determined from this parameter, as 0.693 / *k_DEP_*.

The model was applied to experimental Ca^2+^ mobilization data to estimate agonist efficacy and the rate constants. This was done using data for the GnRH_1_ receptor (Merelli et al., 1992). [In this case, we can assume the response is unaffected by receptor desensitization because this receptor does not undergo short-term desensitization (Section 3.2.1).] This can be done by fitting the data to the generic rise-and-fall equation introduced in Section 3.3.1:

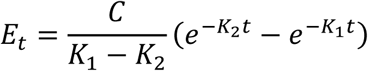

Agonist efficacy (*k_τ_*) can be determined from this fit – it is equal to *C*. Fitting the equation to Ca^2+^mobilized by the GnRH_1_ receptor in α3T-1 clonal pituitary gonadotrophs (Merelli et al., 1992) gives a *k_τ_* value of 290 nM.sec^−1^ (Fig 8). The fit also provides estimates of *K*_1_ and *K*_2_. In principle it isn’t possible to determine which of these parameters can be ascribed to which of the model rate constants (*k_DEP_* or *k_D_*) (see Section 3.3.1.). However, in practice it is likely that the faster of the two fitted rate values is *k_DEP_*, because this is the rate of emptying of the Ca^2+^ store, which presumably proceeds more rapidly than clearance of Ca^2+^ from the cytoplasm (defined by *k_D_*). Applying this logic to the fit to the GnRH data gives a *k_DEP_* value of 0.49 sec^−1^ and *k_D_* value of 0.071 sec^−1^ (Fig. 8), corresponding to half-times for store emptying and cytoplasmic clearance of 1.4 sec and 9.8 sec, respectively.

#### 3.3.3. Mechanism-agnostic analysis of rise-and-fall time course data for estimating agonist efficacy and affinity

Data from the rise-and-fall mechanisms in this study can be analyzed using a generic equation to estimate agonist efficacy (*k_τ_*). When using a maximally-stimulating concentration of agonist, the data are fit to the equation below and the fitted parameter *C* is equal to *k_τ_*:

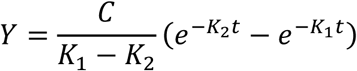

A Prism template is provided in the Supplementary material in which this equation, termed “[Pharmechanics] Rise-and-fall to zero time course” is pre-loaded.

Agonist affinity, *K_A_*, can also be determined using this agnostic analysis if the agonist equilibrates rapidly with the receptor. [If it does not, the *E vs t* profile deviates progressively from a horizontal exponential curve as the agonist concentration is decreased (Supplemental Fig. S2).] For non-saturating concentrations of agonist the equations conform to the following general equation:

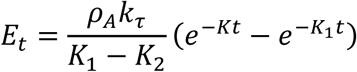

This equation is of the same form as the generic rise-and-fall equation above where *ρ*_A_*k*_*τ*_ is equal to *C*. Expanding *ρ*_A_ gives an equation including *K_A_*:

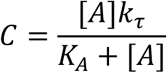

This is an equation for a hyperbola when [*A*] is the dependent variable. Plotting *C* against [*A*] gives a hyperbola where the half-maximally-effective concentration is *K_A_* and the asymptote is *k_τ_*. The analysis can be done using a four parameter-logistic equation as described in Section 3.2.4.

### 3.4. Other mechanisms

More complex mechanisms involve multiple regulation processes acting in concert. These mechanisms result in equations that are more complex than for the mechanisms described thus far.

#### 3.4.1. Receptor desensitization and response degradation with recycling (Model 7)

This model is an extension of Model 6. The receptor desensitizes and the response degrades but in this case the response degrades back to the response precursor, i.e. it recycles. The model is represented by Scheme 7 below:

**Scheme 7.**
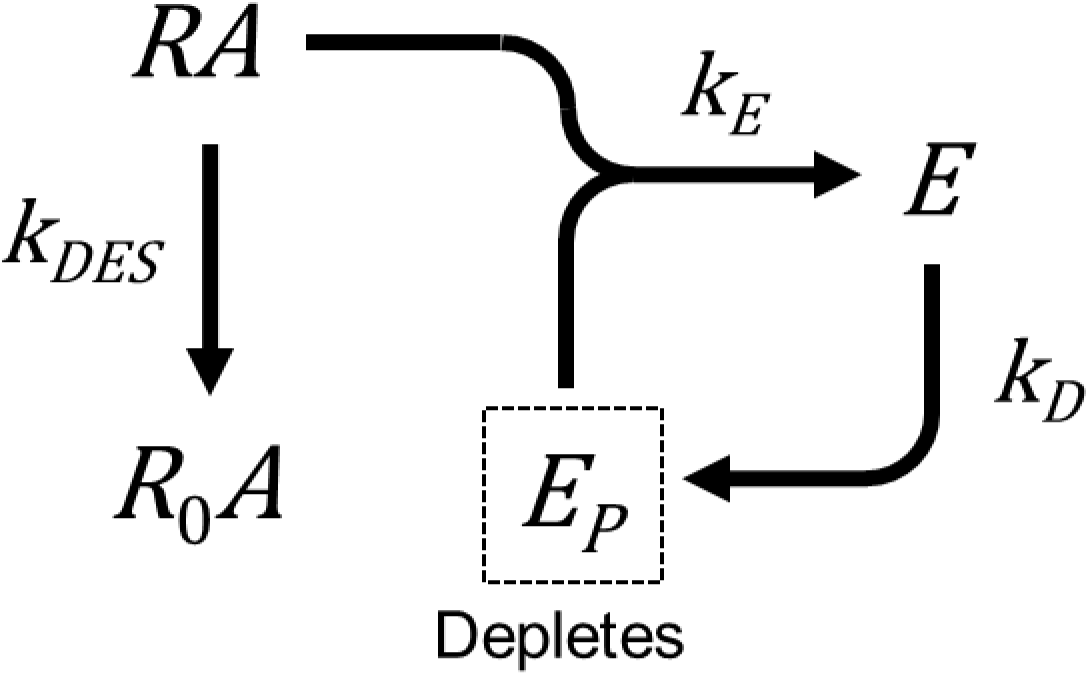

An *E vs t* equation could not be found for this model (see Appendix A.7. for details). The *E vs t* profile was simulated by numerical solution of the differential equations and is shown in Fig. 9A. The profile is a modified form of the rise-and-fall profile in which there is a lag in the decline phase. This results in a bulge on the right-hand, downward part of the curve (Fig. 9A). This results because the recycling regenerates the precursor, delaying the decline of the response that results from degradation of the response.

**Fig. 9.**
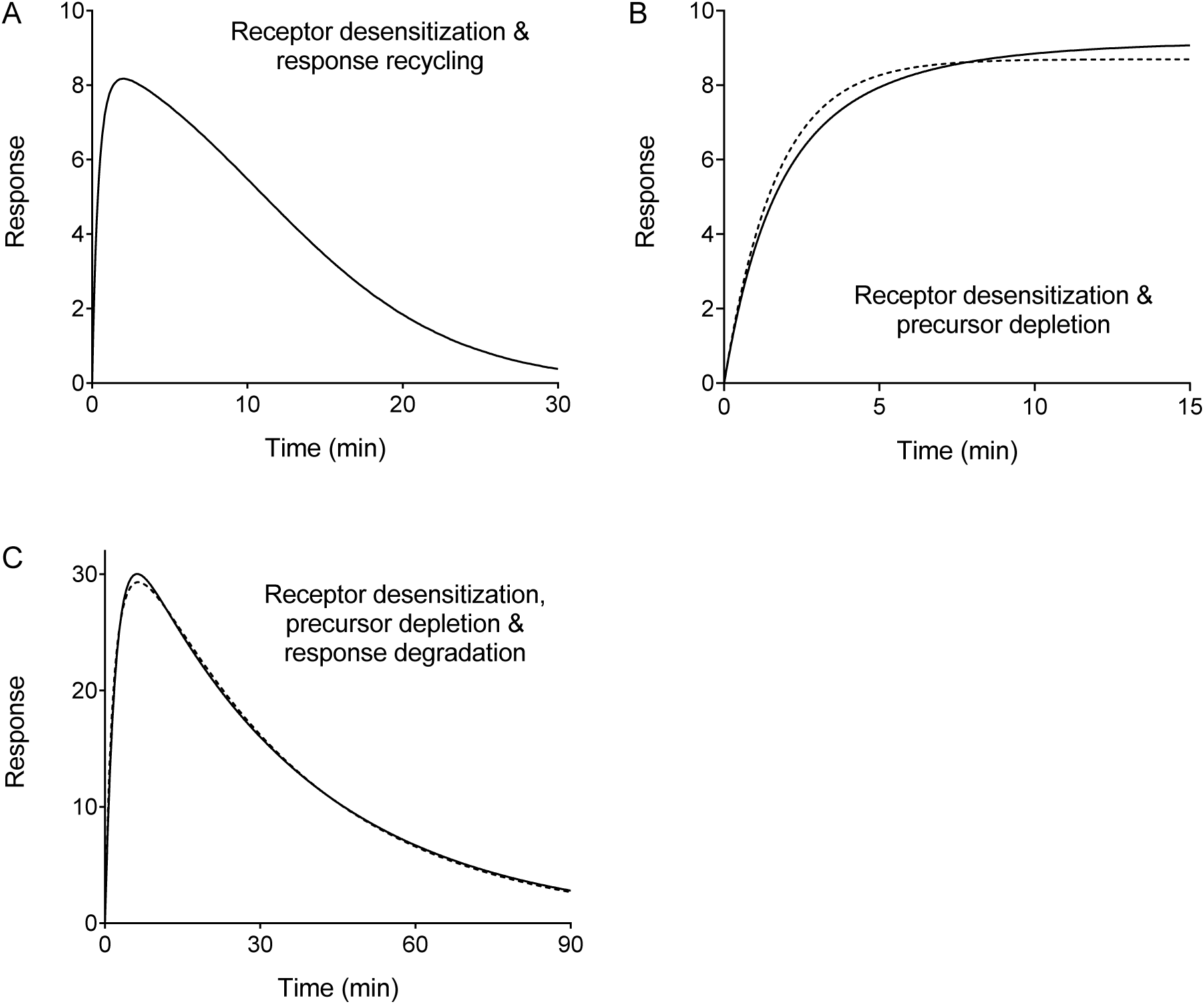
Simulated data for more complex kinetic response models. Multiple regulation mechanisms acting in concert result in more complex time course equations (see Section 3.4). (a) Receptor desensitization and response recycling (Model 7). Data were simulated numerically using the differential equations in Appendix A.7., with *ρ_A_* set to unity (representing a maximally-stimulating concentration of agonist) with the following parameter values: *k_τ_*, 20 response units.min^−1^; *E_P_*_(*TOT*)_, 10 response units; *k_DES_*, 0.20 min^−1^; *k_D_*, 0.30 min^−1^. (b) Receptor desensitization and precursor depletion (Model 9). The *E vs t* profile is a Gompertz curve, which ascends more slowly than a horizontal exponential curve at later time points (Kirkwood, 2015). A horizontal exponential curve is shown for comparison (dashed line). Data were simulated using Eq. 9 with the following parameter values: *k_τ_*, 5.0 response units.min^−1^; *E_P_*_(*TOT*)_, 10 response units; *k_DES_*, 0.20 min^−1^. (c) Receptor desensitization, precursor depletion and response degradation (Model 10). Data were simulated numerically using the differential equations in Appendix A.10, with *ρ_A_* set to unity (representing a maximally-stimulating concentration of agonist) with the following parameter values: *k_τ_*, 20 response units.min^−1^; *E_P_*_(*TOT*)_, 40 response units; *k_DES_*, 0.20 min^−1^; *k_D_*, 0.030 min^−1^. The dashed line is a fit of the simulated data to the generic rise-and-fall exponential equation (Section 3.3.4.), giving a *k_τ_* value of 17 response units.min^−1^, *K*_1_ of 0.48 min^−1^ and *K*_2_ of 0.029 min^−1^.

#### 3.4.2. Receptor desensitization and response precursor depletion (Model 9)

In this model, the receptor desensitizes and the precursor is depleted as it is being converted to the response. The model is represented by Scheme 9:

**Scheme 9.**
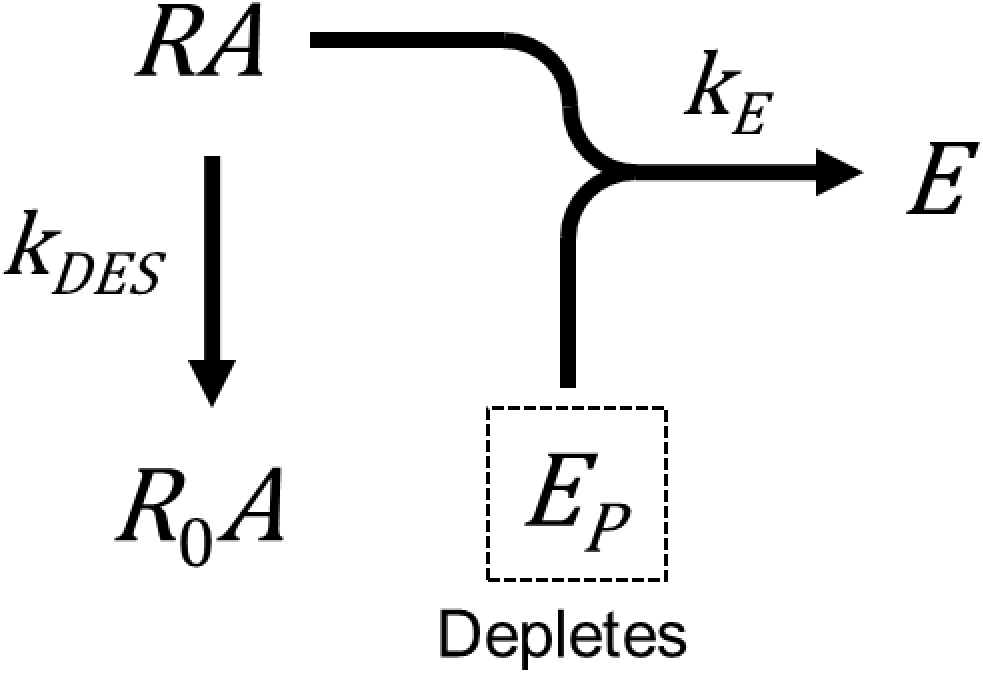

This model is included in part to demonstrate that analytical solutions can be obtained even though the system of differential equations is nonlinear. [Another example is the slow agonist equilibration variant of Model 5 (Appendix A.5.).] This is because the equation defining E contains the product of two time-dependent terms, *E_P_* and [*RA*]:

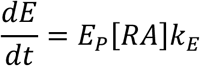

The derivation method is shown in Appendix A.9. and the *E vs t* equation for a maximally-stimulating concentration of agonist is Eq. 9:

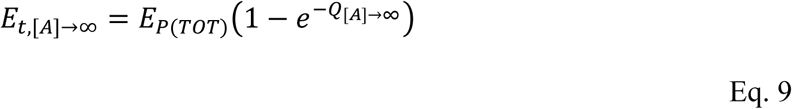

where,

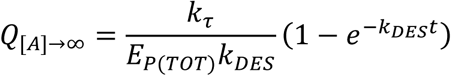

This is the equation for a Gompertz curve, originally derived to describe human mortality and then applied generally to modeling population biology (Kirkwood, 2015). It resembles the horizontal exponential equation but the resulting curve ascends more slowly at the later time points. This is shown in the simulated *E vs t* data in Fig. 9B.

#### 3.4.3. Receptor desensitization, precursor depletion and response degradation (Model 10)

In this model all three mechanisms are in operation. The model is represented by Scheme 10 below:

**Scheme 10.**
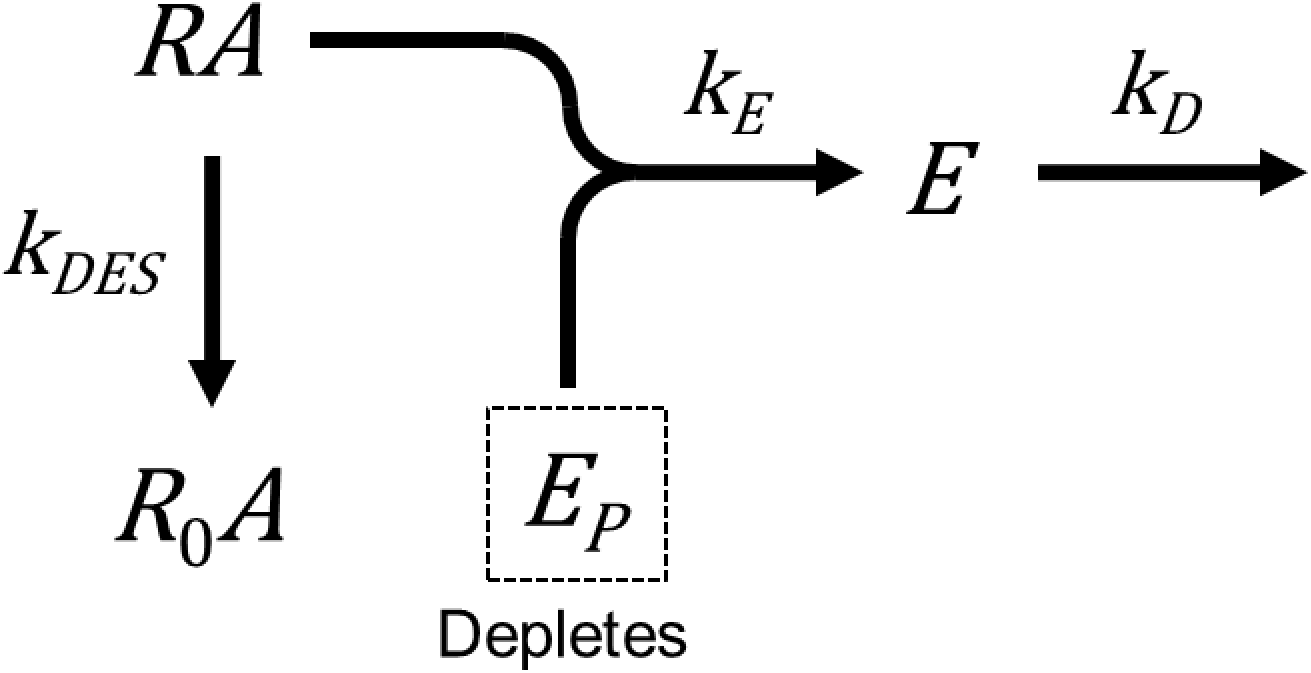

An analytical equation for response could not be found [although a solution for response precursor could be found (Eq. A.14, Appendix A.10, Fig. S3B)]. *E vs t* data were simulated by numerical solution of the differential equations (Appendix A.10.). A rise-and-fall type profile results (Fig. 9C). It is evident by visual comparison that this profile appears similar in shape to rise-and-fall profiles fitted by the generic rise-and-fall equation, for example Fig 2D). Even though mathematically the model does not conform to the generic rise-and-fall equation, simulated data generated using the model were very well, although not precisely, fit by the generic equation (Fig. 9C). This makes sense intuitively because the effects balance out. The more depletion occurs, the less influence receptor desensitization has on limiting the response, and vice-versa. Conceptually the model contains two macroscopic regulation processes – regulation of the input that generates the response (receptor desensitization and precursor depletion), and regulation of the output (response degradation). We ran data simulations with a diverse array of model parameter values for a maximally-stimulating concentration of agonist and fit the data to the generic rise-and-fall equation in Section 3.3.3. In all cases the fitted *k_τ_* value was within 15% of the simulation value (data not shown).

## 4.0 Discussion

In this study a kinetic pharmacological model of GPCR signaling is presented that incorporates regulation of signaling. Regulation mechanisms control the magnitude of the signal over time and so impact the measurement of pharmacological activity of agonist ligands. The model incorporates the two canonical mechanisms of short-term regulation, receptor desensitization and response degradation, providing a comprehensive model that can be broadly applied to signaling assays used in drug discovery. The model was verified using historical data from studies in which the regulation processes were carefully manipulated. Meaningful drug parameters were obtained using familiar curve fitting methods applied to time course data. The parameters quantify the initial rate of signal generation by the agonist (*k_τ_*, a measure of efficacy) and the rate(s) of signal regulation. These parameters are designed for determining structure-activity relationships of agonist efficacy and signaling regulation, especially in the lead-optimization stage of drug discovery.

The drug efficacy term that emerges from the model is *k_τ_* (Hoare et al., 2018). This is simply the initial rate of signaling stimulated by the agonist, defined as the product of the precursor concentration, receptor concentration and a rate constant (Section 2.2.1). As an efficacy parameter, *k_τ_* has a number of benefits. It is biochemically meaningful and, being analogous to the initial rate of enzyme activity, conceptually familiar to most investigators. From a pharmacological theory perspective, *k_τ_* has the benefit of being independent of receptor reserve, as described previously (Hoare et al., 2018). Consequently, it might be a better predictor in in vivo efficacy than efficacy measurements that are dependent on receptor reserve. *k_τ_* is more straightforward to measure than other efficacy parameters that take into account receptor reserve. For example, measuring *τ* of the operational model usually requires system manipulations and/or grouped analyses (Black and Leff, 1983; Kenakin et al., 2012; Klein Herenbrink et al., 2016; Slack and Hall, 2012; Van der Graaf and Stam, 1999; Zernig et al., 1996). All that is required for measuring *k_τ_* is a time course for a maximally-stimulating concentration of agonist (see below).

The regulation parameters in the model are the rate constants for receptor desensitization (*k_DES_*) and response degradation (*k_D_*) (Section 2.2.2). These parameters enable regulation of signaling to be quantified. Measuring *k_DES_* enables ligands to be compared in terms of their receptor desensitization activity. Measuring *k_DES_* enables response degradation activity to be compared across different response systems. These parameters are separate from the efficacy parameter *k_τ_* (Section 2.2.2). This means that in SAR campaigns, regulation activity can be optimized separately from ligand efficacy. This capability could facilitate optimization for the duration of efficacy, separate from the magnitude of efficacy (defined by *k_τ_*). This is not possible in single time-point pharmacological models, such as the operational model. At a single time point, the measured response is a combination of response generation (efficacy) and response regulation (assuming the response is regulated at that time point) (Riccobene et al., 1999). This can result in time-dependence of single time-point efficacy parameters, such as *τ* in the operational model, an effect observed experimentally (Klein Herenbrink et al., 2016). This effect might contribute to the time-dependence of biased agonism, especially if the regulation kinetics of the pathways being compared are different. In principle, the kinetic model could circumvent these complications.

The model parameters are reasonably straightforward to measure. The time course for a maximally-stimulating concentration of agonist is fitted to a generic equation, familiar in most cases to drug discovery pharmacologists. The fitted generic parameters are then combined or used directly to calculate *k_τ_* (Sections 3.2.4 and 3.3.3). For example, for a horizontal exponential fit, the plateau is multiplied by the rate constant, giving *k_τ_* (Section 3.2.4). The fit also provides an estimate of the regulation rate constant(s), for example *k_DES_* and *k_D_*. For the horizontal exponential fit, this value is simply the fitted value of the rate constant. Remarkably, it is not necessary to know the regulatory mechanism in order to determine *k_τ_*, providing the data conform to a horizontal exponential or the rise-and-fall curve. This means *k_τ_* can be determined using a mechanism-agnostic analysis; the data are fit to the generic equation and *k_τ_* determined from the fitted generic parameters. These fits also provide an agnostic regulation parameter, *k_REG_*. This parameter defines the rate at which the signal is dampened by the regulatory process. If the regulatory mechanism is subsequently determined, the generic rate constants can be ascribed to *k_DES_*, *k_D_* or *k_DEP_*.

The notable limitation of the model is the technical difficulty of measuring equilibrium binding affinity of the agonist for the receptor. It cannot be generally assumed that agonist binding is at equilibrium with the receptor throughout the time course of response measurement. It can take time for the concentration of the agonist-receptor complex to rise and closely approach its equilibrium value (Section 2.2.3). While likely not an issue for low potency ligands (affinity in the μM range), lack of equilibration is likely to affect the time course of response for high affinity ligands, leading to erroneous estimates of ligand affinity. This problem is exacerbated for rapid responses, such as calcium mobilization (Bdioui et al., 2018; Charlton and Vauquelin, 2010). In principal, the problem can be avoided by incorporating receptor binding kinetics of the agonist into the model equations. Such equations were derived for most of the models (Appendix A). Simulated data indicated a lag in the initial rise of response when agonist equilibrates slowly with receptor (Supplemental Fig. S2). This feature is potentially diagnostic of slow agonist equilibration. Slow equilibration can also be detected in agonist washout or blockade experiments (Hoare et al., 2018). These considerations notwithstanding, the quantification of agonist binding kinetics using the functional models here is likely to be challenging because the equations are highly parameterized.

In drug discovery, a simple application of the model is as follows. A biosensor is used to measure response over time. Biosensors enable real-time measurement of signaling using a single well/plate for all the time points, greatly improving the efficiency time course measurement (Lohse et al., 2012; Marullo and Bouvier, 2007). The time course is measured across a range of agonist concentrations, in a standard concentration-response format. The time course data for a maximally-stimulating concentration of agonist, maximally-stimulating at all time points, is then used for the kinetic model analysis. The data are fit to the appropriate equation. Let us assume this is a horizontal exponential fit, commonly encountered in second messenger assays. The fit parameters are the plateau and the rate constant. *k_τ_*, the efficacy parameter is then calculated by multiplying the plateau by the rate constant (Section 3.2.4.). The generic regulation rate constant, *k_REG_*, is taken as the rate constant. Next, a time point is selected late in the time course. The concentration response curve is obtained at this time point and fitted to determine the EC_50_ and the E_max_ at this time point. Finally, these parameters are incorporated into an SAR table, giving four columns; *k_τ_*, the efficacy value; *k_REG_*, the regulation parameter; and the conventional EC_50_ and E_max_ parameters measured at a single time point. The data from such a paradigm could be used as follows in a lead optimization project. As done conventionally, the EC_50_ at a single time point gives an estimate of potency, albeit one affected by receptor reserve and potentially affected by differences of efficacy. Efficacy can now be assessed using *k_τ_*. This provides a better estimate of efficacy than E_max_ from a concentration-response curve at a single time point because it is independent of receptor reserve. This potentially aids selection of compounds for in vivo testing, and rationalization of results from these studies. The regulation rate constant might also be useful translationally. Since it defines the rate of blunting of the response, it might contribute to defining the duration of efficacy in vivo, depending on the relevance of the regulation mechanisms in the signaling assay to the duration of the response in vivo.

In summary, this study reports a new pharmacological model for measuring drug parameters in conventional signaling assays that, for the first time, considers regulation of signaling. The model will facilitate the development of new therapeutics by enabling distinction of agonist efficacy from regulation in lead optimization and by improving the translation of pharmacological data from in vitro assays to in vivo efficacy.

## Appendix A: Derivation of equations

In this study, equations were derived that define the response over time for the kinetic models. The equations are of the analytic form, *y* = *f*(*t*), where *y* is response and *f*(*t*) is a function of time and pharmacological parameters. The analytic form is accommodated by curve-fitting software commonly used by pharmacologists. The model equations were derived as follows. First, the model mechanism was drawn out schematically in chemical terms (see Fig. 1 for general scheme). Next, differential equations describing the time-dependent processes were derived based on first principles, specifically the law of mass action for interactions between species (e.g. response precursor and receptor), and exponential decay for the decline of a single species over time (e.g. response degradation). Next, the differential equations were re-arranged to enable straightforward solution to the analytic form. In some cases, this step involved the Laplace transform method (Hoare, 2017; Mayersohn and Gibaldi, 1970). Finally, the analytic solution was obtained. The derivation assumes the *RA-E_P_* complex is sufficiently transient that it does not contribute appreciably to the mass balance of receptor or response precursor.

The equations were used to simulate model behavior and analyze experimental data (Section 3). In addition, a side-by-side comparison of *E vs t* data from the models is presented in Supplemental information. The following models have been described previously – 1,3,4 and 8 (Hoare et al., 2018).

### A.1. Unregulated response (Model 1)

In this model the response is not regulated – the receptor does not desensitize, the response does not degrade and there is no precursor depletion. This is Model 1 of the original kinetic model (Hoare et al., 2018). The model is represented by Scheme 1:

**Scheme 1.**
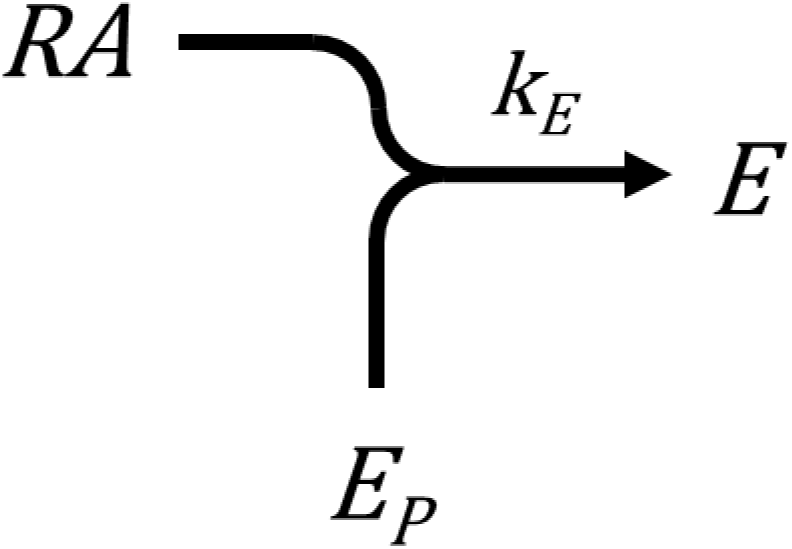

The *E vs t* equation for a rapidly-equilibrating agonist was derived in (Hoare et al., 2018), here named Eq. A.1:

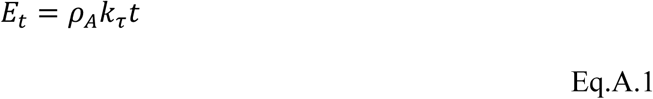

*k_τ_* is the agonist efficacy parameter, termed the transduction rate constant. It is defined as,

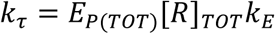

where *E_P_*_(*TOT*)_ is the total response precursor available in the system, [*R*]*_TOT_* the total concentration of receptors, and *k_E_* the response generation rate constant. *ρ*_*A*_ is fractional occupancy of receptor by agonist, defined as,

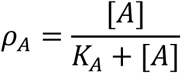

where [*A*] is the concentration of agonist and *K_A_* the agonist-receptor equilibrium dissociation constant.

At maximally-effective agonist concentrations, fractional occupancy is unity, i.e.,

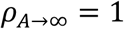

The *E vs t* equation for a saturating concentration of agonist is then (Eq. 1),

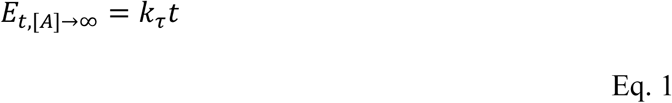

Note that this is the equation for a straight line intersecting the origin, where the gradient is *k_τ_*.

An equation has been derived that assumes agonist binding is not at equilibrium with the receptor [Eq. 9 of (Hoare et al., 2018)]. Here *k*_1_ is the agonist-receptor association rate constant and *k*_2_ the dissociation rate constant. The equation is,

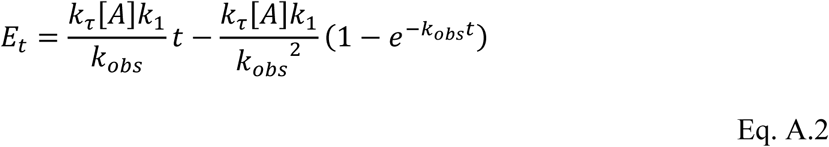

where *k_obs_* = [*A*]*k*_1_ + *k*_2_.

### A.2. Receptor desensitization (Model 2)

Receptor desensitization is incorporated into the model as a decline of the agonist-bound receptor concentration over time, shown in Scheme 2:

**Scheme 2.**
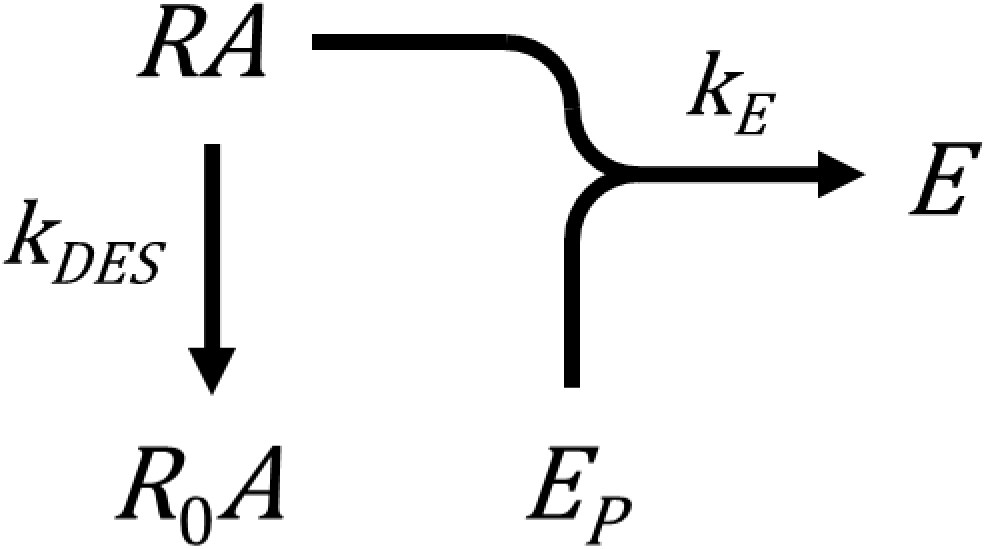

*RA* is the receptor species that can couple to the response precursor to generate the response, referred to as the active receptor. *R*_0_*A* is the desensitized receptor, which cannot couple to the response precursor and so is inactive with respect to signaling. *k_DES_* is the receptor desensitization rate constant. In this particular model, the response does not decay and the response precursor concentration remains constant over time, i.e. it is not depleted by generation of the response.

The differential equation for the response is,

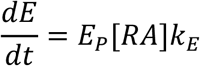

The goal is an equation in which *E* is the only time-dependent variable. This can be achieved using Laplace transforms to substitute the expression for [*RA*] into the expression for *E*. The Laplace transforms are,

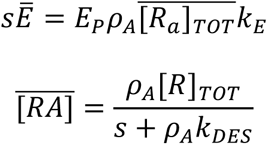

The transform for [*RA*] is Eq. B.4, derived in Appendix B. The transform for [*RA*] is now substituted into the transform for *E* and, after rearranging, we obtain:

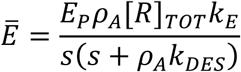

Since *E_p_* [*R*]*_TOT_k_E_* = *k*_*τ*_, this expression can be written as,

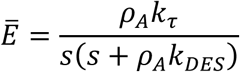

The *E vs t* equation is now obtained by taking the inverse Laplace transform, giving Eq. A.3:

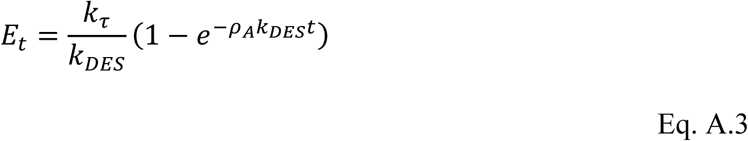

For the case of a saturating concentration of agonist, *ρ*_*A*_ is unity, and the *E vs t* equation reduces to Eq. 2:

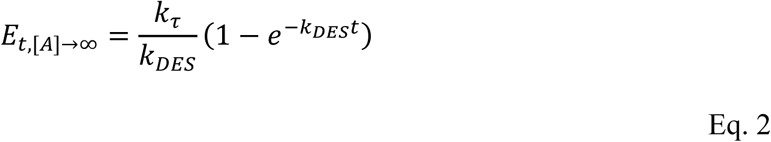

For the scenario in which agonist is not at equilibrium with the receptor, an equation can be derived that incorporates agonist-receptor binding kinetics. This is achieved here using by substituting the Laplace transform for [*RA*] into that for *E*. The transform for [*RA*] is Eq. B.6, derived in Appendix B:

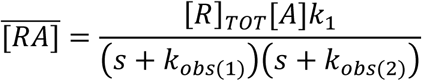

The terms *k_obs_*_(1)_ and *k_obs_*_(2)_ are defined in Appendix B. Substituting the transform for [*RA*] into the transform for *E* and rearranging yields,

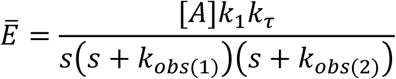

The *E vs t* equation is obtained by taking the inverse Laplace transform:

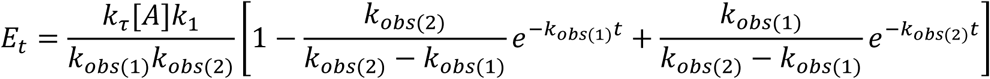

It can be shown that *k_obs_*_(1)_ *k_obs_*_(2)_ reduces to [*A*]*k_DES_*. Consequently, the *E vs t* equation reduces to the final form, Eq. A.4:

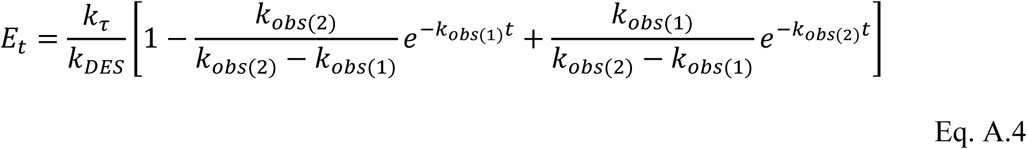

### A.3. Response degradation (Model 3)

In this model the response is degraded over time. There is no receptor desensitization or precursor depletion. This is Model 2 of the original kinetic model (Hoare et al., 2018). The model is represented by Scheme 3:

**Scheme 3.**
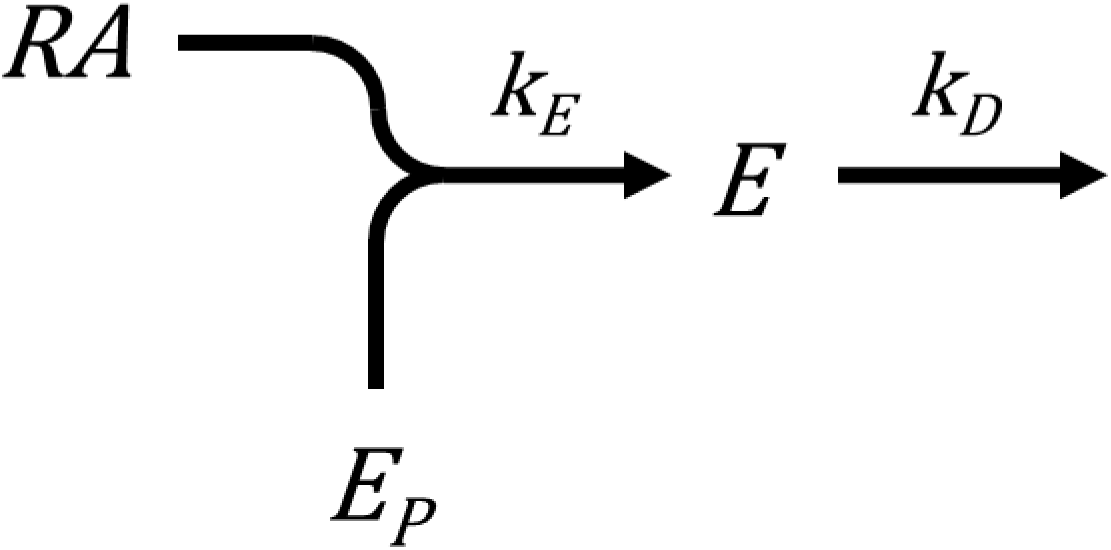

The *E vs t* equation for this mechanism has been derived previously (Hoare et al., 2018) and is given here as Eq. A.5:

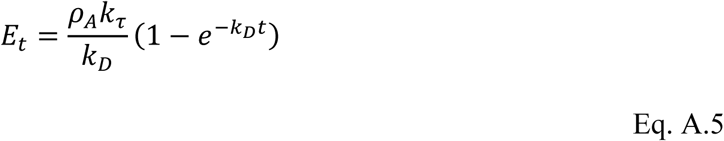

At a saturating concentration of agonist, the equation reduces to Eq. 3:

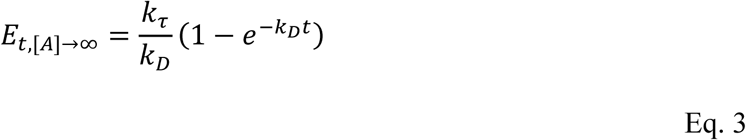

When agonist is not at equilibrium with the receptor, the equation, derived as Eq 10 in (Hoare et al., 2018), is:

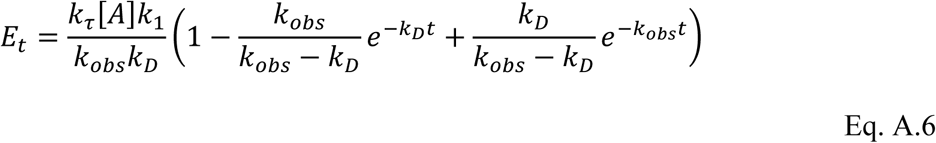

where *k_obs_* = [*A*]*k*_1_+*k*_2._

### A.4. Response degradation and recycling (Model 4)

In a variant of the response degradation model (Model 3), the response degradation process results in the reformation of the response precursor. In other words, the response recycles to the response precursor. This is Model 3 of the original kinetic model (Hoare et al., 2018) and is represented by Scheme 4:

**Scheme 4.**
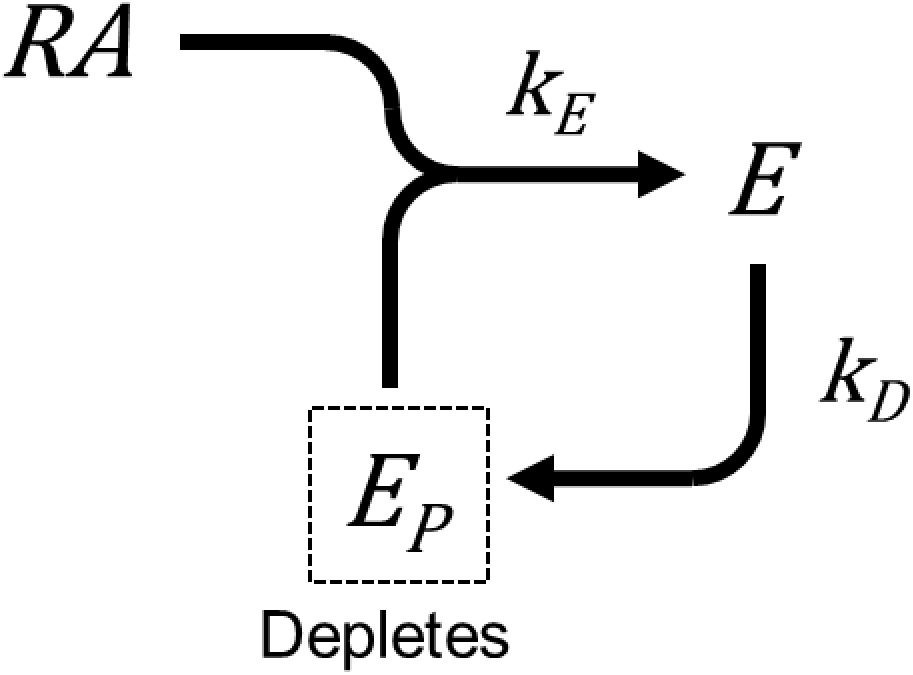

Note in this variant the response precursor becomes partially depleted. The *E vs t* equation here, Eq. A.7 below, is a rearranged form of the equation in the original study (Eq. 3 of (Hoare et al., 2018)). (Specifically *ρ*_*A*_ is introduced by substituting for [*A*]/(*K_A_*+[*A*]) in the original equation.)

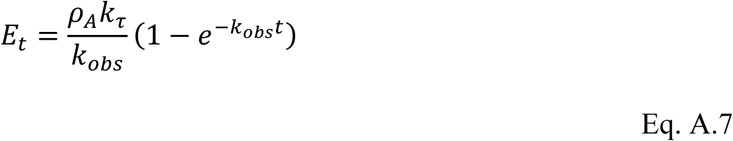

where,

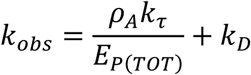

At a saturating concentration of agonist, the *E vs t* equation reduces to Eq. 4:

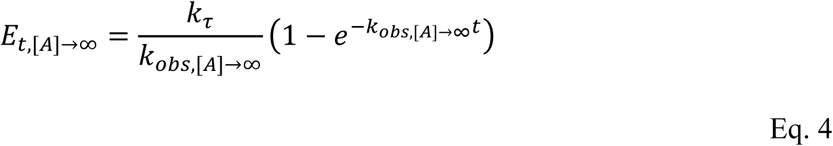

where,

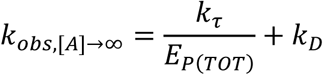

For the scenario in which agonist slowly equilibrates the receptor, there is no known analytical solution, as elaborated previously (Hoare et al., 2018). The *E vs t* data can be simulated by numerical solution of the differential equations for *E* and *E_P_*, and the analytical equation for [*RA*]. The equations for *E* and *E_P_* are,

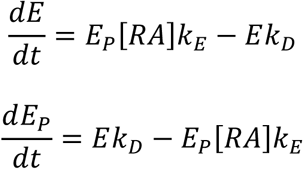

The analytical equation for [*RA*] is Eq. B.2, derived in Appendix B:

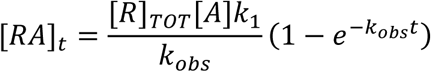

where *k_obs_* = [*A*]*k*_1_ + *k*_2_.

### A.5. Depletion of response precursor (Model 5)

In this model, the only response-limiting process is depletion of response precursor. There is no receptor desensitization or response degradation. The response precursor is depleted by its conversion to the response during the response generation process. This model is represented by Scheme 5:

**Scheme 5.**
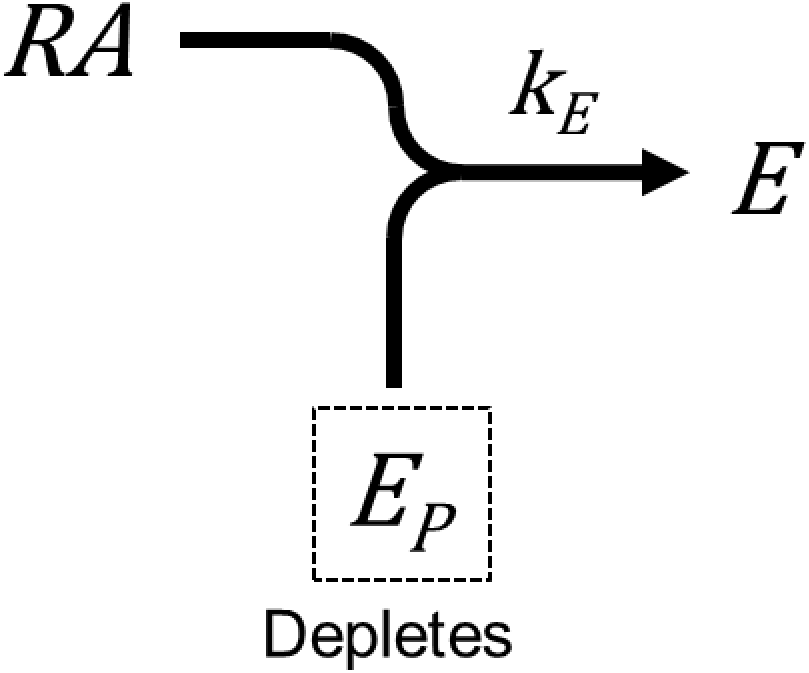

Two species are time-dependent in this model, *E* and *E_P_*. The derivation reduces this to the single time-dependent variable of interest, *E*. This is done using a conservation of mass equation for *E*. First the differential equation for *E* is written:

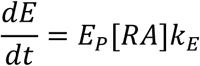

*E_P_* can be expressed as a function of the total amount of response material, *E_P_*_(*TOT*)_, which remains constant over time and is defined by the following conservation of mass equation:

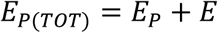

Solving for *E_P_* and substituting into the differential equation for *E* gives,

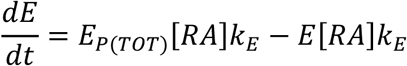

This equation can be integrated to obtain the *E vs t* equation

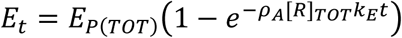

[*RA*]*k_E_* can be substituted with a function of *k_τ_*. Since *k_τ_* is defined as *E_P_*_(*TOT*)_[*RA*]*k_E_*, it is evident that,

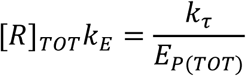

Substituting gives the final *E vs t* equation, Eq. A.8:

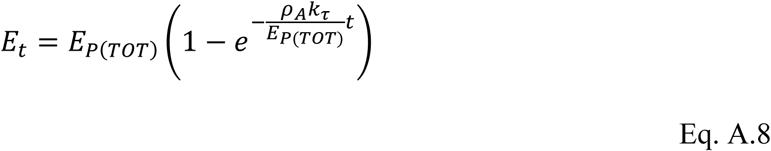

At saturating concentrations of agonist, the *E vs t* equation reduces to Eq. 5:

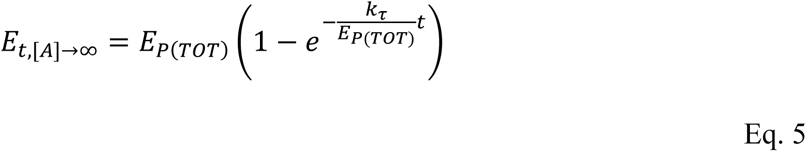

For the slow agonist equilibration scenario, we were able to derive an analytical solution. This involved handling a system of nonlinear differential equations. This was done using separation of variables. Because the ODE for [*RA*] decouples from that for *E* (Appendix B.1) we can substitute [*RA*] in the differential equation for *E* with the expression for [*RA*]*_t_* (Eq. B.2). After some re-arranging this gives:

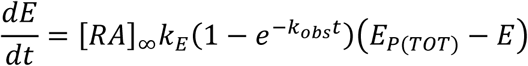

where [*RA*]_∞_ = [*R*]*_TOT_*[*A*]*k*_1_/*k_obs_*, introduced here for the purpose of clarity, and *k_obs_* = [*A*]*k*_1_ + *k*_2_. This equation can be solved by separation of variables as follows:

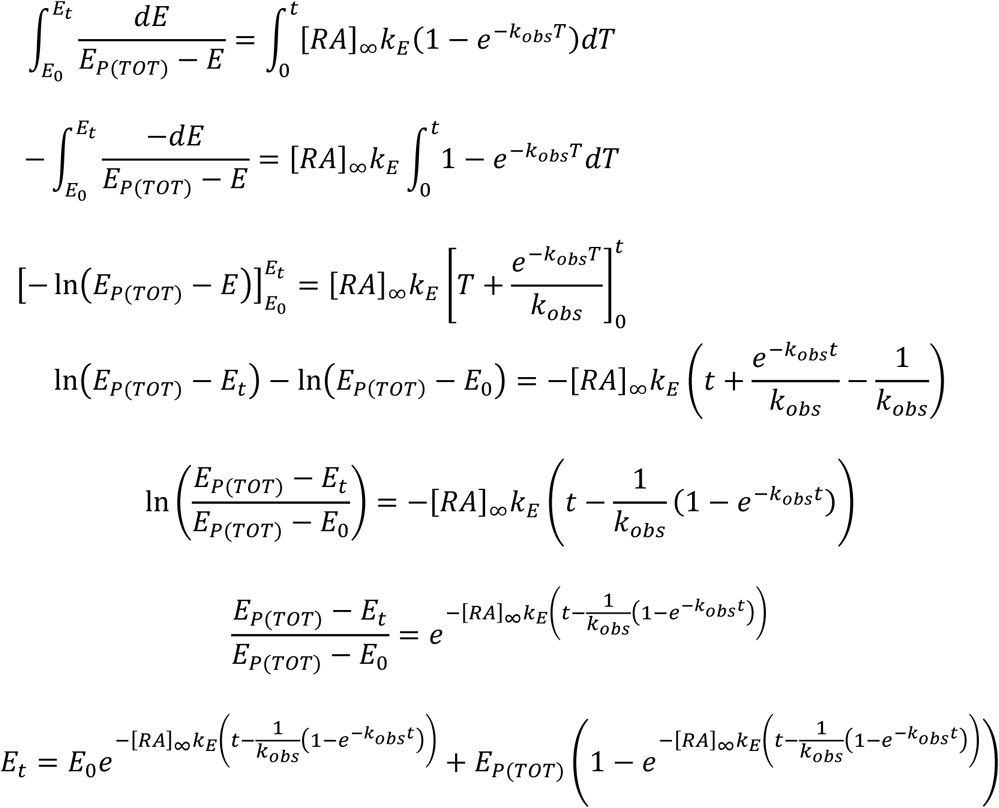

In the specific case when *EE*_0_ = 0, this simplifies to

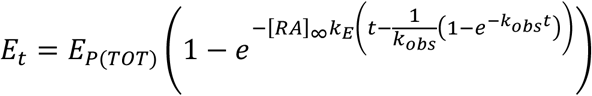

The [*RA*]_∞_*k*_*E*_ term can be rearranged to:

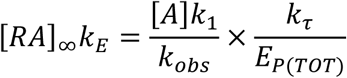

Substituting into the *E vs t* equation and rearranging gives the final equation, Eq. A.9:

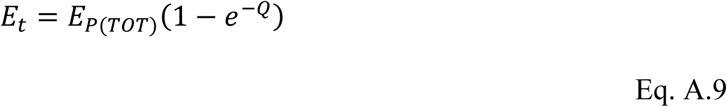

where,

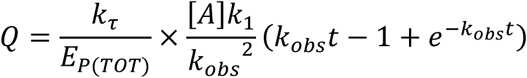

### A.6. Receptor desensitization and response degradation (Model 6)

In this model, the response is regulated by two processes; the receptor undergoes receptor desensitization, and the response is degraded. This model combines Models 2 and 3 and is represented by Scheme 6:

**Scheme 6.**
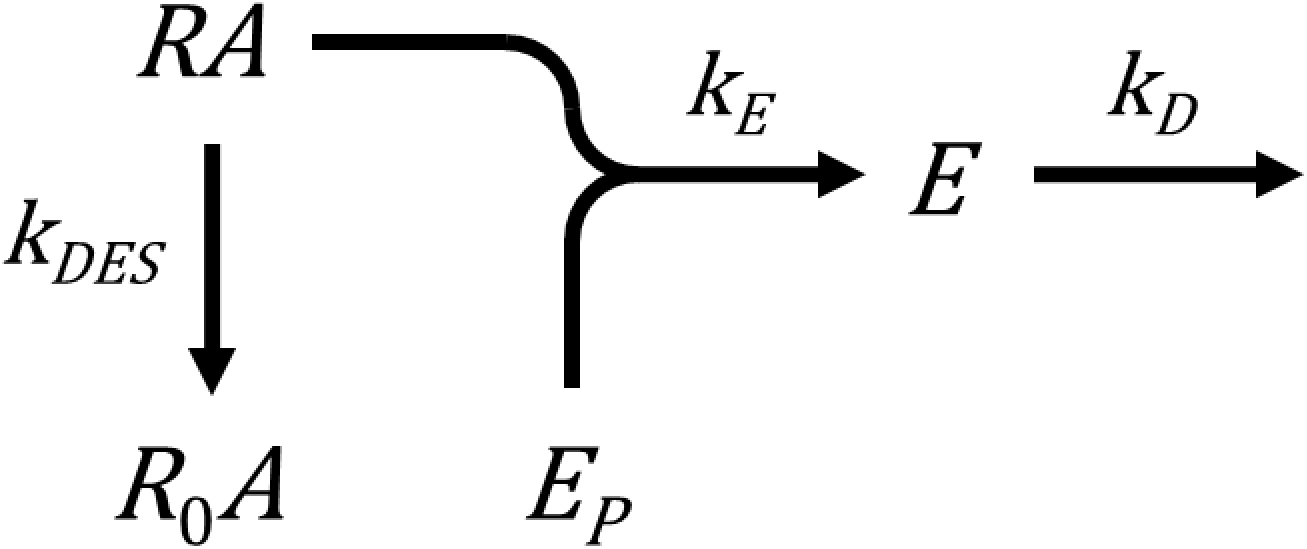

The *E vs t* equation is obtained as follows. The differential equation for *E* is,

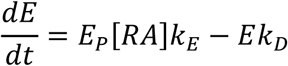

The derivation proceeds by taking the Laplace transform for [*RA*] (Eq. B.4) and substituting it into the transform for *E*, giving, after some rearranging,

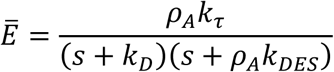

The *E vs t* equation is now obtained by taking the inverse Laplace transform, giving Eq. A.10:

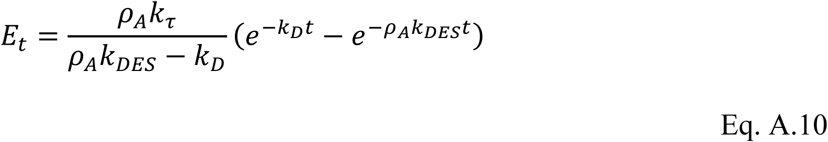

At saturating concentrations of agonist, the *E vs t* equation reduces to Eq. 6:

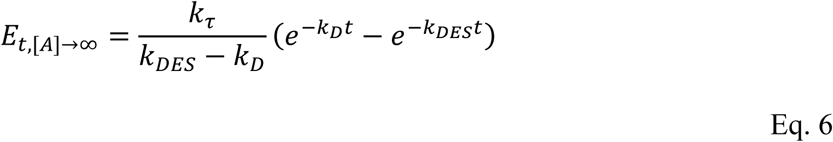

For the slow agonist equilibration scenario, the Laplace transform for [*RA*] is Eq. B.6. Substituting into the transform for *E* and rearranging gives,

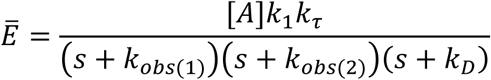

Taking the inverse transform gives the *E vs t* equation, Eq. A.11:

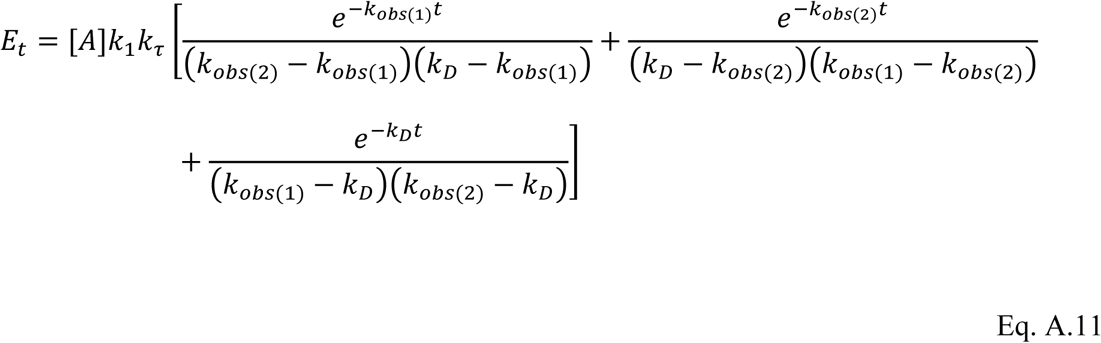

### A.7. Receptor desensitization and response degradation with recycling (Model 7)

In this variant of the receptor desensitization and response degradation model (Appendix A.6), the response degradation process results in the reformation of the response precursor. In other words, the response recycles to the response precursor. The model is represented by Scheme 7:

**Scheme 7.**
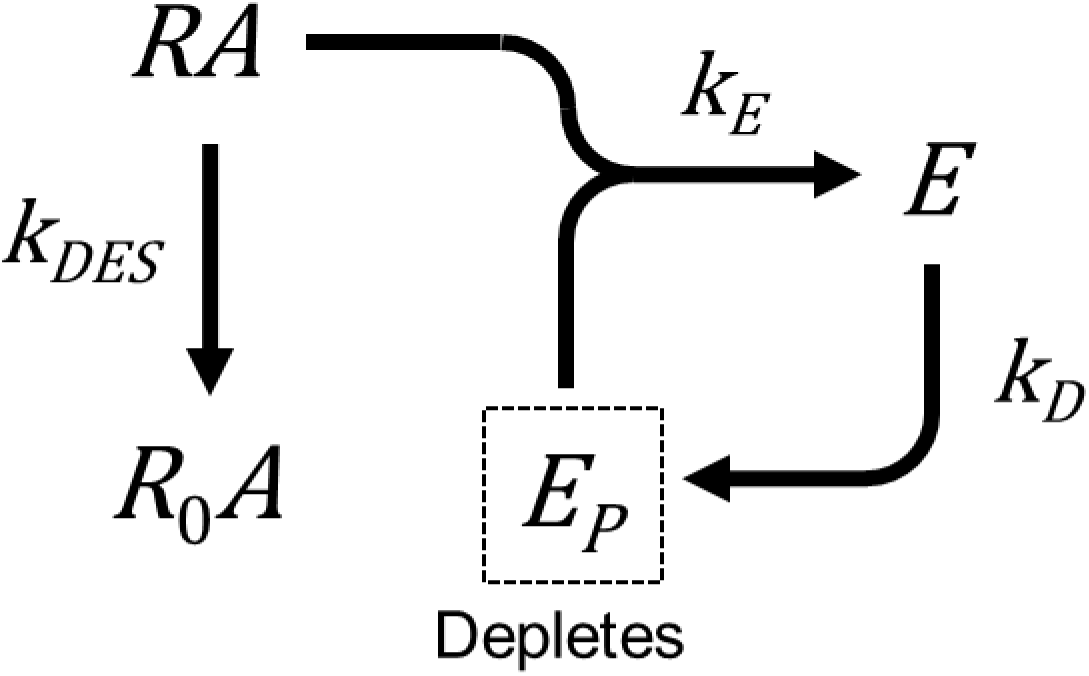

An analytical solution for *E* as a function of time could not be found for this system. The *E vs t* data can be simulated by numerical solution of the differential equations for *E* and *E_P_*. and the analytical solution for [*RA*]. The equations for *E* and *E_P_* are,

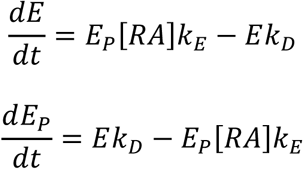

The analytical equation for [*RA*] is Eq. B.3 for the rapid equilibration scenario and Eq. B.5 for the slow equilibration model.

### A.8. Precursor depletion and response degradation (Model 8)

In this model, the precursor is depleted as it is being converted to the response and the response degrades. This is Model 4 of the original kinetic model (Hoare et al., 2018). The model is represented by Scheme 8:

**Scheme 8.**
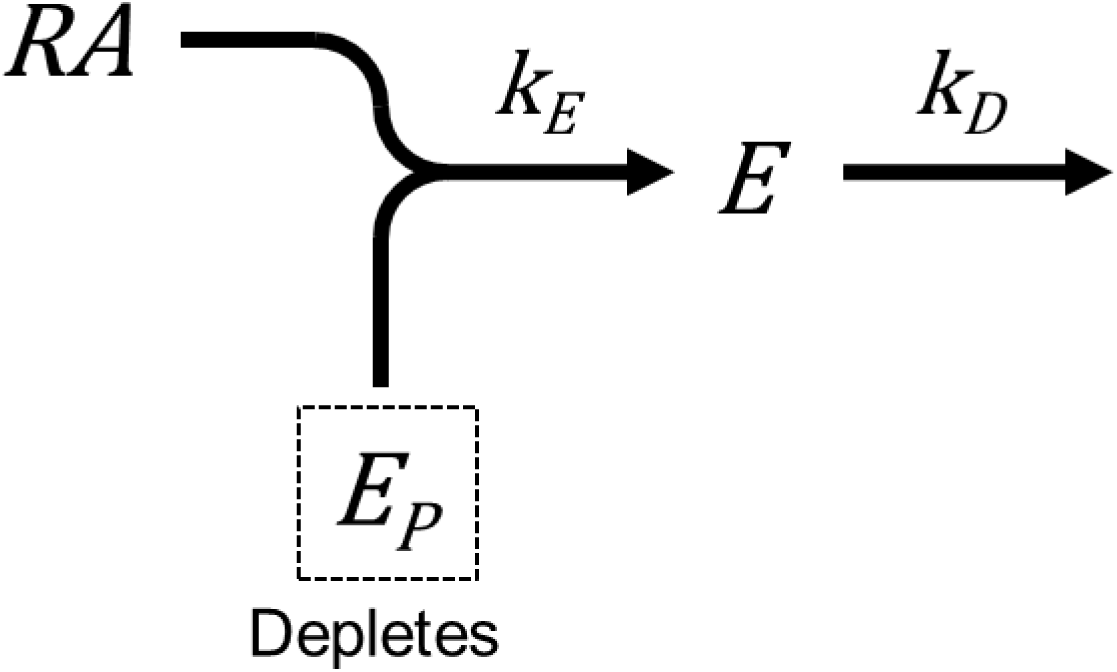

The equation used in this study is a modified form of Eq. 4 in (Hoare et al., 2018). A new, macroscopic term is introduced here called the depletion rate constant, *k_DEP_*. This term represents the rate at which the response precursor is depleted as it is being converted to the response. It is introduced as follows. The *E vs t* equation for the model is, as derived previously (Hoare et al., 2018),

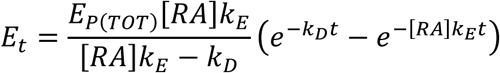

Substituting [*RA*] for *ρ_A_*[*R*]_TOT_ and introducing *k_τ_* gives,

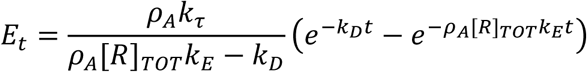

*k_DEP_* is defined here as the product of the receptor concentration and the response generation rate constant:

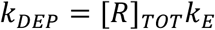

Substituting gives the *E vs t* equation (Eq. A.12):

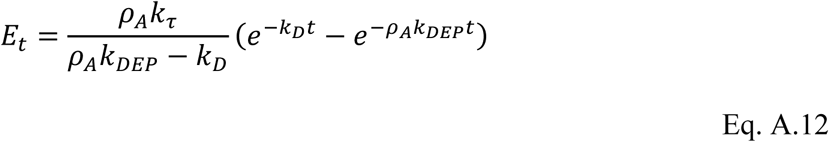

At saturating concentrations of agonist, the *E vs t* equation reduces to Eq. 8:

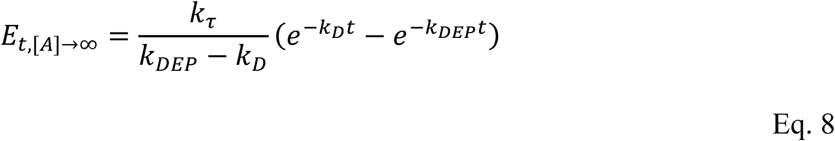

For the slow agonist equilibration scenario, an analytical solution for *E* could not be obtained, as described previously (Hoare et al., 2018). The *E vs t* data can be simulated by numerical solution of the differential equations for *E* and *E_P_*, and the analytical equation for [*RA*] (Eq. B.2). The differential equations are,

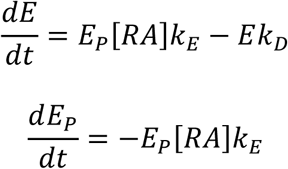

We have discovered an analytical solution is obtainable for response precursor, in other words the decline of *E_P_* which results from generation of *E*. Owing to decoupling of [*RA*] from *E* (Appendix B.1), the analytical equation for [*RA*] (Eq. B.2) can be substituted into the differential equation for *E_P_*:

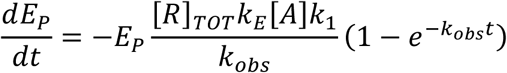

which can be rearranged to,

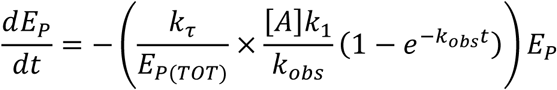

This ODE is related to the Gompertz growth/decay model (Kirkwood, 2015):

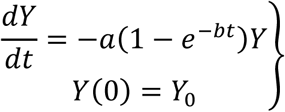

This problem has a known solution:

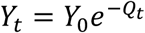

where,

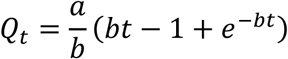

Applying to the initial value problem for *E_P_* gives the analytic solution for decline of the response precursor, Eq. A.13:

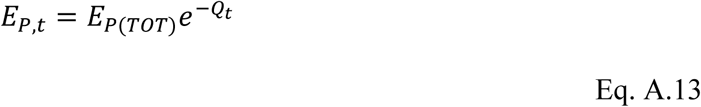

where,

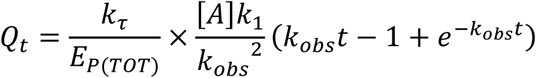

### A.9. Receptor desensitization and precursor depletion (Model 9)

**Scheme 9.**
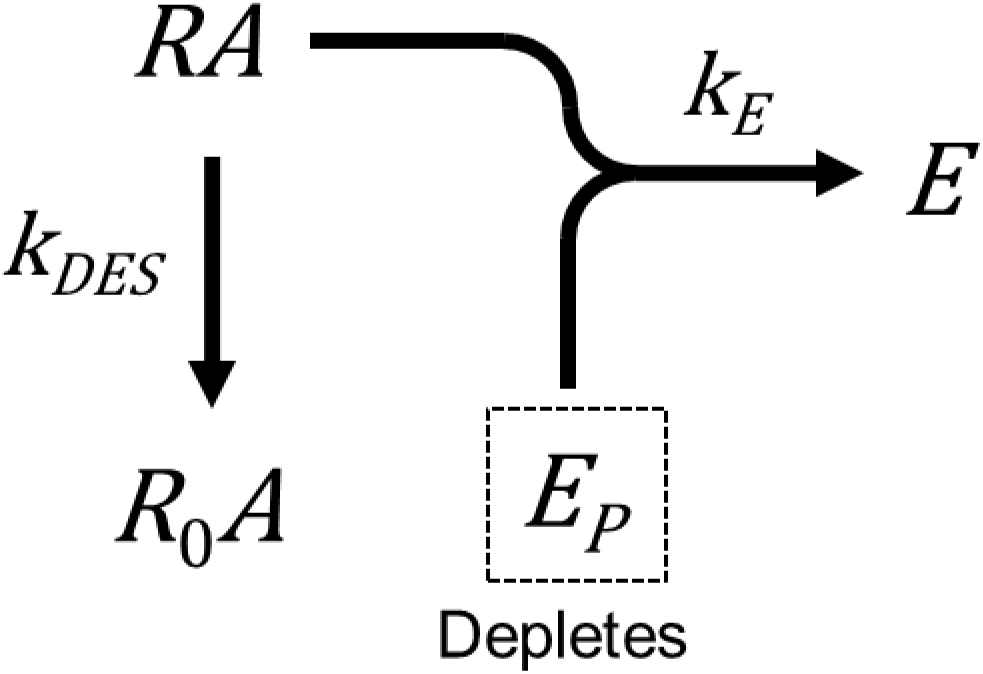

The *E vst t* equation is obtained by first deriving the equation for *E_P_* and then subtracting this from *E_P_*_(*TOT*)_ using the conservation of mass equation. The differential equation for *E_P_* is,

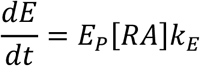

Owing to decoupling of the ODEs for [*RA*] and *E* (Appendix B.1), [*RA*]*_t_* can be represented by a time-dependent coefficient in the differential equations and *E_P_*. This is Eq. B.3. Substituting into the differential equation for *E_P_* gives,

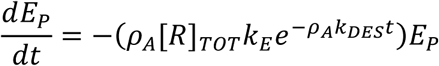

This expression is an initial value problem of the Gompertz growth/decay model type (Kirkwood, 2015):

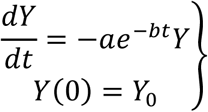

This problem has a known solution:

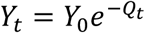

where,

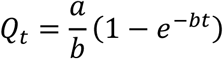

Applying to the expression for *E_P_* gives the analytic solution for decline of the response precursor (Eq. A.14):

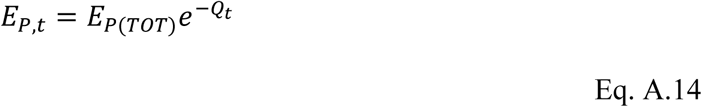

where,

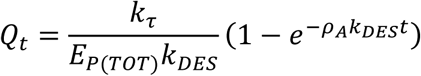

since [*R*]_TOT_*k*_*E*_ = *k*_*τ*_/*E*_*p(TOT)*_.

The *E vs t* equation, Eq. A.15, can now be determined from conservation of response precursor (*E_p(TOT)_* = *E_p_* + *E*):

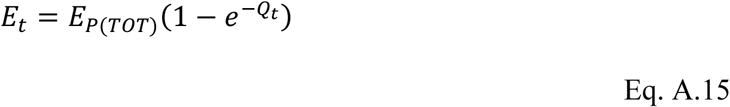

For the case of a saturating concentration of agonist, *ρ*_*A*_ is unity, and the *E vs t* equation reduces to Eq. 9:

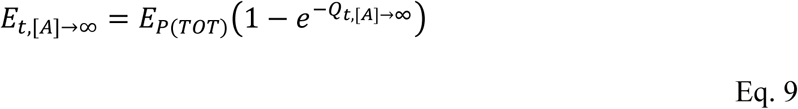

where,

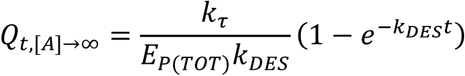

For the slow agonist equilibration scenario, we first derive an analytical equation for *E_P_*. The governing differential equations are,

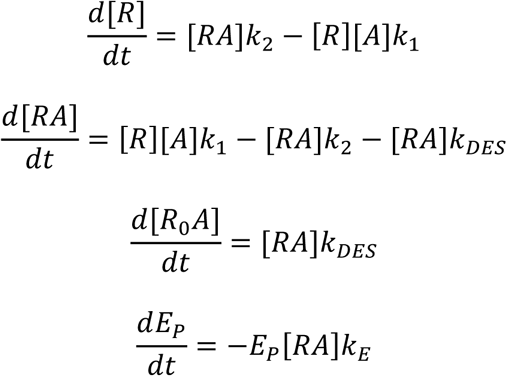

with initial conditions,

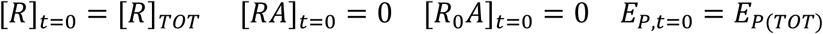

The system decouples (Appendix B.1.). The resulting system for [*R*] and [*RA*] is inhomogeneous, with a forcing term:

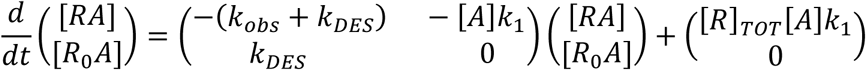

where *k_obs_*= [*A*]*k*_1_ + *k*_2_. The analytical solution for [*RA*]_*t*_ is Eq. B.5:

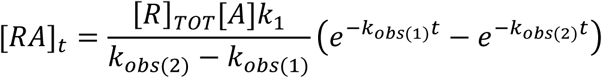

*k*_*obs*(1)_ and *k*_*obs*(2)_ are defined in Appendix B.2. We can now solve for *E_P_* by writing the differential equation for *E_P_* as a linear ODE:

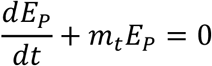

where,

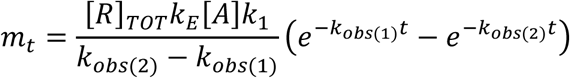

Solving with *E*_*p,t* =0_ = *E*_P(TOT)_, we find the *E_P_* vs t equation, Eq. A.16:

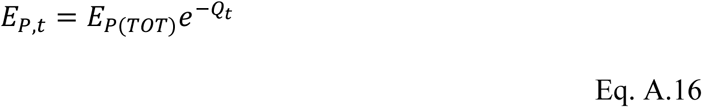

where,

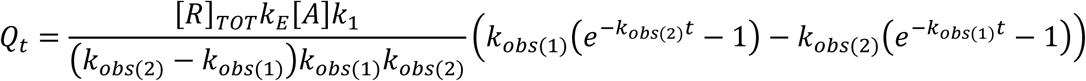

which can be rearranged to,

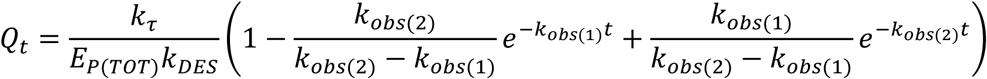

since *k*_obs(1)_*k_obs_*_(2)_ = [*A*]*k*_1_*k_DES_*.

The *E vs t* equation, Eq. A.17, is now obtained from the conservation of response precursor:

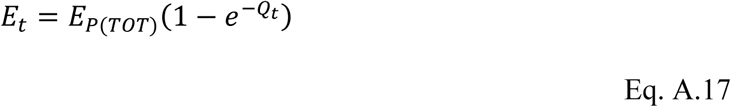

### A.10. Receptor desensitization, precursor depletion and response degradation (Model 10)

In this model, all three regulation mechanisms are in operation. The receptor desensitizes, the response precursor depletes as it is being converted to the response, and the response degrades. The model is represented by Scheme 10:

**Scheme 10.**
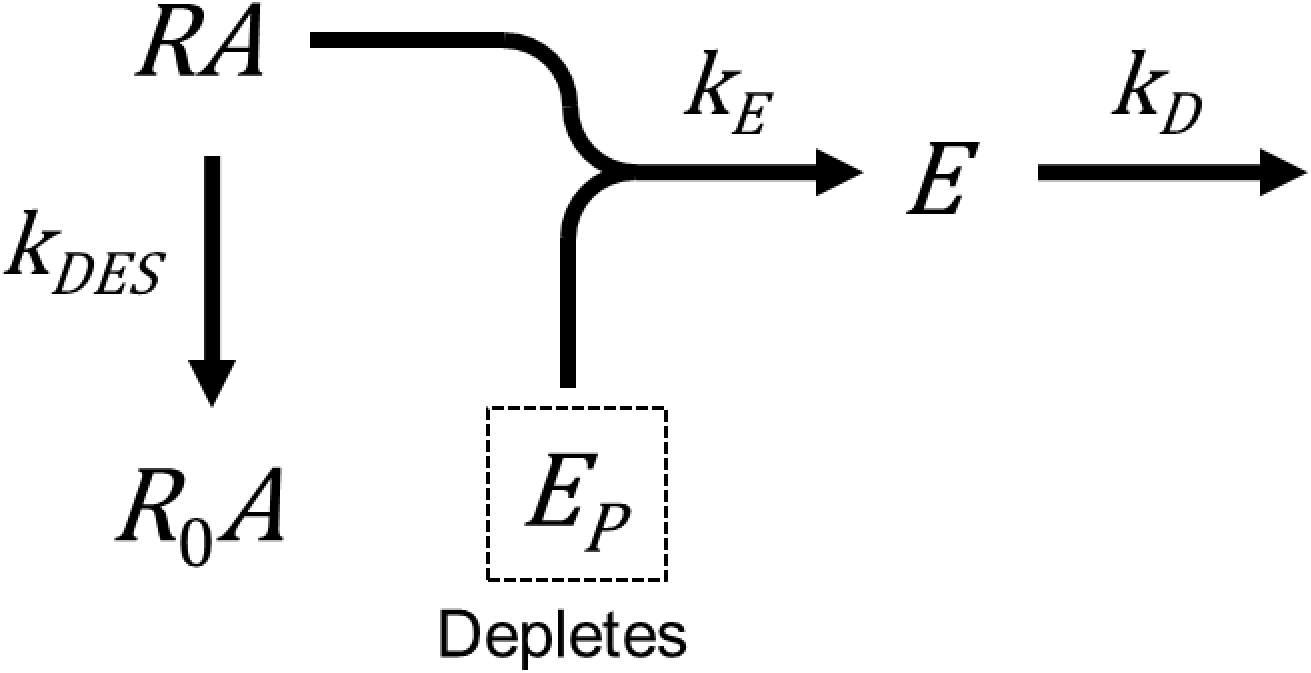

For the rapid agonist equilibration scenario, an analytical solution for *E* could not be obtained. (Hoare et al., 2018). The *E vs t* data can be simulated by numerical solution of the differential equations for *E* and *E_P_*, and the analytical solution for [*RA*]. The differential equations are:

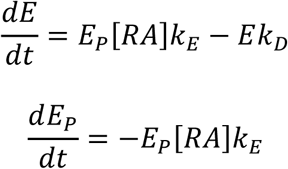

The analytical equation for [*RA*] is Eq. B.3 for the rapid equilibration scenario and Eq. B.5 for the slow equilibration model.

We were able to obtain a solution for response precursor, in other words the decline of *E_P_* which results from generation of *E*. These equations are the same as those for Model 9 (Appendix A.9), the receptor desensitization and precursor depletion model. (The additional step in Model 10 here compared with Model 9 is the degradation of *E*, which does not affect *E_P_* because there is no recycling.) For the rapid ligand equilibration scenario, the equation for *E_P_* is Eq. A.14 (and for the slow scenario, Eq. A.16 (Appendix A.9).

## Appendix B: Receptor-agonist binding equations

### Appendix B.1. Decoupling of [*RA*] and *E*

In all the models in this study the ODE for [*RA*] decouples from the ODE for *E* because there is no feedback from *E* to *RA*. In other words, the level of *E* does not affect the concentration of *RA*. This means that [*RA*]*_t_* can be considered a time-dependent coefficient in the differential equations for *E_P_*. This approach is applied to the slow agonist equilibration scenarios for Models 5 and 8, and both agonist binding scenarios for Models 9 and 10 (Appendix A.5, A.8, A.9 and A.10). In addition, if there is no response recycling, there is further decoupling of the ODEs for *E* and *E_P_*. This allows for the solution of *E_P_* in Models 8, 9 and 10 (Appendix A.8, A.9 and A.10).

### Appendix B.2. [*RA*] *vs t* equations

The response time course equations incorporate expressions for agonist-bound receptor species. Here these equations are derived. When agonist rapidly equilibrates with the receptor and there is no receptor desensitization, [*RA*] is given by the following equilibrium binding equation (Eq. B.1):

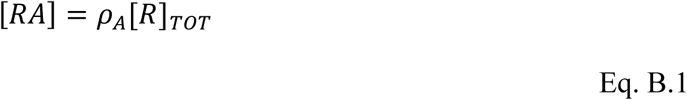

where *ρ_A_* is fractional occupancy of receptor by *A*:

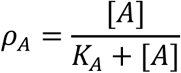

When agonist equilibrates slowly with the receptor and there is no receptor desensitization the [*RA*] *vs t* equation is the standard mass-action kinetic equation (Eq. B.2):

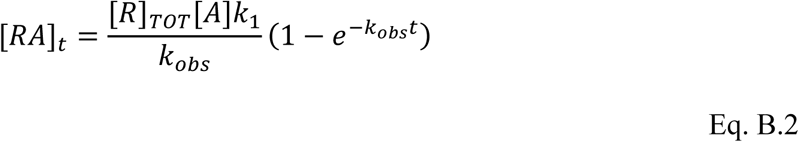

where *k_obs_* = [*A*]*k*_1_ + *k*_2_

When agonist equilibrates rapidly and the receptor desensitizes the [*RA*] *vs t* equation is derived as follows. Assuming agonist equilibrates rapidly with the receptor, [*RA*] can be expressed as a fraction of non-desensitized receptors occupied by agonist, as follows:

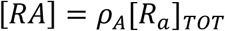

where [*R_a_*]*_TOT_* is the total concentration of non-desensitized receptors, i.e. [*R*]+[*RA*]. [*R_a_*]*_TOT_* is a time-dependent term because the receptor desensitizes. It is assumed the receptor desensitizes only if the agonist is bound to it. Consequently, the rate of desensitization of the receptor population is proportional to the fraction of non-desensitized receptors bound by agonist, i.e. *ρ_A_*. Under these conditions, the differential equation for the active receptor population is,

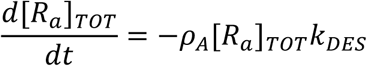

From this the differential equation for [*RA*] can be obtained:

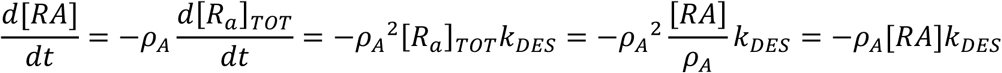

Integrating gives the [*RA*] *vs t* equation, Eq. B.3:

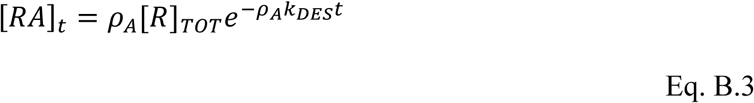

The Laplace transform is used in the derivation for Model 2 (Appendix B.2):

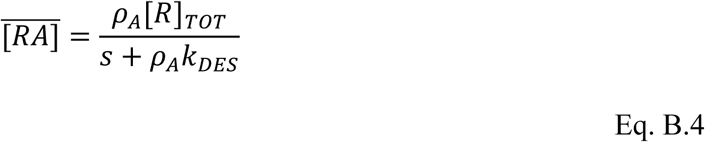

When agonist equilibrates slowly and the receptor desensitizes the [*RA*] *vs t* equation is derived as follows. The differential equations are,

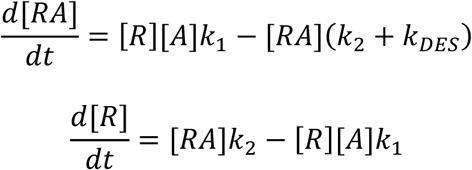

The derivation proceeds by taking the Laplace transform for [*R*] and substituting it into the transform for [*RA*]:

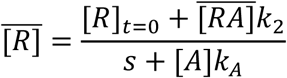

[*R*]*t*=0 is equal to [*R*]*_TOT_*. Hence,

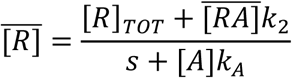

The transform for [*RA*] is,

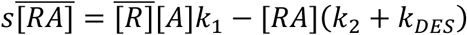

Substituting,

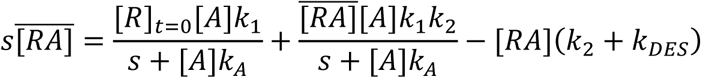

Solving for 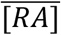 the procedures described in (Hoare, 2017) and taking the inverse transform gives the kinetic equation for [*RA*], Eq. B.4:

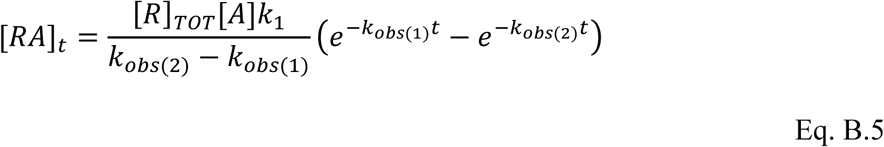

where

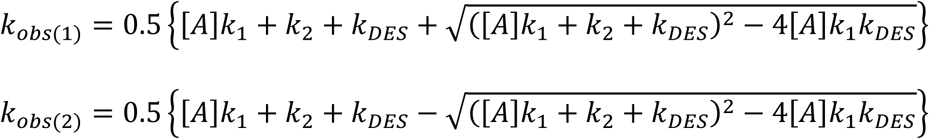

The Laplace transform is used in the derivation for Model 2 (Appendix B.2):

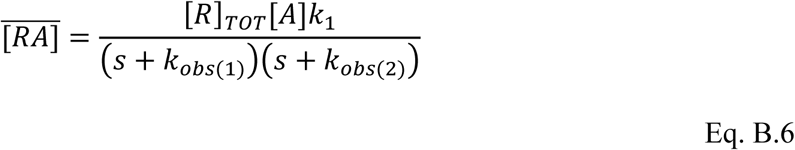

## Funding

This research did not receive any specific grant from funding agencies in the public, commercial, or not-for-profit sectors.

## Supplemental information

### Effect of multiple agonist concentration, equilibrium agonist binding assumption

In Figure S1, response to multiple agonist concentrations is presented for each of the models, assuming agonist rapidly equilibrates with the receptor.

### Model 1: Unregulated response

The gradient of the straight line is dependent on [*A*] (Fig. S1A). A plot of gradient vs [*A*] gives a hyperbola with [*A*]_50_ equal to *K_A_* and asymptote equal to *k_τ_*. EC_50_ does not change over time. For an example, see Fig 1 of (Rodbell et al., 1974).

### Model 2: Receptor desensitization

The *t*_1/2_ of the horizontal exponential curve is dependent on [*A*] (decreasing as [*A*] increases) whereas the plateau is the same for all [*A*] (Fig. S1B). As a result, EC_50_ decreases over time.

### Model 3: Response degradation

The *t*_1/2_ of the horizontal exponential curve is the same for all [*A*] whereas the plateau is dependent on [*A*] (Fig. S1C). Consequently, EC_50_ is constant over time. See Fig. 10 of (Hoare et al., 2018) for an example.

### Model 4: Response degradation and recycling

Both the *t*_1/2_ and plateau are dependent on agonist concentration (Fig. S1D). EC_50_ decreases over time. See Table 2 of (Traynor et al., 2002) for an example.

### Model 5: Precursor depletion

The *t*_1/2_ of the horizontal exponential curve is dependent on [*A*] (decreasing as [*A*] increases) whereas the plateau is the same for all [*A*] (Fig. S1E). As a result, EC_50_ decreases over time.

### Model 6: Receptor desensitization and response degradation

In this rise-and-fall exponential curve, the gradient of the rise, and the peak value, are dependent on [*A*] (Fig. S1F). Response declines to zero.

### Model 7: Receptor desensitization and response degradation with recycling

A distorted rise-and-fall curve is evident in which the decline phase is delayed, resulting in a bulge on the right-hand, downward part of the curve. The gradient of the rise, and the peak value, are dependent on [*A*] (Fig. S1G). Response declines to zero.

### Model 8: Precursor depletion and response degradation

In this rise-and-fall exponential curve, the gradient of the rise, and the peak value, are dependent on [*A*] (Fig. S1H). Response declines to zero. Note at later time points the complex relationship between response and [*A*]. Numerous examples are provided by cytoplasmic Ca^2+^ mobilization response data: Fig. 1 of (Princen et al., 2003), Fig. 4A of (Malysz et al., 2009), Fig. 5 of (Kassack et al., 2002), and Fig. 1 of (Milligan and Rees, 1999).

### Model 9: Receptor desensitization and precursor depletion

The response is described by a Gompertz curve, which resembles the horizontal exponential curve but it ascends more slowly at the later time points (see Fig. 9B). The initial gradient is dependent on [*A*] whereas the plateau is the same for all [*A*] (Fig. S1I).

### Model 10: Receptor desensitization, precursor depletion and response degradation

The response closely approaches a rise-and-fall exponential curve (see Section 3.4.3 for details). The gradient of the rise, and the peak value, are dependent on [*A*] (Fig. S1J). Response declines to zero.

**Fig. S1.**
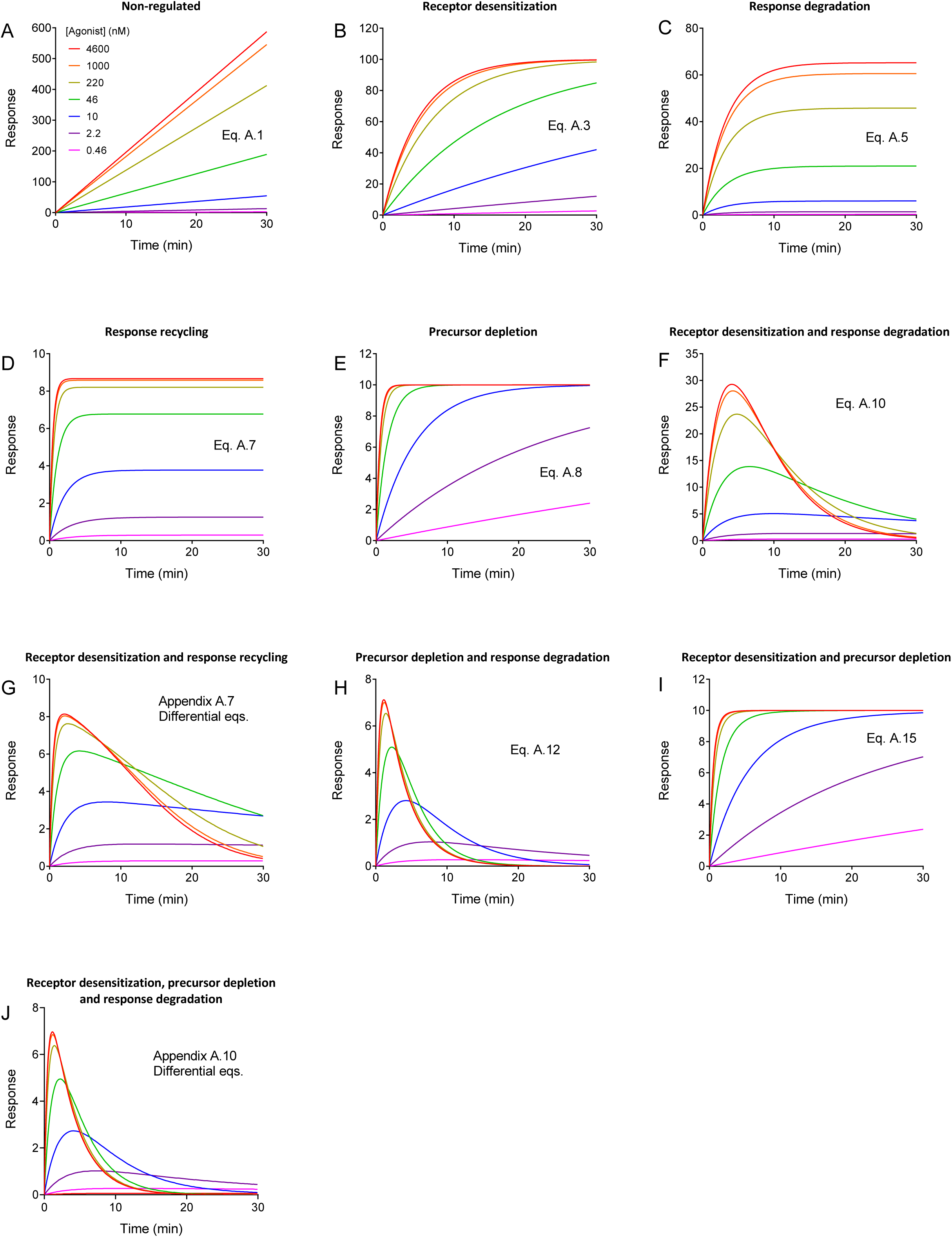
Simulated response time course data for response models (rapid agonist equilibration assumption). Data were simulated using the relevant equations indicated on the individual panels. The parameter values were: *K_A_*, 100 nM; *k*_τ_, 20 response units.min^−1^; *E_P_*_(*TOT*)_, 10 response units; *k_DES_*, 0.2 min^−1^; *k_D_*, 0.3 min^−1^.

**Effect of multiple agonist concentration, slow agonist equilibration assumption**

For these models, the curve shapes resemble the corresponding shape for the rapid equilibration but there is a clear lag in the rise phase of the curves (compare the corresponding panels of Figs. S1 and S2). The lag is dependent on the agonist concentration. The higher the concentration, the less evident the lag, a result of the increasing rate of agonist-receptor association due to mass action.

**Fig. S2.**
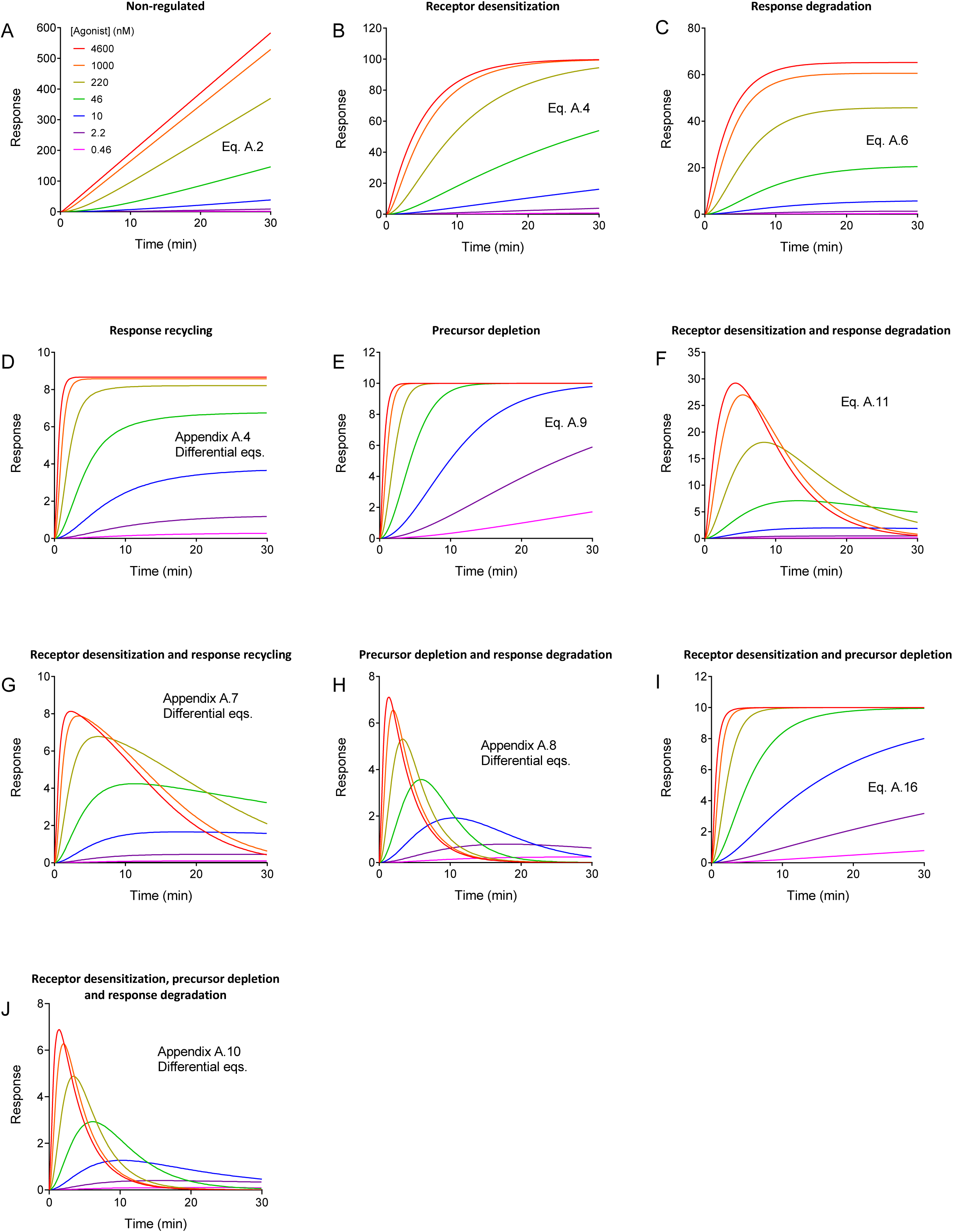
Simulated response time course data for response models (slow agonist equilibration assumption). Data were simulated using the relevant equation indicated on the individual panels. The parameter values were: *k*_1_, 10^6^ M^−1^min^−1^; *k*_2_, 0.1 min^−1^; *k*_τ_, 20 response units.min^−1^; *E_P_*_(*TOT*)_, 10 response units; *k_DES_*, 0.2 min^−1^; *k_D_*, 0.3 min^−1^.

**Depletion of response precursor models**

In some of the more complicated models, it was possible to obtain an analytical solution for *E_P_* even though it was not possible to obtain the solution for *E*. In some experimental paradigms it is possible to measure the decline of the precursor, for example the decline of phosphatidylinositol 4,5-bisphosphate (Tewson et al., 2013). Data for these models are simulated in Fig. S3: (A) Precursor depletion and response degradation, slow agonist equilibration (Model 8, Eq. A.13) (B) Receptor desensitization, precursor depletion and response degradation, rapid equilibration assumption (Model 10, Eq. 20). (C) Receptor desensitization, precursor depletion and response degradation, slow equilibration assumption (Model 10, Eq. A.14).

**Fig. S3.**
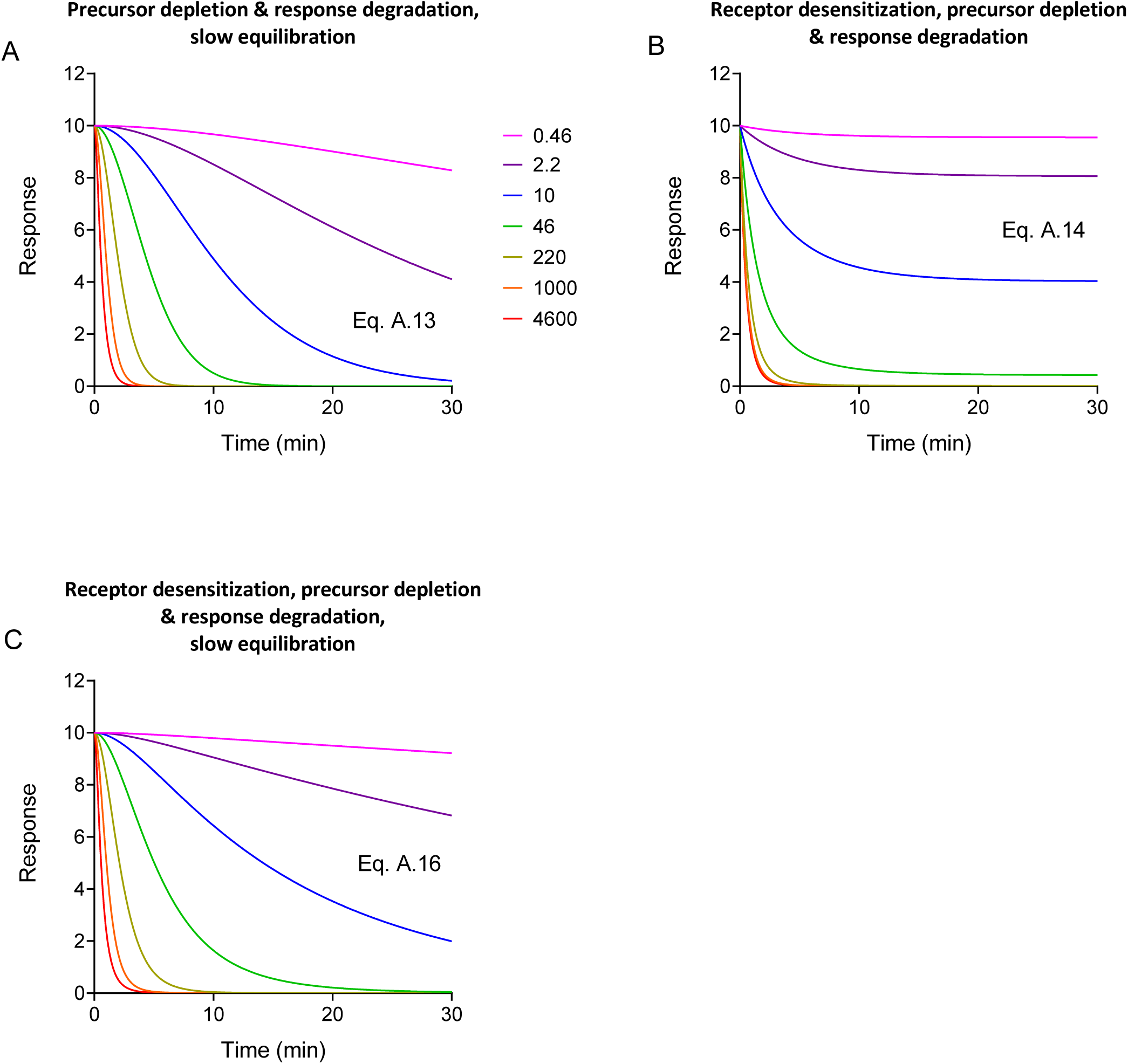
Simulated response time course data for response precursor. Models are those for which an analytical solution is not available for E but is available for EP. Data were simulated using the relevant equations indicated on the individual panels. The parameter values were: *k*_1_, 10^6^ M^−1^min^−1^; *k*_2_, 0.1 min^−1^; *k*_τ_, 20 response units.min^−1^; *E_P_*_(*TOT*)_, 10 response units; *k_DES_*, 0.2 min^−1^; *k_D_*, 0.3 min^−1^.

## References

Bdioui, S., Verdi, J., Pierre, N., Trinquet, E., Roux, T., Kenakin, T., 2018. Equilibrium Assays Are Required to Accurately Characterize the Activity Profiles of Drugs Modulating Gq-Protein-Coupled Receptors. Mol Pharmacol 94, 992–1006, doi:10.1124/mol.118.112573.

Berglund, M. M., Schober, D. A., Statnick, M. A., McDonald, P. H., Gehlert, D. R., 2003. The use of bioluminescence resonance energy transfer 2 to study neuropeptide Y receptor agonist-induced beta-arrestin 2 interaction. J Pharmacol Exp Ther 306, 147–56, doi:10.1124/jpet.103.051227.

Berridge, M. J., 1993. Inositol trisphosphate and calcium signalling. Nature 361, 315–25, doi:10.1038/361315a0.

Berridge, M. J., Bootman, M. D., Roderick, H. L., 2003. Calcium signalling: dynamics, homeostasis and remodelling. Nat Rev Mol Cell Biol 4, 517–29, doi:10.1038/nrm1155.

Black, J. W., Leff, P., 1983. Operational models of pharmacological agonism. Proc R Soc Lond B Biol Sci 220, 141–62.

Carafoli, E., 1991. Calcium pump of the plasma membrane. Physiol Rev 71, 129–53, doi:10.1152/physrev.1991.71.1.129.

Chang, Y. Y., 1968. Cyclic 3’,5’-adenosine monophosphate phosphodiesterase produced by the slime mold Dictyostelium discoideum. Science 161, 57–9.

Charlton, S. J., Vauquelin, G., 2010. Elusive equilibrium: the challenge of interpreting receptor pharmacology using calcium assays. Br J Pharmacol 161, 1250–65, doi:10.1111/j.1476-5381.2010.00863.x.

Colquhoun, D., 1998. Binding, gating, affinity and efficacy: the interpretation of structure-activity relationships for agonists and of the effects of mutating receptors. Br J Pharmacol 125, 924–47.

Davidson, J. S., Wakefield, I. K., Millar, R. P., 1994. Absence of rapid desensitization of the mouse gonadotropin-releasing hormone receptor. Biochem J 300 (Pt 2), 299–302.

Davis, J. S., West, L. A., Farese, R. V., 1986. Gonadotropin-releasing hormone (GnRH) rapidly stimulates the formation of inositol phosphates and diacyglycerol in rat granulosa cells: further evidence for the involvement of Ca2+ and protein kinase C in the action of GnRH. Endocrinology 118, 2561–71, doi:10.1210/endo-118-6-2561.

Eidne, K. A., Sellar, R. E., Couper, G., Anderson, L., Taylor, P. L., 1992. Molecular cloning and characterisation of the rat pituitary gonadotropin-releasing hormone (GnRH) receptor. Mol Cell Endocrinol 90, R5–9.

Ferguson, S. S., 2001. Evolving concepts in G protein-coupled receptor endocytosis: the role in receptor desensitization and signaling. Pharmacol Rev 53, 1–24.

Ferrandon, S., Feinstein, T. N., Castro, M., Wang, B., Bouley, R., Potts, J. T., Gardella, T. J., Vilardaga, J. P., 2009. Sustained cyclic AMP production by parathyroid hormone receptor endocytosis. Nat Chem Biol 5, 734–42, doi:10.1038/nchembio.206.

Fredriksson, R., Lagerstrom, M. C., Lundin, L. G., Schioth, H. B., 2003. The G-protein-coupled receptors in the human genome form five main families. Phylogenetic analysis, paralogon groups, and fingerprints. Mol Pharmacol 63, 1256–72, doi:10.1124/mol.63.6.1256.

Gilman, A. G., 1987. G proteins: transducers of receptor-generated signals. Annu Rev Biochem 56, 615–49, doi:10.1146/annurev.bi.56.070187.003151.

Gimenez, L. E., Baameur, F., Vayttaden, S. J., Clark, R. B., 2015. Salmeterol Efficacy and Bias in the Activation and Kinase-Mediated Desensitization of beta2-Adrenergic Receptors. Mol Pharmacol 87, 954–64, doi:10.1124/mol.114.096800.

Hausdorff, W. P., Caron, M. G., Lefkowitz, R. J., 1990. Turning off the signal: desensitization of beta-adrenergic receptor function. FASEB J 4, 2881–9.

Heding, A., Vrecl, M., Hanyaloglu, A. C., Sellar, R., Taylor, P. L., Eidne, K. A., 2000. The rat gonadotropin-releasing hormone receptor internalizes via a beta-arrestin-independent, but dynamin-dependent, pathway: addition of a carboxyl-terminal tail confers beta-arrestin dependency. Endocrinology 141, 299–306, doi:10.1210/endo.141.1.7269.

Heding, A., Vrecl, M., Bogerd, J., McGregor, A., Sellar, R., Taylor, P. L., Eidne, K. A., 1998. Gonadotropin-releasing hormone receptors with intracellular carboxyl-terminal tails undergo acute desensitization of total inositol phosphate production and exhibit accelerated internalization kinetics. J Biol Chem 273, 11472–7.

Hoare, S. R. J., 2017. Receptor binding kinetics equations: Derivation using the Laplace transform method. J Pharmacol Toxicol Methods, doi:10.1016/j.vascn.2017.08.004.

Hoare, S. R. J., Pierre, N., Moya, A. G., Larson, B., 2018. Kinetic operational models of agonism for G-protein-coupled receptors. J Theor Biol 446, 168–204, doi:10.1016/j.jtbi.2018.02.014.

Hofer, A. M., Landolfi, B., Debellis, L., Pozzan, T., Curci, S., 1998. Free [Ca2+] dynamics measured in agonist-sensitive stores of single living intact cells: a new look at the refilling process. EMBO J 17, 1986–95, doi:10.1093/emboj/17.7.1986.

Hoskin, P. J., Hanks, G. W., 1991. Opioid agonist-antagonist drugs in acute and chronic pain states. Drugs 41, 326–44.

Hothersall, J. D., Brown, A. J., Dale, I., Rawlins, P., 2016. Can residence time offer a useful strategy to target agonist drugs for sustained GPCR responses? Drug Discov Today 21, 90–6, doi:10.1016/j.drudis.2015.07.015.

Houslay, M. D., Schafer, P., Zhang, K. Y., 2005. Keynote review: phosphodiesterase-4 as a therapeutic target. Drug Discov Today 10, 1503–19, doi:10.1016/S1359-6446(05)03622-6.

Inglese, J., Freedman, N. J., Koch, W. J., Lefkowitz, R. J., 1993. Structure and mechanism of the G protein-coupled receptor kinases. J Biol Chem 268, 23735–8.

Kenakin, T., 2009a. “Hit” to drug: Lead optimization. A Pharmacology Primer: Theory, Applications and Methods. Academic Press, pp. 239–272.

Kenakin, T., 2009b. Quantifying biological activity in chemical terms: a pharmacology primer to describe drug effect. ACS Chem Biol 4, 249–60, doi:10.1021/cb800299s.

Kenakin, T., Watson, C., Muniz-Medina, V., Christopoulos, A., Novick, S., 2012. A simple method for quantifying functional selectivity and agonist bias. ACS Chem Neurosci 3, 193–203, doi:10.1021/cn200111m.

Kirkwood, T. B., 2015. Deciphering death: a commentary on Gompertz (1825) ‘On the nature of the function expressive of the law of human mortality, and on a new mode of determining the value of life contingencies’. Philos Trans R Soc Lond B Biol Sci 370, doi:10.1098/rstb.2014.0379.

Klein Herenbrink, C., Sykes, D. A., Donthamsetti, P., Canals, M., Coudrat, T., Shonberg, J., Scammells, P. J., Capuano, B., Sexton, P. M., Charlton, S. J., Javitch, J. A., Christopoulos, A., Lane, J. R., 2016. The role of kinetic context in apparent biased agonism at GPCRs. Nat Commun 7, 10842, doi:10.1038/ncomms10842.

Klueppelberg, U. G., Gates, L. K., Gorelick, F. S., Miller, L. J., 1991. Agonist-regulated phosphorylation of the pancreatic cholecystokinin receptor. J Biol Chem 266, 2403–8.

Kohout, T. A., Lin, F. S., Perry, S. J., Conner, D. A., Lefkowitz, R. J., 2001. beta-Arrestin 1 and 2 differentially regulate heptahelical receptor signaling and trafficking. Proc Natl Acad Sci U S A 98, 1601–6, doi:10.1073/pnas.041608198.

Krupnick, J. G., Benovic, J. L., 1998. The role of receptor kinases and arrestins in G protein-coupled receptor regulation. Annu Rev Pharmacol Toxicol 38, 289–319, doi:10.1146/annurev.pharmtox.38.1.289.

Kuzmic, P., 2009. DynaFit--a software package for enzymology. Methods Enzymol 467, 247–80, doi:10.1016/S0076-6879(09)67010-5.

Lane, J. R., May, L. T., Parton, R. G., Sexton, P. M., Christopoulos, A., 2017. A kinetic view of GPCR allostery and biased agonism. Nat Chem Biol 13, 929–937, doi:10.1038/nchembio.2431.

Leff, P., 1986. Potential errors in agonist dissociation constant estimation caused by desensitization. J Theor Biol 121, 221–32.

Lohse, M. J., Nuber, S., Hoffmann, C., 2012. Fluorescence/bioluminescence resonance energy transfer techniques to study G-protein-coupled receptor activation and signaling. Pharmacol Rev 64, 299–336, doi:10.1124/pr.110.004309.

Lohse, M. J., Benovic, J. L., Codina, J., Caron, M. G., Lefkowitz, R. J., 1990. beta-Arrestin: a protein that regulates beta-adrenergic receptor function. Science 248, 1547–50.

Marullo, S., Bouvier, M., 2007. Resonance energy transfer approaches in molecular pharmacology and beyond. Trends Pharmacol Sci 28, 362–5, doi:10.1016/j.tips.2007.06.007.

Mayersohn, M., Gibaldi, M., 1970. Mathematical methods in pharmacokinetics. I. Use of the Laplace transform for solving differential rate equations. Vol. 34, American Journal of Pharmaceutical Education, pp. 608–614.

Merelli, F., Stojilkovic, S. S., Iida, T., Krsmanovic, L. Z., Zheng, L., Mellon, P. L., Catt, K. J., 1992. Gonadotropin-releasing hormone-induced calcium signaling in clonal pituitary gonadotrophs. Endocrinology 131, 925–32, doi:10.1210/endo.131.2.1379169.

Merida, I., Avila-Flores, A., Merino, E., 2008. Diacylglycerol kinases: at the hub of cell signalling. Biochem J 409, 1–18, doi:10.1042/BJ20071040.

Miyawaki, A., Llopis, J., Heim, R., McCaffery, J. M., Adams, J. A., Ikura, M., Tsien, R. Y., 1997. Fluorescent indicators for Ca2+ based on green fluorescent proteins and calmodulin. Nature 388, 882–7, doi:10.1038/42264.

Motte, E., Le Stunff, C., Briet, C., Dumaz, N., Silve, C., 2017. Modulation of signaling through GPCR-cAMP-PKA pathways by PDE4 depends on stimulus intensity: Possible implications for the pathogenesis of acrodysostosis without hormone resistance. Mol Cell Endocrinol 442, 1–11, doi:10.1016/j.mce.2016.11.026.

Motulsky, H. J., 2019a. Equation: One phase association. GraphPad Curve Fitting Guide. Accessed 4 March 2019. https://www.graphpad.com/guides/prism/8/curve-fitting/index.htm?reg_exponential_association.htm.

Motulsky, H. J., 2019b. Equation: log(agonist) vs. response -- Variable slope. GraphPad Curve Fitting Guide. Accessed 4 March 2019. https://www.graphpad.com/guides/prism/8/curve-fitting/index.htm?reg_dr_stim_variable.htm.

Naccarato, W. F., Ray, R. E., Wells, W. W., 1974. Biosynthesis of myo-inositol in rat mammary gland. Isolation and properties of the enzymes. Arch Biochem Biophys 164, 194–201.

Navratilova, E., Waite, S., Stropova, D., Eaton, M. C., Alves, I. D., Hruby, V. J., Roeske, W. R., Yamamura, H. I., Varga, E. V., 2007. Quantitative evaluation of human delta opioid receptor desensitization using the operational model of drug action. Mol Pharmacol 71, 1416–26, doi:10.1124/mol.106.030023.

Paing, M. M., Stutts, A. B., Kohout, T. A., Lefkowitz, R. J., Trejo, J., 2002. beta-Arrestins regulate protease-activated receptor-1 desensitization but not internalization or Down-regulation. J Biol Chem 277, 1292–300, doi:10.1074/jbc.M109160200.

Pohl, S. L., Birnbaumer, L., Rodbell, M., 1969. Glucagon-sensitive adenyl cylase in plasma membrane of hepatic parenchymal cells. Science 164, 566–7.

Pohl, S. L., Birnbaumer, L., Rodbell, M., 1971. The glucagon-sensitive adenyl cyclase system in plasma membranes of rat liver. I. Properties. J Biol Chem 246, 1849–56.

Rang, H. P., 2006. The receptor concept: pharmacology’s big idea. Br J Pharmacol 147 Suppl 1, S9–16, doi:10.1038/sj.bjp.0706457.

Riccobene, T. A., Omann, G. M., Linderman, J. J., 1999. Modeling activation and desensitization of G-protein coupled receptors provides insight into ligand efficacy. J Theor Biol 200, 207–22, doi:10.1006/jtbi.1999.0988.

Rohatgi, A., 2018. WebPlotDigitizer. Web based tool to extract data from plots, images, and maps. Accessed 4 February 2018.

Schulz, D. W., Mailman, R. B., 1984. An improved, automated adenylate cyclase assay utilizing preparative HPLC: effects of phosphodiesterase inhibitors. J Neurochem 42, 764–74.

Slack, R. J., Hall, D. A., 2012. Development of operational models of receptor activation including constitutive receptor activity and their use to determine the efficacy of the chemokine CCL17 at the CC chemokine receptor CCR4. Br J Pharmacol 166, 1774–92, doi:10.1111/j.1476-5381.2012.01901.x.

Sriram, K., Insel, P. A., 2018. GPCRs as targets for approved drugs: How many targets and how many drugs? Mol Pharmacol, doi:10.1124/mol.117.111062.

Stadel, J. M., Nambi, P., Shorr, R. G., Sawyer, D. F., Caron, M. G., Lefkowitz, R. J., 1983. Catecholamine-induced desensitization of turkey erythrocyte adenylate cyclase is associated with phosphorylation of the beta-adrenergic receptor. Proc Natl Acad Sci U S A 80, 3173–7.

Streb, H., Heslop, J. P., Irvine, R. F., Schulz, I., Berridge, M. J., 1985. Relationship between secretagogue-induced Ca2+ release and inositol polyphosphate production in permeabilized pancreatic acinar cells. J Biol Chem 260, 7309–15.

Tay, D., Cremers, S., Bilezikian, J. P., 2018. Optimal dosing and delivery of parathyroid hormone and its analogues for osteoporosis and hypoparathyroidism - translating the pharmacology. Br J Clin Pharmacol 84, 252–267, doi:10.1111/bcp.13455.

Traynor, J. R., Nahorski, S. R., 1995. Modulation by mu-opioid agonists of guanosine-5’-O-(3-[35S]thio)triphosphate binding to membranes from human neuroblastoma SH-SY5Y cells. Mol Pharmacol 47, 848–54.

Traynor, J. R., Clark, M. J., Remmers, A. E., 2002. Relationship between rate and extent of G protein activation: comparison between full and partial opioid agonists. J Pharmacol Exp Ther 300, 157–61.

Tsien, R. W., Tsien, R. Y., 1990. Calcium channels, stores, and oscillations. Annu Rev Cell Biol 6, 715–60, doi:10.1146/annurev.cb.06.110190.003435.

Van der Graaf, P. H., Stam, W. B., 1999. Analysis of receptor inactivation experiments with the operational model of agonism yields correlated estimates of agonist affinity and efficacy. J Pharmacol Toxicol Methods 41, 117–25.

Violin, J. D., Dewire, S. M., Barnes, W. G., Lefkowitz, R. J., 2006. G protein-coupled receptor kinase and beta-arrestin-mediated desensitization of the angiotensin II type 1A receptor elucidated by diacylglycerol dynamics. J Biol Chem 281, 36411–9, doi:10.1074/jbc.M607956200.

Violin, J. D., DiPilato, L. M., Yildirim, N., Elston, T. C., Zhang, J., Lefkowitz, R. J., 2008. beta2-adrenergic receptor signaling and desensitization elucidated by quantitative modeling of real time cAMP dynamics. J Biol Chem 283, 2949–61, doi:10.1074/jbc.M707009200.

Xin, W., Tran, T. M., Richter, W., Clark, R. B., Rich, T. C., 2008. Roles of GRK and PDE4 activities in the regulation of beta2 adrenergic signaling. J Gen Physiol 131, 349–64, doi:10.1085/jgp.200709881.

Xu, C., Watras, J., Loew, L. M., 2003. Kinetic analysis of receptor-activated phosphoinositide turnover. J Cell Biol 161, 779–91, doi:10.1083/jcb.200301070.

Yu, R., Hinkle, P. M., 1997. Desensitization of thyrotropin-releasing hormone receptor-mediated responses involves multiple steps. J Biol Chem 272, 28301–7.

Yu, R., Hinkle, P. M., 2000. Rapid turnover of calcium in the endoplasmic reticulum during signaling. Studies with cameleon calcium indicators. J Biol Chem 275, 23648–53, doi:10.1074/jbc.M002684200.

Zernig, G., Issaevitch, T., Woods, J. H., 1996. Calculation of agonist efficacy, apparent affinity, and receptor population changes after administration of insurmountable antagonists: comparison of different analytical approaches. J Pharmacol Toxicol Methods 35, 223–37.

## References

Kassack, M. U., Hofgen, B., Lehmann, J., Eckstein, N., Quillan, J. M., Sadee, W., 2002. Functional screening of G protein-coupled receptors by measuring intracellular calcium with a fluorescence microplate reader. J Biomol Screen 7, 233–46, doi:10.1177/108705710200700307.

Malysz, J., Daza, A. V., Kage, K., Grayson, G. K., Yao, B. B., Meyer, M. D., Gopalakrishnan, M., 2009. Characterization of human cannabinoid CB2 receptor coupled to chimeric Galpha(qi5) and Galpha(qo5) proteins. Eur J Pharmacol 603, 12–21, doi:10.1016/j.ejphar.2008.11.047.

Milligan, G., Rees, S., 1999. Chimaeric G alpha proteins: their potential use in drug discovery. Trends Pharmacol Sci 20, 118–24.

Princen, K., Hatse, S., Vermeire, K., De Clercq, E., Schols, D., 2003. Evaluation of SDF-1/CXCR4-induced Ca2+ signaling by fluorometric imaging plate reader (FLIPR) and flow cytometry. Cytometry A 51, 35–45, doi:10.1002/cyto.a.10008.

Rodbell, M., Lin, M. C., Salomon, Y., 1974. Evidence for interdependent action of glucagon and nucleotides on the hepatic adenylate cyclase system. J Biol Chem 249, 59–65.

Tewson, P. H., Quinn, A. M., Hughes, T. E., 2013. A multiplexed fluorescent assay for independent second-messenger systems: decoding GPCR activation in living cells. J Biomol Screen 18, 797–806, doi:10.1177/1087057113485427.

